# Structure-aware protein function prediction at isoform resolution

**DOI:** 10.64898/2026.04.24.720502

**Authors:** Felicia T. Jiang, Runhao Zhao, Feng Liang, Xiang Zhao, Taoyong Cui, Zhenchao Tang, Yinghan Zhang, Danyu Chen, Minghao Xu, Yi Shuai, Tianli Luo, Xiangeng Wang, Chenchen Xu, Ziwei Wang, Weixin Zeng, Jia Xu, Wentao Zhang, Jiuyang Tang, Predrag Radivojac, Xin Wang

## Abstract

Understanding and accurately predicting function across protein isoforms has been a long-standing challenge with profound implications for both biological and translational research. However, most functional annotations and benchmarks remain tied to genes or reference protein sequences, leaving limited support for distinguishing protein isoforms from the same gene whose sequence, structure and domain composition may lead to different functions. To address this gap, we developed an isoform-centric framework for protein function and domain annotation that integrates a dense graph of sequence and structure similarity with protein features. We implemented this framework as 3DisoDeepPF, a graph-based multimodal model, and applied it to a breast cancer isoform atlas. Across canonical benchmarks and evaluations at isoform resolution, 3DisoDeepPF showed strong performance in predicting GO terms and Pfam domains and remained robust in tests with homology control. It further captured changes in Pfam domain composition among isoforms from the same gene, including reference-relative domain gain and loss. An evidence tracing module links predicted labels to supporting proteins in the graph, as illustrated by the calcium and integrin binding protein 1 (CIB1). Together, this study provides a framework informed by protein structure for function and domain annotation at isoform resolution, converting human protein isoform diversity into traceable functional evidence for cancer atlas interpretation.

---

Protein function prediction, Isoform, Proteoform, Gene Ontology, Pfam, Breast cancer Protein function provides the mechanistic link between molecular variation and biological phenotype, making functional annotation essential for genome interpretation, disease mechanism discovery and therapeutic target prioritization (Friedberg, 2006; Radivojac et al., 2013). As the expansion of omics data outpaces experimental characterization, computational annotation has become a complex yet critical challenge addressing the annotation gap increasingly visible in the post-AlphaFold era (Jumper et al., 2021; UniProt Consortium, 2025). Community efforts such as the Critical Assessment of Functional Annotation (CAFA), alongside numerous computational methods, have advanced protein function prediction (PFP) significantly (Jiang et al., 2016; Zhou et al., 2019). However, most existing approaches are developed and evaluated using gene-level annotations or reference protein sequences, rather than functions resolved for individual proteoforms. The lower resolution of this annotation framework impedes biomedical interpretation and prioritization (Smith et al., 2013).

To address this problem, we focus on alternative splicing (AS) derived protein isoforms as a biologically important and systematically analyzable entry point to proteoform-level function annotation. AS generates multiple transcripts from the same gene, often in a tissue-specific manner, yielding distinct protein sequences and structures (**Extended Data Fig. 1, Supplementary Note 1**) to substantially expand proteomic diversity (Graveley, 2001; Nilsen and Graveley, 2010; Liu et al., 2017). AS is pervasive in humans and consequential in disease (Graveley, 2001; Nilsen and Graveley, 2010; Weatheritt et al., 2016; Liu et al., 2017). It is estimated to generate roughly 100,000 human protein isoforms; however, only a minority have been experimentally characterized (Weatheritt et al., 2016; Liu et al., 2017; Mudge et al., 2025). For example, in a curated breast cancer isoform atlas, proxy analyses indicated that non-reference isoforms frequently differ from their predefined reference isoforms in Pfam domain architecture (**Methods**; **Extended Data Fig. 2c to 2e**; **Supplementary Note 3**) and inferred localization patterns (**Extended Data Fig. 2f to 2h**) (Mistry et al., 2021; Jiang et al., 2026). However, as summarized in **Supplementary Note 1**, most existing methods were developed for canonical proteins, typically transfer annotations by sequence similarity, and were trained with labels that do not distinguish isoforms, thus limiting their ability to account for subtle domain and functional differences between isoforms.

This unresolved gap highlights the need for isoform-specific PFP. Addressing this need, however, is non-trivial: available annotations are typically defined at the gene level, creating a mismatch with isoform-level prediction; closely related isoforms pose a substantial risk of homology leakage during model development and benchmarking; functional differences may arise from local sequence or structural remodeling rather than global changes, making informative signals sparse; and existing evaluation protocols are not designed to assess isoform-level functional differences. Together, these challenges motivate dedicated methods and evaluation frameworks for AS-derived isoform-level PFP.

To address these limitations, we present an isoform-centric PFP framework (Yu et al., 2020; Qiu et al., 2022) that simultaneously establishes evaluation principles for isoform-level analyses, including homology-aware and isoform-family-aware assessments **(Methods**). Methodologically, it embeds canonical proteins and isoforms in a dense sequence- and structure-similarity graph, integrating multimodal representations to capture subtle differences through informative graph links (**Methods, Supplementary Note 5**). We implemented this framework in 3DisoDeepPF (**3D iso**form **deep** learning for protein **f**unction **p**rediction), a graph attention model for Pfam domain and Gene Ontology (GO) term annotation over a protein similarity graph, designed for both canonical protein and isoform-level prediction (**Fig. 1a**). To support reproducibility and community reuse, we provide standardized knowledge bases for 76,804 canonical proteins (**Fig. 1b**) and 46,411 breast cancer isoforms (**Extended Data Fig. 2a,b**), together with an interactive web portal for prediction outputs (http://3disodeeppf.com/).

**Fig. 1.**
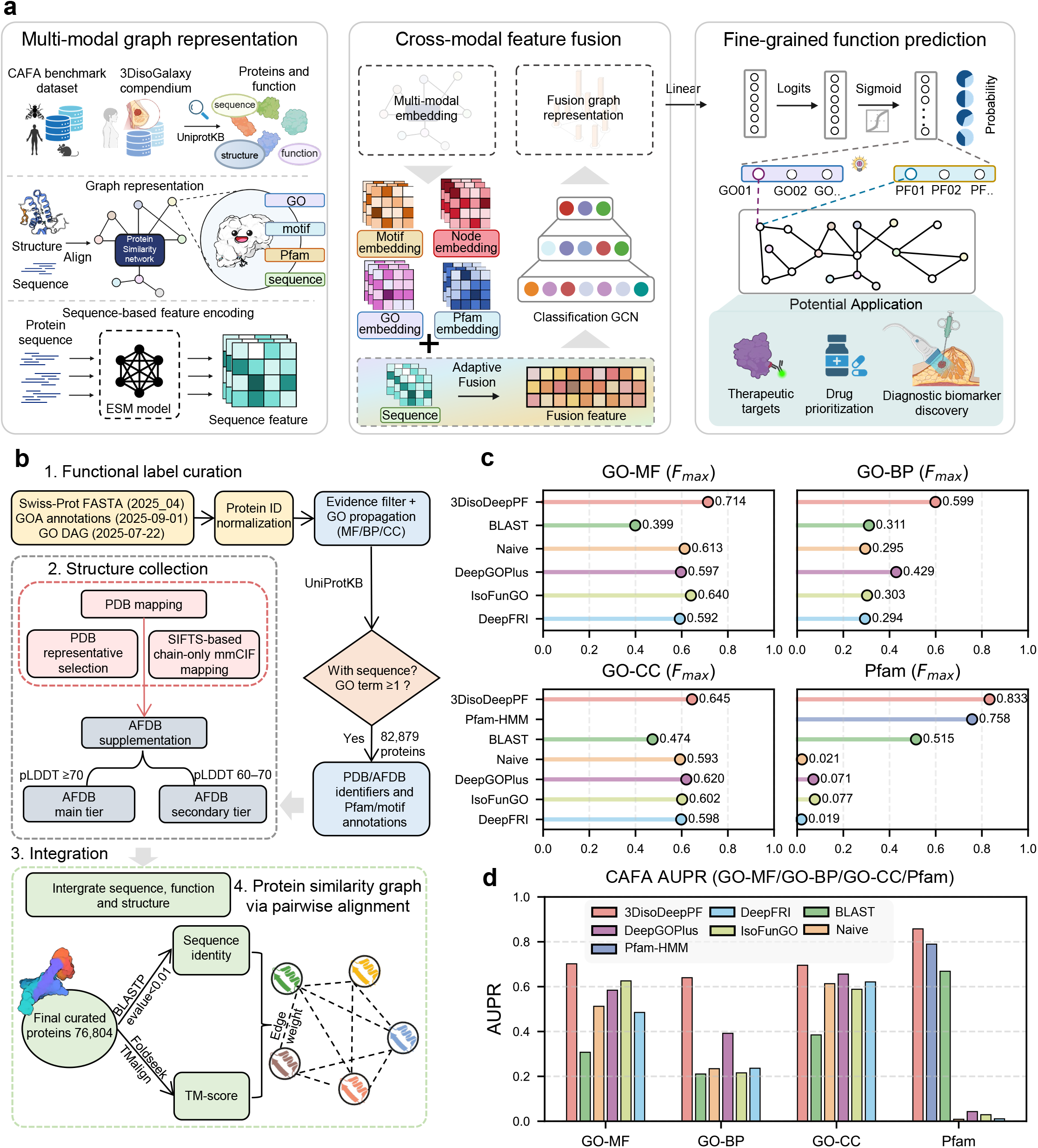
3DisoDeepPF framework, benchmark construction and performance. **a**, An overview of 3DisoDeepPF. Sequence and structure features are fused in a protein similarity graph to predict GO and Pfam annotations. **b**, Assembly of a CAFA-aligned, structure-supported benchmark from Swiss-Prot, the PDB and AlphaFoldDB, followed by graph construction using sequence identity and TM-score. **c**, Dot plots of protein-centric *F*_max_ scores across five bootstrap iterations, summarized across all test proteins and all GO and Pfam terms. Comparisons include CAFA baselines (BLAST and Naive), a structure-based deep learning method (DeepFRI), a sequence-based method (DeepGOPlus), and an isoform-oriented model (IsoFunGO). **d**, Bar plots of function-centric AUPR across methods for GO-MF, GO-BP, GO-CC and Pfam. Bars show means across five bootstrap iterations; error bars show s.d.

We first evaluated 3DisoDeepPF on canonical protein benchmarks to determine whether the framework remained competitive in standard annotation settings (**Methods, Supplementary Note 4**). 3DisoDeepPF consistently outperformed comparative methods in both threshold-based and ranking-based evaluations. It achieved the highest maximum F-score (*F*_max_) across GO molecular function (GO-MF), biological process (GO-BP), cellular component (GO-CC) and Pfam predictions (**Fig. 1c, Supplementary Note 2**). Consistent with this pattern, it yielded the best area under the precision-recall curve (AUPR) across all categories **(Fig. 1d**), with particularly strong gains in GO-BP and GO-CC, where annotation sparsity and class imbalance make prediction more challenging. These results show that 3DisoDeepPF remains competitive on standard canonical benchmarks, while the main analyses focus on isoform-level resolution.

We next evaluated 3DisoDeepPF at isoform resolution using a curated breast cancer isoform atlas (http://3disogalaxy.com/; **Extended Data Fig. 2a,b**; **Supplementary Note 4**) (Jiang et al., 2026). We visualized the retained sequence-structure similarity graph in a two-dimensional embedding and colored the nodes by curated Pfam annotations (**Supplementary Note 5**). Using the same graph construction and cutoff settings as the main model (**Methods** and **Supplementary Note 5**), local neighborhoods were often associated with the same Pfam domains (**Fig. 2a**) (Mistry et al., 2021), supporting the qualitative biological coherence of the integrated graph. Across AUPR and Fmax evaluations, 3DisoDeepPF performed strongly across Pfam and Gene Ontology prediction tasks, with the clearest gains in Pfam, GO-MF and GO-BP, and competitive GO-CC performance that was more apparent by *F*_max_ than by AUPR (**Fig. 2b,c**). To assess the contribution of the graph and multimodal components to isoform-level PFP, we performed component ablation in the breast cancer isoform-resolved setting. Removing the graph produced the largest performance drop, whereas sequence and structure provided substantial complementary gains, and Pfam or motif features added smaller but consistent improvements (**Extended Data Fig. 3**).

**Fig. 2.**
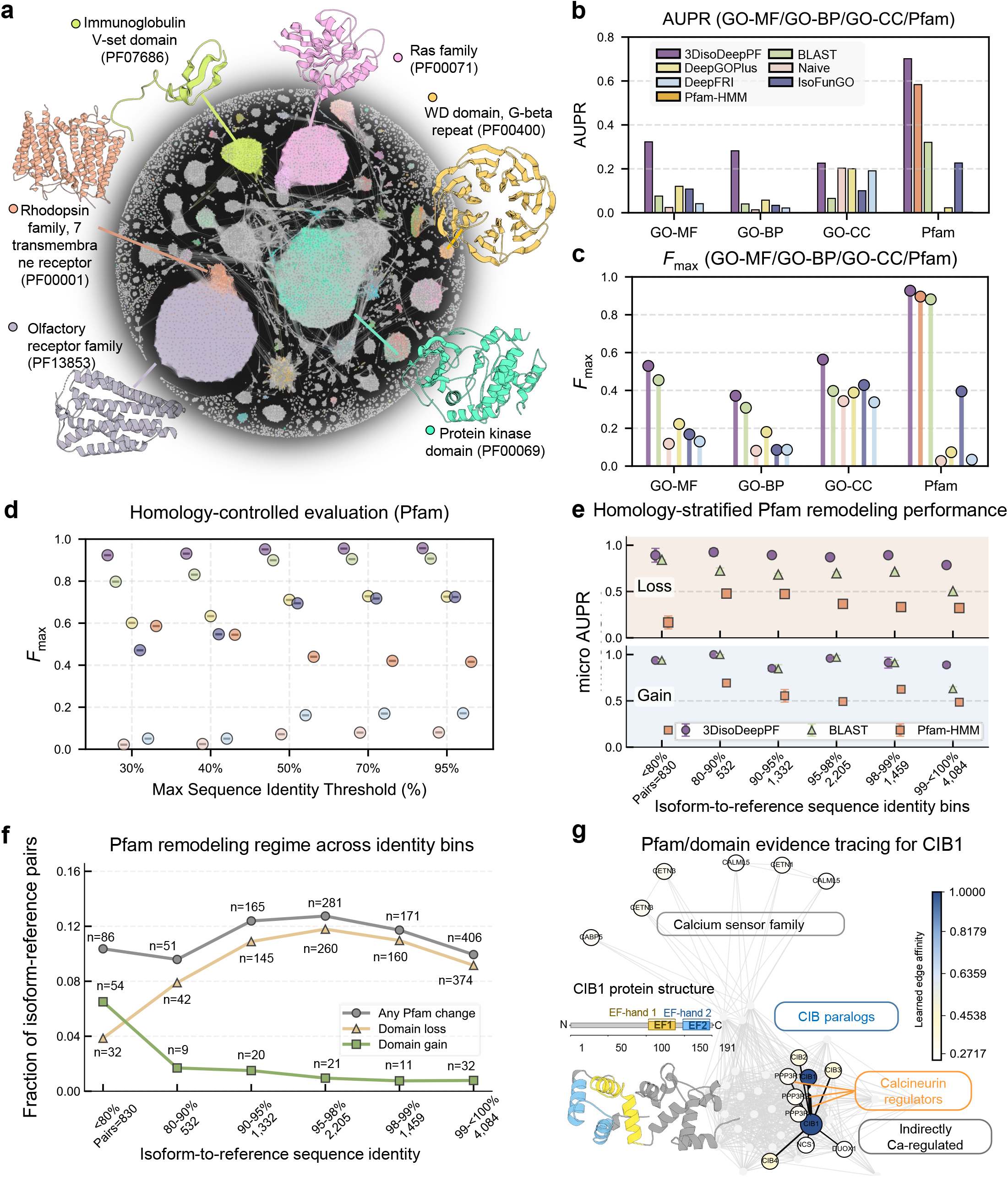
Isoform-resolved breast cancer landscape and isoform-specific evaluation of 3DisoDeepPF. **a**, Breast cancer protein isoform similarity graph. Nodes are colored by the 15 most frequent Pfam domains; isoforms without Pfam annotation are shown in grey. Six representative structures highlight selected domain-enriched regions with locally shared domain annotations. **b**,**c**, Function-centric AUPR (**b**) and protein-centric *F*_max_ (**c**) for GO-MF, GO-BP, GO-CC and Pfam across methods. In **b**, bars show means across five bootstrap iterations and error bars show s.d. In **c**, colored points denote mean *F*_max_ values across five bootstrap iterations. **d**, Pfam prediction under homology control across maximum allowed sequence-identity thresholds of 30–95%. **e**, Micro-AUPR for directional Pfam remodeling, evaluated separately for domain loss and gain relative to the reference isoform across isoform-to-reference sequence-identity bins. **f**, Fractions of isoform-to-reference pairs with any Pfam change, domain loss or domain gain across sequence-identity bins; numbers indicate pair counts. **g**, Domain-level evidence tracing for CIB1 (calcium and integrin-binding protein 1), showing predicted EF-hand annotations and the top 15 isoform neighbors supporting the predicted Pfam assignment; node color summarizes learned edge affinity to CIB1, with darker color indicating stronger model-inferred relatedness.

Because isoforms are often highly similar in sequence, we evaluated 3DisoDeepPF across a range of controlled settings to assess performance at different levels of similarity between training and test proteins. Under a controlled Pfam evaluation, 3DisoDeepPF remained strong across maximum sequence-identity cutoffs of 30-95% between training and test proteins (**Fig. 2d**), indicating that performance was broadly stable across all settings. GO-term predictions also remained competitive under the same evaluation scheme (**Extended Data Fig. 4a–c**), but the clearest gains were observed in the Pfam analyses, which directly assess changes in domain composition between isoforms.

We then defined a directional Pfam remodeling task across isoforms from the same gene, where Pfam remodeling denotes changes in domain composition relative to a predefined reference isoform and is quantified as domain gain or loss (**Methods**). Here, gain denotes domains present in an isoform but absent from the reference, whereas loss denotes domains present in the reference but absent from the isoform. 3DisoDeepPF achieved the strongest micro-AUPR for directional Pfam remodeling, demonstrating robust performance on both gain and loss predictions (**Extended Data Fig. 4d**). This advantage was maintained across bins defined by sequence identity to the reference isoform (**Fig. 2e**), indicating that the model captures remodeling signals across different levels of isoform divergence. The curated Pfam annotations showed that loss events were more frequent than gains across sequence-identity bins of isoform-reference pairs (**Fig. 2f**). This asymmetry may partly reflect the reference-based comparison, in which MANE or UniProtKB/Swiss-Prot canonical isoforms provide well-supported reference products against which shorter or locally altered isoforms are compared. Accordingly, loss-side performance is evaluated under denser annotation support, providing a more stable basis for performance assessment.

To examine the support for individual predictions, we applied an evidence tracing utility to the 3DisoDeepPF model (**Extended Data Fig. 5a**). Using a candidate therapeutic target CIB1 (calcium and integrin-binding protein 1) as an illustrative case (**Fig. 2g**), we focused on a test-set example in which 3DisoDeepPF predictions agreed with the curated Pfam and GO-MF annotations (**Supplementary Note 7**). The evidence-tracing utility identified the top 15 neighboring nodes providing the strongest support for the predicted Pfam domains and GO-MF terms (**Fig. 2g**; **Extended Data Fig. 5c**). For GO-MF, these supporting neighbors were enriched for proteins associated with calcium-related functions (**Extended Data Fig. 5b**). Notably, the supporting nodes were not confined to the immediate neighborhood of CIB1, indicating that 3DisoDeepPF can draw on broader graph context beyond the closest neighbors. To visualize the prediction results of 3DisoDeepPF within the graph framework used as model input, the graph embedding showed that predicted Pfam labels closely matched UniProtKB annotations, both in the full dataset and the held-out test set (**Extended Data Fig. 6a**). We also provide standardized knowledge bases and a web portal for browsing the embedding space, inspecting predicted structures, and accessing cross-linked views of function-matched isoforms (**Extended Data Fig. 6b,c**). These resources may facilitate downstream candidate prioritization and biomarker-oriented hypothesis generation in isoform-resolved studies. Together, they support an isoform-centric framework for Pfam domain and GO term annotation at isoform resolution, instantiated in 3DisoDeepPF with matched evaluation schemes.

---

**Extended Data Figure 1.**
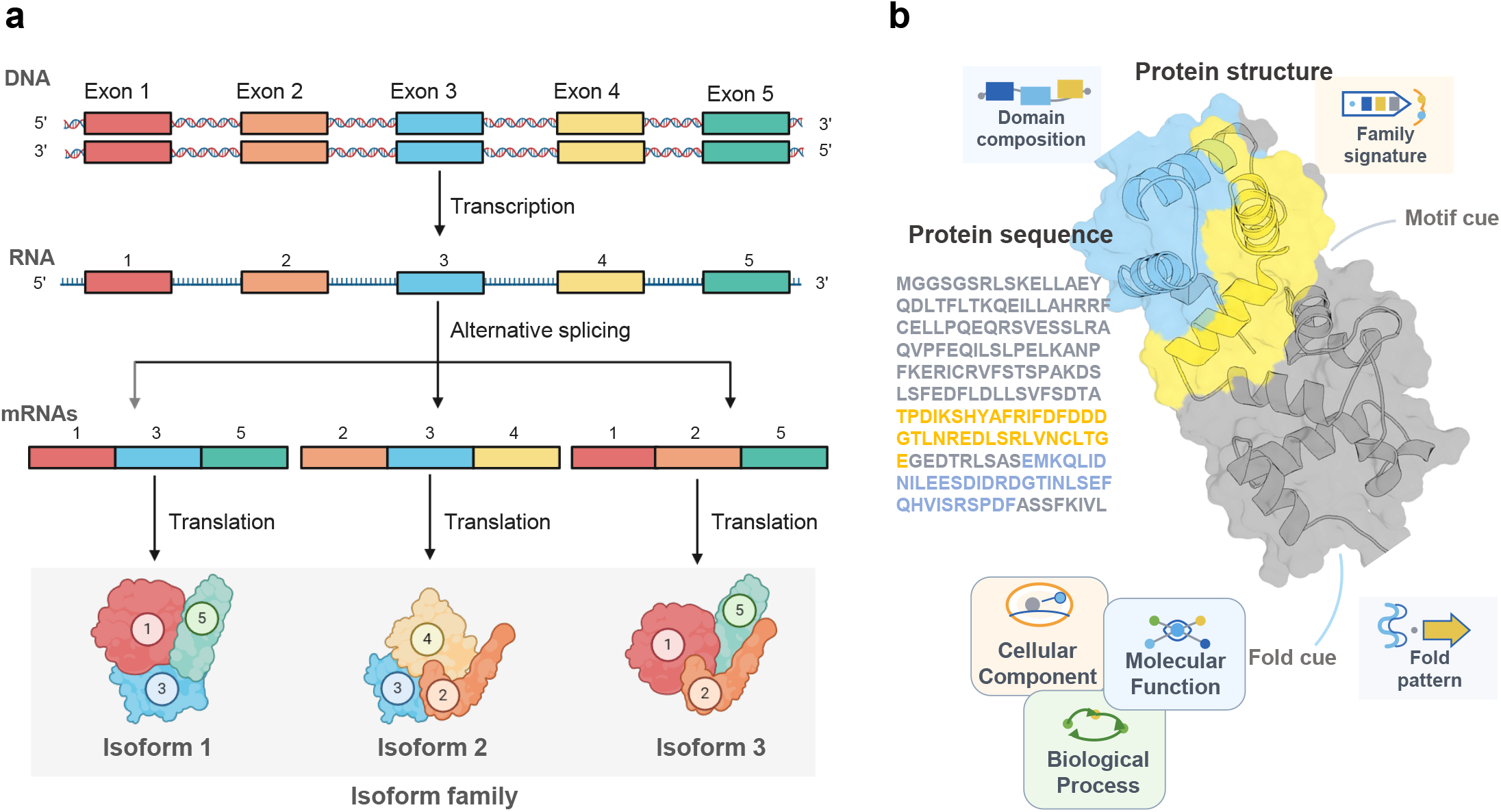
Alternative splicing-driven isoform generation and conceptual rationale for isoform annotation. **a**, Schematic illustration of alternative splicing-driven generation of multiple protein isoforms from a single gene, forming an isoform family. **b**, Conceptual schematic illustrating how protein sequence and structure provide complementary cues for domain and function annotation, including domain composition, family-level signatures, motif features and fold-related patterns, which together inform Pfam and Gene Ontology annotation.

**Extended Data Figure 2.**
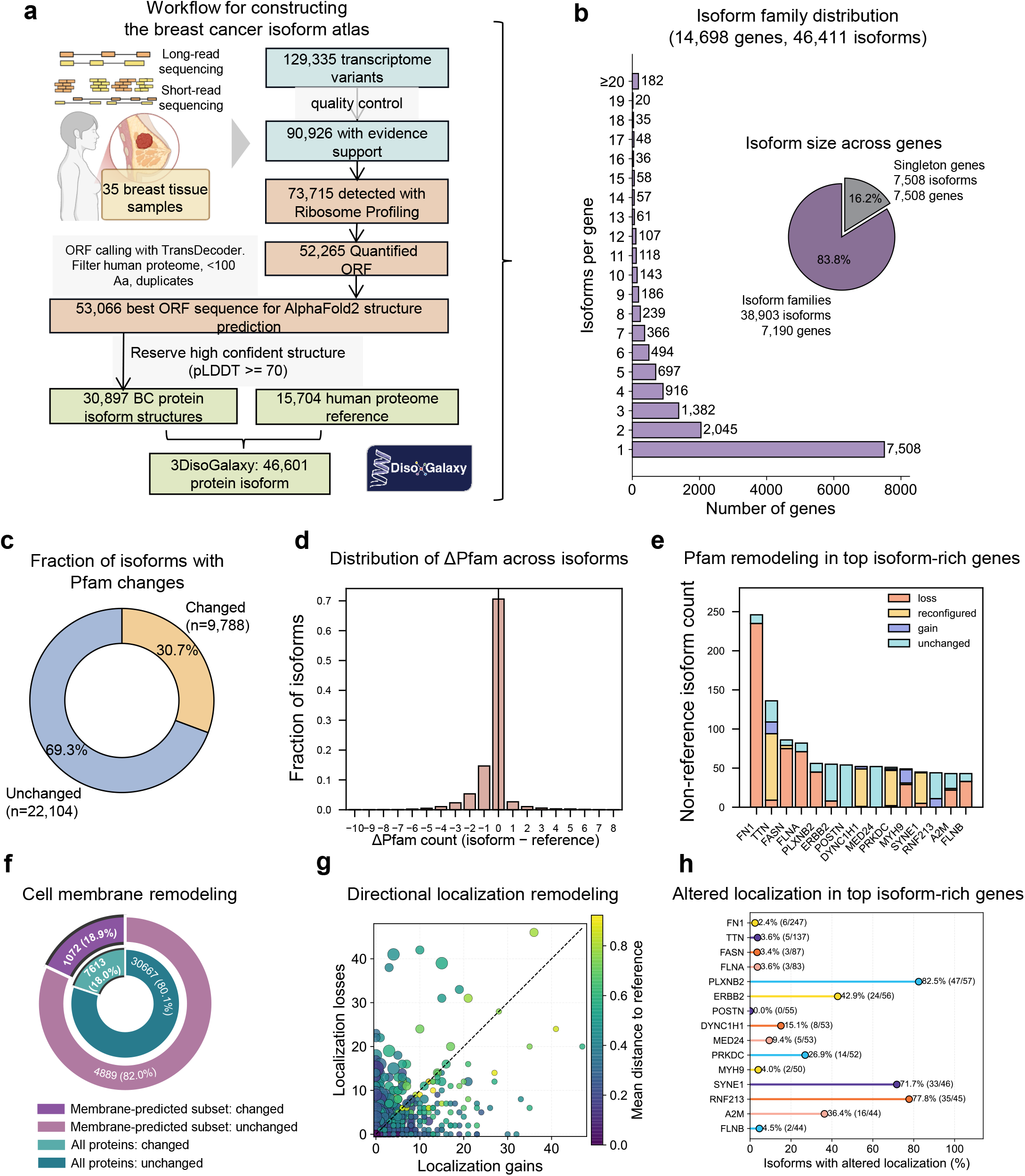
Atlas construction and proxy evidence for isoform remodeling. **a**, Workflow for construction of the 3DisoGalaxy breast cancer isoform atlas. **b**, Distribution of isoform family sizes across the atlas. **c**, Fraction of non-reference isoforms with Pfam changes relative to the reference isoform. **d**, Distribution of ΔPfam counts across non-reference isoforms. **e**, Pfam remodeling categories in the top 15 isoform-rich genes. **f**, Summary of predicted membrane-localization remodeling. **g**, Directional remodeling of inferred localization labels relative to the reference isoform; color indicates mean Jaccard distance between label sets. **h**, Fraction of non-reference isoforms with altered inferred localization labels in the top 15 isoform-rich genes. These atlas-scale proxies support isoform-aware modeling, but do not constitute experimental ground truth.

**Extended Data Figure 3.**
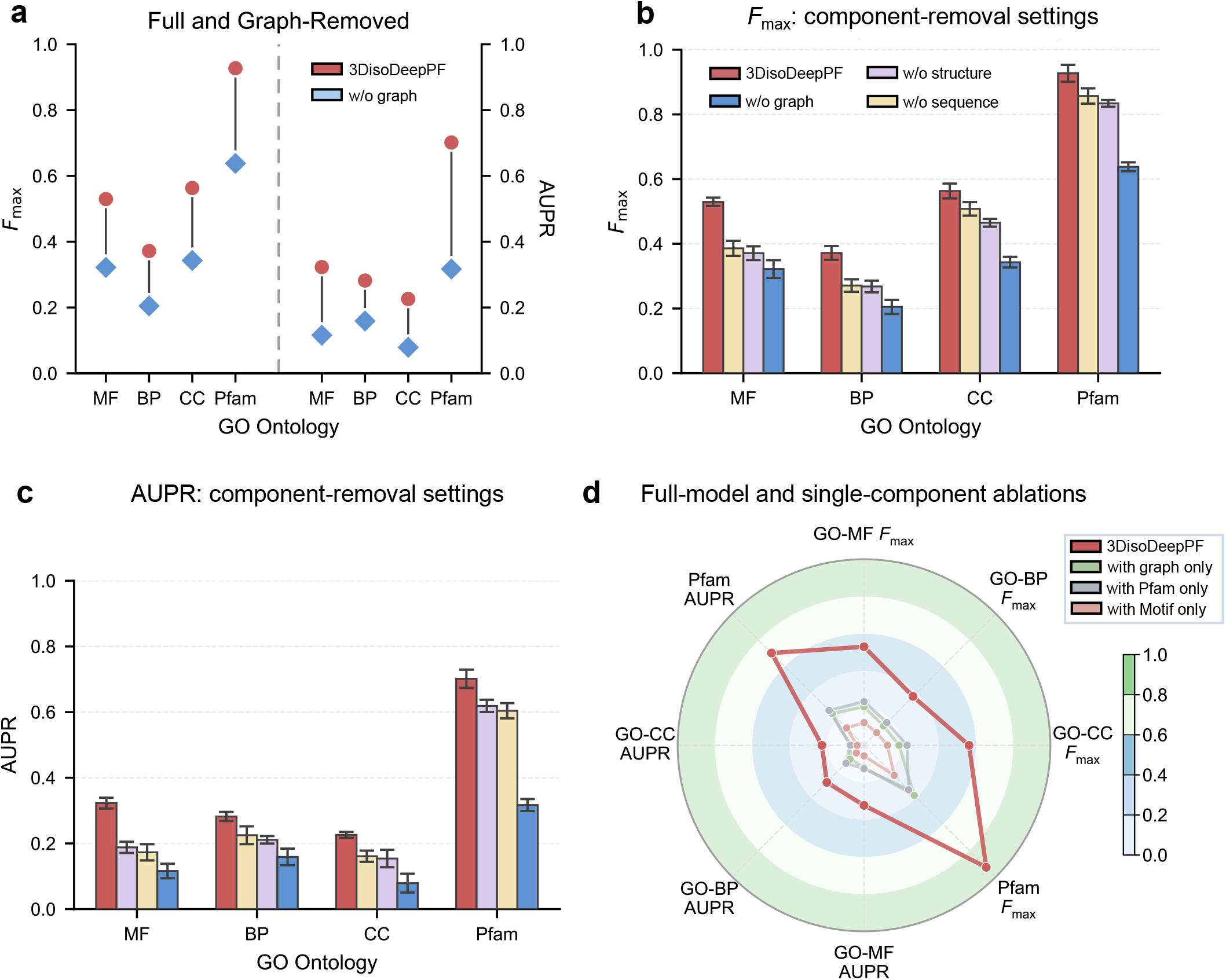
Ablation analysis of graph and feature contributions. **a**, Comparison of the full 3DisoDeepPF model and a graph-removed variant across GO-MF, GO-BP, GO-CC and Pfam prediction tasks. Left, protein-centric Fmax; right, function-centric AUPR. Red circles denote 3DisoDeepPF and blue diamonds denote the graph-removed model. Vertical lines indicate absolute differences between the two models. **b**,**c**, Performance of the full model and feature-removal variants after removing the sequence– structure graph, structure features or sequence features. **b**, *F*_max_. **c**, AUPR. Error bars indicate variability across repeated runs. **d**, Radar plot comparing 3DisoDeepPF with single-component models using graph-only, Pfam-only or motif-only inputs across GO-MF, GO-BP, GO-CC and Pfam prediction tasks. Axes indicate task–metric pairs evaluated by Fmax or AUPR. Radial values range from 0 to 1, with larger values indicating higher performance.

**Extended Data Figure 4.**
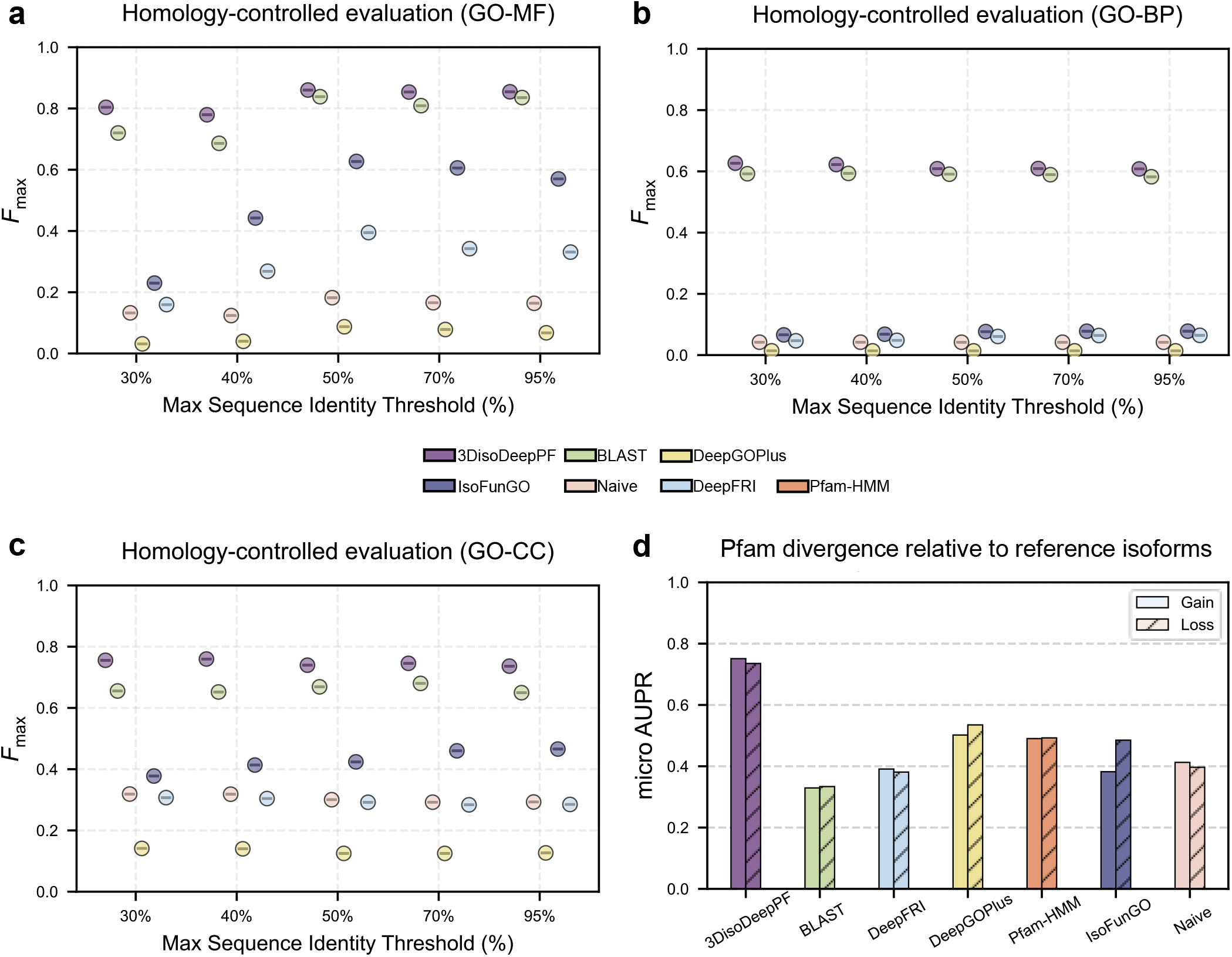
Homology-controlled evaluation and directional Pfam remodeling. **a–c**, Homology-controlled evaluation across maximum train–test sequence-identity thresholds, shown separately for GO-MF (**a**), GO-BP (**b**) and GO-CC (**c**). Each color denotes a different method, as indicated in the legend, and the y axis shows *F*_max_. **d**, Micro-AUPR for directional Pfam remodeling within isoform families relative to the selected reference isoform. Gain denotes Pfam domains present in an isoform but absent from the reference isoform, whereas loss denotes domains present in the reference isoform but absent from the isoform. Bar color denotes method, and fill pattern distinguishes gain and loss, as indicated in the legend.

**Extended Data Figure 5.**
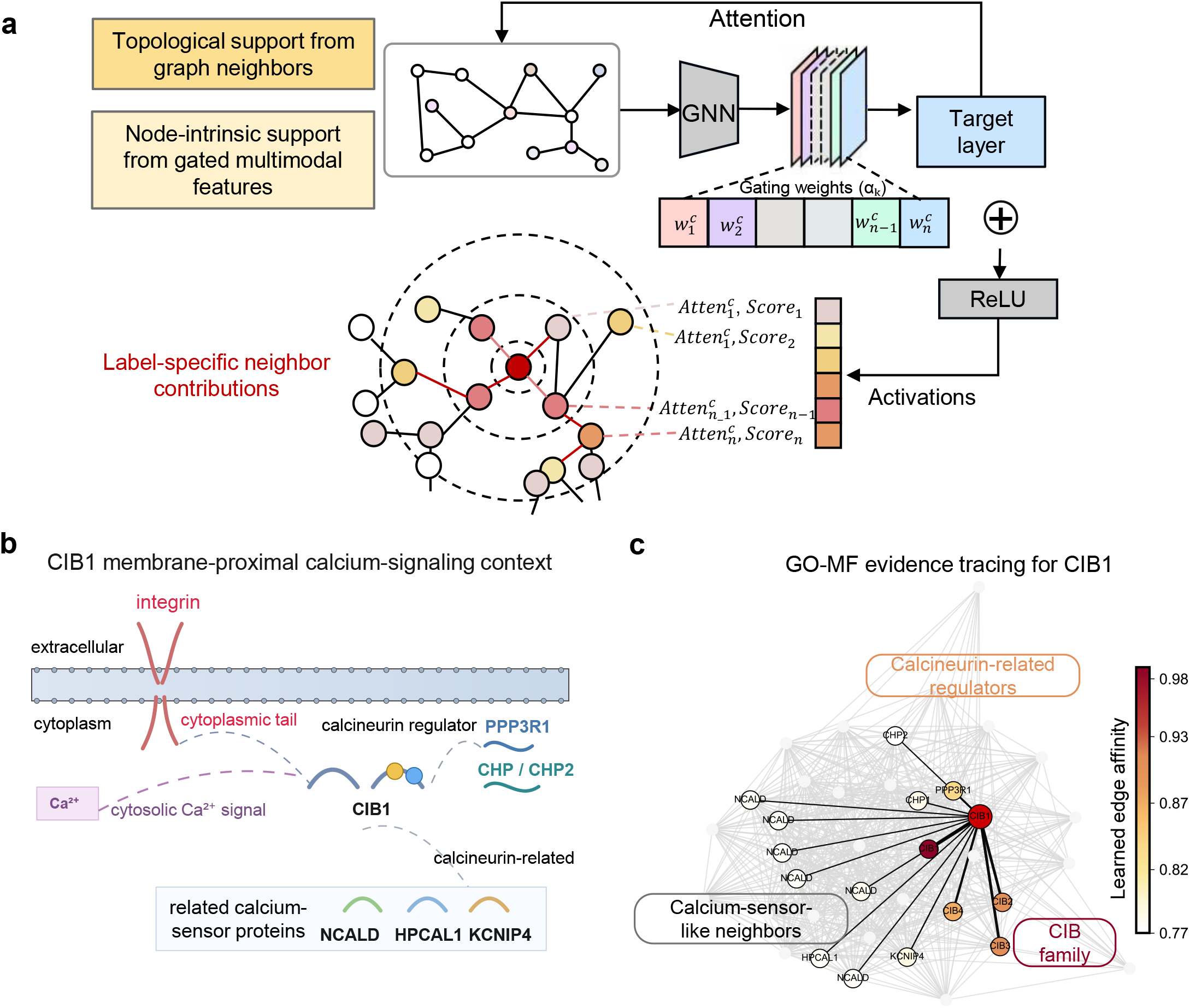
Evidence-tracing utilities for 3DisoDeepPF. **a**, Schematic overview of the evidence-tracing framework, showing how each prediction is decomposed into topological support from graph-associated protein nodes and node-intrinsic support from gated multimodal features. **b**, A schematic overview of the membrane-proximal calcium-signaling context of CIB1, including integrin-associated, calcineurin-related and calcium-sensor-related proteins. **c**, GO-MF evidence tracing for CIB1, showing CIB1 and traced protein nodes most strongly associated with the predicted GO-MF labels; node color summarizes learned edge affinity to CIB1, with darker color indicating stronger model-inferred relatedness.

**Extended Data Figure 6.**
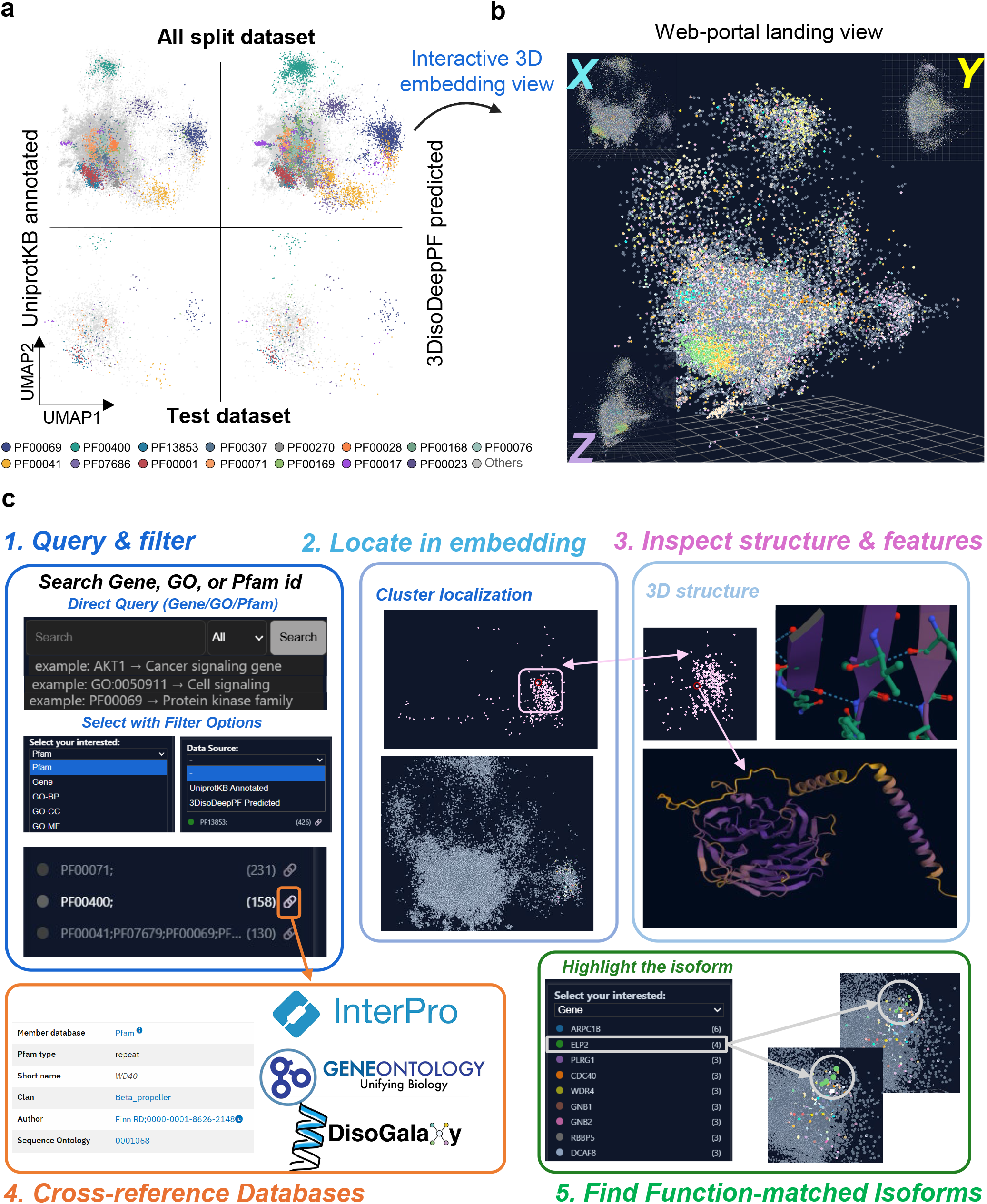
Hyperbolic embedding visualization and portal workflow. **a**, Hyperbolic UMAP of the 3DisoGalaxy atlas colored by the 15 most frequent Pfam domains, shown for the full and held-out test sets with curated UniProtKB/Swiss-Prot annotations and 3DisoDeepPF predictions in parallel. **b**, Web-portal landing view with an interactive 3D embedding for navigation and neighborhood inspection. **c**, Portal workflow for query/filter, locating hits in the embedding, inspecting structures and features, cross-referencing external resources (e.g., InterPro and Gene Ontology), and retrieving function-matched isoforms.

## Methods

### An overview of isoform-resolution protein function prediction

This study introduces a framework to predict protein function at the isoform level, with an addition of a CAFA-aligned evaluation in the conventional mode. This allowed performance to be assessed both under standard community conditions and in a biologically more relevant setting enriched for isoform diversity. The task was formulated as a multi-label node classification problem on a graph, in which each node represents a protein or isoform and edges encode integrated sequence and structural similarity. Evaluation focused on Pfam domain and GO term prediction, using established community metrics together with homology-aware assessment and isoform-family directional remodeling analysis, in which isoforms from the same gene were compared against a predefined reference isoform to evaluate domain gain and loss within isoform families. For human data, a random split was used due to the close release times of isoform sequences, which made traditional temporal splits ineffective. Leakage control was implemented by ensuring strict separation of isoforms from the same gene or UniProt accession across training, validation, and test sets, and by applying modality isolation to prevent cross-task leakage.

### CAFA-aligned benchmark curation

#### Benchmark Curation: Sequence, GO Annotations, Pfam Domains, Motifs, and Evidence-based Filtering

We curated a CAFA-aligned benchmark to enable reproducible evaluation on a public dataset (Radivojac et al., 2013). We obtained 573,661 protein sequences from UniProtKB/Swiss-Prot (release 2025_04) (UniProt Consortium, 2025), functional annotations from the Gene Ontology Annotation (GOA) database (Camon et al., 2004), and the GO graph from the go-basic.obo release dated 2025-07-22 (The Gene Ontology Consortium, 2026). We retained only annotations supported by experimental or curated evidence codes (EXP, IDA, IPI, IMP, IGI, IEP, HDA, HMP, HGI, HEP, TAS, IC). We excluded any electronically inferred or low-confidence annotations (IEA, ND, NAS) as well as negated annotations (Camon et al., 2004; The Gene Ontology Consortium, 2026). Proteins were included if a Swiss-Prot sequence was available and at least one GO term in MF, BP, or CC remained after evidence filtering and ontology propagation. We also retrieved Pfam domain and short linear motif (SLiM) annotations from UniProtKB/Swiss-Prot as auxiliary features (Mistry et al., 2021). The final benchmark includes 76,804 proteins across all species.

#### Structure-aware preparation and analysis

We assembled a unified structural compendium by integrating experimentally determined protein structures from the Protein Data Bank (PDB; Kouranov et al., 2006) with computationally predicted models from the AlphaFold Database (Varadi et al., 2024). We mapped PDB chains to canonical UniProt accessions using the Structure Integration with Function, Taxonomy and Sequences (SIFTS) resource (Dana et al., 2019). We resolved multi-chain PDB entries at the individual-chain level. When multiple experimental structures mapped to the same accession, we retained a single representative structure by ranking candidates based on experimental method, nominal resolution, and chain residue count. This yielded 29,392 experimental representative structures. For proteins lacking experimental coverage, we supplemented the compendium with AlphaFold models, retaining 47,272 structures with mean pLDDT ≥ 70 for primary analyses and 16,928 structures with 60 ≤ pLDDT < 70 for sensitivity analyses. Models with pLDDT < 60 were excluded.

#### CAFA-aligned temporal split

We followed the CAFA time-delayed no-knowledge (TDNK) protocol, in which models are trained only on annotations available before a fixed historical cutoff and evaluated on annotations that became available later (Radivojac et al., 2013; Jiang et al., 2016; Zhou et al., 2019). We used two fixed cutoffs anchored to GOA timestamps: the training visibility cutoff t_0_ = 2024-04-01 and the evaluation horizon t_1_ = 2025-09-01, using a GOA snapshot as of 2025-09-01. All labels dated after t_1_ were excluded from both training and evaluation. True-path propagation of GO terms was then applied separately within each time window: labels of train/validation datasets were propagated from atomic annotations dated ≤ t_0_, whereas test gold labels were propagated from atomic annotations dated within (t_0_, t_1_].

### Breast cancer isoform-centric data processing

#### Construction of an isoform-specific breast cancer structural atlas

We constructed a breast cancer-specific protein isoform dataset, 3DisoGalaxy, integrating PacBio Iso-Seq, RNA-seq, and Ribo-seq data to characterize isoform variations (Jiang et al., 2026; https://github.com/FeliciaTJiang/3DisoGalaxy). This dataset integrates PacBio Iso-Seq (Veiga et al., 2022) (*n* = 35), short-read RNA-seq from five cohorts (Li et al., 2024; Jiang et al., 2019; Varley et al., 2014; Nayarisseri et al., 2024) (*n* = 339), and Ribo-seq datasets (Navickas et al., 2023; Vaklavas et al., 2020) (*n* = 42). We identified and quantified open reading frame-coding isoforms using ribosome profiling analysis and obtained their translated amino acid sequences. Three-dimensional structures were predicted using AlphaFold (v2.3) (Jumper et al., 2021). We identified 129,335 transcript variants via long-read sequencing, with 90,926 supported by orthogonal short reads. We detected translation signals for 73,715 isoforms, yielding 53,066 predicted protein structures. After integrating 15,704 high-confidence human proteome reference structures and excluding predictions with pLDDT < 70, we retained 46,411 high-confidence structures. We curated Pfam, GO, and SLiM annotations using the same multimodal feature-processing procedure described for the CAFA-aligned benchmark.

#### Supervised data splits

Unlike the CAFA-aligned benchmark, we did not evaluate 3DisoGalaxy using a temporal split, as temporally ordered partitioning produced an artificial discontinuity in label coverage (**Supplementary Note 4**). Instead, we applied ontology-specific stratified random splits for GO-MF, GO-BP, GO-CC, and Pfam. For each task, labeled isoforms were partitioned into training, validation, and test sets using an 8:1:1 ratio. Unlabeled isoforms were excluded from loss computation and used only for inference. To prevent leakage, all isoforms assigned to the same UniProt accession were confined to a single split. We performed model selection on the validation split, evaluated the final model on the held-out test split, and applied it to the unlabeled subset for inference. Detailed data cleaning procedures, dataset splitting criteria and final dataset statistics are provided in **Supplementary Note 4**.

### 3DisoDeepPF architecture and training

#### Multimodal graph construction and representation

We formulated protein function prediction as a node classification task on a multimodal similarity graph *G* = (*V, E, X*). To construct the node features *X*, we extracted and integrated distinct modalities: sequence embeddings from protein language models ESM (*h*_sequence_) (Lin et al., 2023) (obtained via mean-pooling over all residues), short linear motifs (*h*_motif_), available functional annotations (*h*_annotation_), and a learnable base node embedding (*h*_learn_). The *h*_annotation_ modality incorporates known labels for the target task and cross-task auxiliary information. We projected each modality into a joint feature space and concatenated them to form the initial node representation 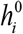for protein *i*.

To construct the edges *E*, we computed pairwise sequence homologies (*A*_seq_) using BLASTP and structural similarities (*A*_str_) using Foldseek in TM-align mode. We integrated these into a unified adjacency matrix *A* using a learned mixing parameter *λ* ∈[0,1] :

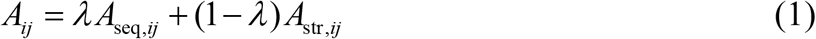

We applied task-specific thresholds *τ* to sparsify the graph, retaining only edges where *A*_*ij*_> *τ* to mitigate noise and optimize computational efficiency. In this study, we used τ = 0.3 to construct a sparse graph (details in **Supplementary Note 5**). This threshold was selected empirically from a range of candidate cutoffs because it balanced graph sparsity with retention of informative edges, preserving neighborhood structure while limiting weaker connections that could dilute biologically coherent message passing.

#### Gated multimodal feature fusion

We updated the initial node representations through graph attention layers to capture local structural and functional contexts. For a given modality at layer *l*, the hidden state of protein *i* is updated by aggregating features from its neighborhood *N* (*i*) :

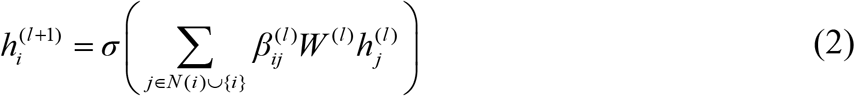

where *σ* denotes a nonlinear activation function and *W*^(*l*)^ is a learnable weight matrix. The attention coefficient 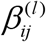determines the importance of neighbor *j* to protein *i*, computed via a LeakyReLU activation over the concatenated features.

Following the final propagation layer *L*, we integrated the updated representations using a gated fusion module. Let *h*_*k*_ denote the final layer output *h*^(*L*)^ for a specific modality *k*. The fused representation *h*_final_ for each node is defined as:

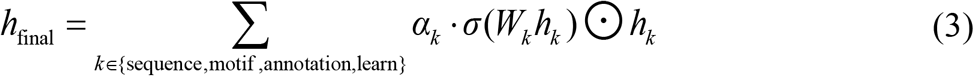

where *α*_*k*_ represents softmax-normalized gating weights that dynamically adjust the contribution of each modality during prediction, and ⊙ denotes the element-wise product.

#### Task-adaptive prediction ensemble

We mapped the fused graph representations to functional predictions using an ensemble of prediction heads. Each head processes the identical node embedding to independently generate logits for target labels. We combined these outputs using learnable weights to yield a consensus prediction. A sigmoid activation applied to the aggregated logits produces the final multi-label probabilities *p*_*i,c*_ for protein *i* and class *c*. To address the label imbalance inherent in protein function datasets we optimized the model using a class-weighted focal loss *L*:

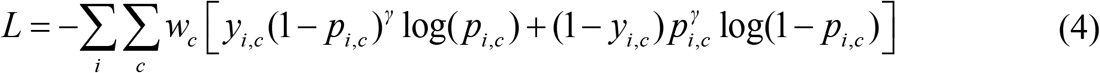

where *y*_*i,c*_ ∈ {0,1} is the ground truth label, *w*_*c*_ is the class-specific weight, defined as the ratio of negative to positive training examples for class *c* (capped to ensure training stability), and *γ* is the focusing parameter that down-weights the loss assigned to easily classified examples. We selected decision thresholds on the validation set to maximize the *F*_max_ metric.

#### Topological and modal attribution reporting (evidence tracing)

We developed utilities to decompose prediction evidence into topological support from graph neighbors and node-intrinsic support from multimodal features. For a predicted label *y* on protein *i*, we ranked supporting neighbors *j* using the model-learned adaptive edge weights *A*_*i, j*_, which encapsulate the integrated sequence-structure similarity network. The term *A*_*i, j*_ represents the model-learned adaptive edge weight that explicitly quantifies the contextual importance and the intensity of functional information propagation between connected nodes during graph convolution. We reported the top *K* scoring neighbors with the highest adaptive weights as the primary topological evidence.

### Evaluation regimes

#### Conventional canonical protein benchmarking

We used the CAFA-aligned benchmark for standard protein annotation evaluation. We generated predictions independently for GO-MF, GO-BP, GO-CC, and Pfam, measuring performance via maximum F-score (*F*_max_) and area under the precision recall curve (AUPR). We compared 3DisoDeepPF against representative baselines spanning homology transfer, domain transfer, naive frequency-based prediction, and graph-based learning methods, all evaluated under identical data partitions and ontology processing rules (**Supplementary Note 4, 6**).

#### Homology-controlled evaluation

We conducted homology-controlled evaluations by imposing maximum sequence identity constraints between training and test proteins. We defined separate evaluation sets at identity cutoffs of 30%, 40%, 50%, 70%, and 95%. For each task, any training or validation protein exceeding the specified cutoff relative to a test protein was excluded. We then evaluated GO and Pfam prediction independently at each cutoff.

#### Isoform-family directional remodeling evaluation

To assess the resolution of functional variation among isoforms from the same gene, we defined a directional remodeling task based on reference-relative Pfam changes. For each isoform family, we used the MANE isoform as the reference. Families lacking a MANE (Morales et al., 2022) annotation were anchored to the UniProtKB/Swiss-Prot canonical isoform. We compared each non-reference isoform with its reference, defining Pfam gain as Pfam labels (PF accessions) present in the query isoform but absent from the reference, and Pfam loss as Pfam labels present in the reference but absent from the query isoform. Presence was defined at the level of Pfam label membership rather than full-versus partial-domain coverage. Isoform to reference pairs were stratified by sequence identity into predefined bins. We evaluated directional remodeling performance separately for gain and loss using micro-AUPR. We also quantified the fraction of pairs showing any Pfam change, domain loss, or domain gain across identity bins to map the support landscape.

#### Ablation analysis of graph topology and multimodal integration

We evaluated component contributions in the isoform-resolved setting using two ablation strategies. In the component removal setting, we independently removed the similarity graph, structure, sequence, Pfam, and motif inputs from the full model. We quantified performance drops using Δ*F*_max_ and ΔAUPR. In the single-component setting, we evaluated isolated inputs to assess the predictive signal contributed by each channel alone.

### Leakage control and validation design

Split isolation for breast cancer isoform-centric dataset is described above under *Supervised data splits*, and temporal no-knowledge and homology-controlled evaluations are described above under Evaluation regimes. The controls below address target-label isolation, graph construction and information flow, and model selection/preprocessing.

#### Target-label isolation

Target-task annotations were excluded from the corresponding prediction task to avoid direct label leakage. For GO-MF, GO-BP, and GO-CC predictions, annotations from the target ontology were not used as input features for that task. For Pfam prediction, Pfam labels were excluded from the corresponding task input. When auxiliary label-derived features were constructed, labels from validation and test nodes were masked for the corresponding prediction task. Task-specific masking rules are summarized in **Supplementary Note 4**.

#### Graph construction and transductive training protocol

We constructed the similarity graph exclusively from sequence and structure similarities. Gene Ontology and Pfam labels served strictly as prediction targets and did not define graph connectivity. To ensure rigorous evaluation hygiene and prevent look-ahead shortcuts, all functional annotations corresponding to the validation and test partitions were strictly masked and withheld from the optimization objective. This layout constrains the loss computation and gradient updates to depend exclusively on the labeled training partition, while preserving the structural context of the global similarity network. Only labels from the training partition contributed to the supervised loss and gradient updates. We used validation labels solely for hyperparameter tuning and model checkpoint selection, reserving test labels exclusively for final evaluation to prevent target label leakage. To maintain strict data hygiene, any preprocessing steps dependent on dataset statistics were derived solely from the training partition and applied unchanged to the validation and test partitions.

### Visualization and hyperbolic embedding

#### Hyperbolic embedding of the breast-cancer isoform similarity graph

For visualization, we used the breast-cancer isoform sequence–structure similarity graph constructed with the same edge definition and cutoff as the graph used for 3DisoDeepPF model input. Weighted node embeddings were learned with node2vec using 64 dimensions. The resulting embeddings were projected into Lorentz-model hyperbolic space through a tangent-space formulation, and the three-dimensional coordinates were used for interactive visualization of annotation and prediction consistency. To assess whether the embedding preserved biologically meaningful organization, we applied k-means clustering to the embedded coordinates and inspected the resulting clusters for annotation coherence.

#### Web portal for isoform-centric visualization

We developed an interactive portal at http://3disodeeppf.com/ for querying proteins and isoforms, browsing predicted Pfam and GO annotations, comparing isoforms from the same gene and inspecting selected protein structures. Protein structures are rendered in-browser using Mol* (Sehnal et al., 2021).

## Data availability

All data required to evaluate the conclusions of this study are provided within the paper and its Supplementary Information. Curated datasets, model outputs and additional materials are publicly available through the 3DisoGalaxy portal (http://3disogalaxy.com/) and the 3DisoDeepPF GitHub repository (https://github.com/FeliciaTJiang/3DisoDeepPF). Model predictions generated in this study are accessible through the interactive 3DisoDeepPF portal at http://3disodeeppf.com/, with documentation and usage tutorials available at http://3disodeeppf.com/about.html.

## Code availability

Source code and pre-trained models for running 3DisoDeepPF are freely available for non-commercial academic use at https://github.com/FeliciaTJiang/3DisoDeepPF. The analysis R and Python scripts for bioinformatic analysis are available at https://github.com/FeliciaTJiang/3DisoGalaxy.

## Notes

http://www.3disodeeppf.com/

### Competing Interest Statement

The authors have declared no competing interest.

### Summary of Updates

This version represents the submitted manuscript, in which the title and abstract have been revised.

## References

Friedberg, I. Automated protein function prediction—the genomic challenge. Brief. Bioinform. 7, 225–242 (2006).

Radivojac, P. et al. A large-scale evaluation of computational protein function prediction. Nat. Methods 10, 221–227 (2013).

Jiang, Y. et al. An expanded evaluation of protein function prediction methods shows an improvement in accuracy. Genome Biol. 17, 184 (2016).

Zhou, N. et al. The CAFA challenge reports improved protein function prediction and new functional annotations for hundreds of genes through experimental screens. Genome Biol. 20, 244 (2019).

The UniProt Consortium. UniProt: the universal protein knowledgebase in 2025. Nucleic Acids Res. 53, D609–D617 (2025).

Jumper, J. et al. Highly accurate protein structure prediction with AlphaFold. Nature 596, 583–589 (2021).

Smith, L. M., Kelleher, N. L. & The Consortium for Top Down Proteomics. Proteoform: a single term describing protein complexity. Nat. Methods 10, 186–187 (2013).

Graveley, B. R. Alternative splicing: increasing diversity in the proteomic world. Trends Genet. 17, 100–107 (2001).

Nilsen, T. W. & Graveley, B. R. Expansion of the eukaryotic proteome by alternative splicing. Nature 463, 457–463 (2010).

Weatheritt, R. J., Sterne-Weiler, T. & Blencowe, B. J. The ribosome-engaged landscape of alternative splicing. Nat. Struct. Mol. Biol. 23, 1117–1123 (2016).

Liu, Y. et al. Impact of alternative splicing on the human proteome. Cell Rep. 20, 1229–1241 (2017).

Mudge, J. M. et al. GENCODE 2025: reference gene annotation for human and mouse. Nucleic Acids Res. 53, D966–D975 (2025).

Mistry, J. et al. Pfam: The protein families database in 2021. Nucleic Acids Res. 49, D412–D419 (2021).

Jiang, F. T. et al. The Structural Code of Breast Cancer Proteoform: Alternative Splicing-driven Protein Isoform Variation and Functional Diversification. Preprint at bioRxiv 10.64898/2026.04.30.722115 (2026).

Yu, G., Wang, K., Domeniconi, C., Guo, M. & Wang, J. Isoform function prediction based on bi-random walks on a heterogeneous network. Bioinformatics 36, 303–310 (2020).

Qiu, S., Yu, G., Lu, X., Domeniconi, C. & Guo, M. Isoform function prediction by Gene Ontology embedding. Bioinformatics 38, 4581–4588 (2022).

Camon, E. et al. The Gene Ontology Annotation (GOA) Database: sharing knowledge in Uniprot with Gene Ontology. Nucleic Acids Res. 32, D262–D266 (2004).

The Gene Ontology Consortium. The Gene Ontology knowledgebase in 2026. Nucleic Acids Res. 54, D1779–D1792 (2026).

Kouranov, A. et al. The RCSB PDB information portal for structural genomics. Nucleic Acids Res. 34, D302–D305 (2006).

Dana, J. M. et al. SIFTS: updated Structure Integration with Function, Taxonomy and Sequences resource allows 40-fold increase in coverage of structure-based annotations for proteins. Nucleic Acids Res. 47, D482–D489 (2019).

Veiga, D. F. T. et al. A comprehensive long-read isoform analysis platform and sequencing resource for breast cancer. Sci. Adv. 8, eabg6711 (2022).

Li, S.-Y. et al. Tumor circadian clock strength influences metastatic potential and predicts patient prognosis in luminal A breast cancer. Proc. Natl. Acad. Sci. U.S.A. 121, e2311854121 (2024).

Jiang, Y.-Z. et al. Genomic and Transcriptomic Landscape of Triple-Negative Breast Cancers: Subtypes and Treatment Strategies. Cancer Cell 35, 428–440.e5 (2019).

Varley, K. E. et al. Recurrent read-through fusion transcripts in breast cancer. Breast Cancer Res. Treat. 146, 287–297 (2014).

Nayarisseri, A. et al. Transcriptomic Analysis of Primary Breast Cancer Utilizing Gene Expression Datasets from Middle-aged Caucasian Women. Research Square preprint, 10.21203/rs.3.rs-4701356/v1 (2024).

Navickas, A. et al. An mRNA processing pathway suppresses metastasis by governing translational control from the nucleus. Nat. Cell Biol. 25, 892–903 (2023).

Vaklavas, C., Blume, S. W. & Grizzle, W. E. Hallmarks and Determinants of Oncogenic Translation Revealed by Ribosome Profiling in Models of Breast Cancer. Transl. Oncol. 13, 452–470 (2020).

Varadi, M. et al. AlphaFold Protein Structure Database in 2024: providing structure coverage for over 214 million protein sequences. Nucleic Acids Res. 52, D368–D375 (2024).

Lin, Z. et al. Evolutionary-scale prediction of atomic-level protein structure with a language model. Science 379, 1123–1130 (2023).

Morales, J. et al. A joint NCBI and EMBL-EBI transcript set for clinical genomics and research. Nature 604, 310–315 (2022).

Sehnal, D. et al. Mol* Viewer: modern web app for 3D visualization and analysis of large biomolecular structures. Nucleic Acids Res. 49, W431–W437 (2021).

