## Supplementary Notes for "Structure-aware protein function prediction at isoform resolution"

#### Contents

**1 Supplementary Note 1 | Design rationale for isoform-resolution protein function prediction ... *4***

1.1 Isoform-resolution protein function prediction as a distinct computational problem..4

1.2 Evaluation logic for isoform-resolution protein function prediction ... 5

1.3 From structure-based clustering to graph-based function inference ... 5

1.4 Framework implementation in 3DisoDeepPF ... 6

**2 Supplementary Note 2 | Interpretation of 3DisoDeepPF results and implications ... *7***

2.1 Biological and structural basis of isoform functional remodeling ... 7

2.2 Canonical benchmark performance as a standard protein function prediction competence test ... 8

2.3 Isoform-resolution performance in the breast cancer isoform atlas ... 9

2.4 Evidence tracing and interpretability of individual predictions ... 10

2.5 Why graph-centered multimodal integration improves isoform-resolution prediction ... 11

2.6 Relationship to existing protein function prediction methods ... 13

**3 Supplementary Note 3 | Structure-aligned knowledge bases and functional proxy annotation ... *14***

3.1 Benchmark construction and structure-support quality of CAFA-aligned canonical dataset ... 14

3.2 Composition and annotation coverage of the 3DisoGalaxy structure atlas ... 15

3.3 Pfam and localization prediction with existing tools ... 16

**4 Supplementary Note 4 | Dataset construction, partitioning and leakage-control design ... *16***

4.1 CAFA-aligned canonical benchmark construction and partitions ... 17

4.1.1 Raw dataset collection and protein inclusion ... 17

4.1.2 Structural coverage and representative selection ... 17

4.1.3 Ontology-specific split statistics ... 18

4.1.4 Overview of a multi-species, structure-annotated CAFA-aligned benchmark ... 18

#### List of Supplementary Figures

1 Micro-averaged precision–recall performance across canonical and isoform-resolved benchmarks

2 Overview of the CAFA-aligned canonical protein benchmark

3 Composition, structural quality and annotation coverage of the breast cancer isoform atlas

4 Cutoff selection for graph construction on the CAFA-aligned benchmark

5 Cutoff selection for graph construction on the 3DisoGalaxy evaluation set

6 Integrated protein isoform similarity graph and representative local neighborhoods

7 Interactive hyperbolic visualization of curated and predicted Pfam annotations across the integrated isoform similarity graph

8 Fmax profiles across canonical and isoform-resolved benchmarks

9 Predicted label coverage and no-prediction-associated features across Pfam and Gene Ontology label spaces

10 Predicted isoform-level annotation remodeling and cancer-driver association

11 GO-label divergence in tumor-associated non-canonical protein isoform candidates

12 Pfam-domain divergence in tumor-associated non-canonical protein isoform candidates

### 1 Supplementary Note 1 | Design rationale for isoform-resolution protein function prediction

#### 1.1 Isoform-resolution protein function prediction as a distinct computational problem

Protein function is mediated by specific protein molecules rather than by genes alone. These molecular forms, often referred to as proteoforms, can arise from genetic variation, alternative splicing, post-translational modification and other molecular processes, creating functional diversity that is not fully captured by a single canonical protein sequence (Smith and Kelleher, 2013; Aebersold et al., 2018; Smith et al., 2018). However, most computational protein function prediction frameworks still assign function in a gene-centric or canonical/reference-protein setting, leaving proteoform-level functional differences largely unresolved.

In this study, we focus on alternative-splicing-derived protein isoforms as a systematic and structurally analyzable entry point into proteoform-resolution function prediction. Alternative splicing enables one gene to produce multiple transcript variants, a subset of which can be translated into distinct protein isoforms. These isoforms can differ in domain composition, regulatory motifs, interaction regions and expression or abundance (Nilsen and Graveley, 2010; Weatheritt et al., 2016; Liu et al., 2017). Tissue-regulated alternative exons can remodel protein-interaction networks, indicating that isoform differences may affect molecular function rather than only transcript structure (Ellis et al., 2012). Dysregulated splicing is also widely implicated in human disease, including cancer, where altered isoforms can affect tumor growth, metastasis and therapeutic vulnerabilities (Scotti and Swanson, 2016; Urbanski et al., 2018; Zhang et al., 2021).

This creates a specific challenge for protein function prediction. Isoforms from the same gene are often highly homologous and share substantial functional background, so standard sequence-transfer or learning-based models may appear accurate by recovering shared gene-level annotations. Yet functionally relevant differences may arise from local truncation, exon skipping, domain gain or loss, altered motifs or remodeled structural regions. These changes may involve only a small part of the protein, but they can affect the molecular features that determine function, localization or interaction specificity.

#### 1.2 Evaluation logic for isoform-resolution protein function prediction

Evaluation presents a central challenge for isoform-resolution protein function prediction. Isoforms from the same gene are often highly homologous and share substantial functional background, making apparent prediction performance susceptible to inflation through homology-driven annotation transfer. Under such conditions, successful recovery of shared labels does not necessarily indicate that a method can resolve isoform-specific functional remodeling. We therefore treated evaluation not only as a performance-reporting step, but as a core component of the framework design.

To address different sources of evaluation ambiguity, we organized the evaluation framework into four complementary layers. First, we used CAFA-aligned canonical protein benchmarks to assess whether the framework remained competitive under conventional protein function prediction settings. Second, we implemented a time-delayed no-knowledge protocol to evaluate performance under temporally realistic annotation visibility and reduce future annotation leakage. Third, we introduced homology-controlled evaluation across multiple sequence-identity thresholds to determine whether model performance remained stable after restricting close homologs between training and test proteins. Finally, because isoform-level function prediction ultimately requires resolving local remodeling among closely related proteins, we defined an isoform-family directional remodeling task based on reference-relative Pfam gain and loss.

These evaluation regimes separate general protein annotation performance from the more specific question of whether the model resolves isoform-level remodeling. In this framework, strong performance under standard benchmarks is necessary but not sufficient; the key question is whether the model can identify functionally meaningful local remodeling among highly related isoforms.

#### 1.3 From structure-based clustering to graph-based function inference

Protein function prediction has traditionally been formulated for canonical proteins, where one representative protein sequence is used to summarize the functional potential of a gene. This formulation is not well matched to alternative-splicing-derived isoforms, because isoforms often share extensive sequence background while differing through local truncation, exon skipping, domain gain or loss, altered structural regions or remodeled interaction surfaces. These changes may affect only a small part of the protein, but they can involve precisely the regions that determine molecular function. Therefore, isoform-level protein function prediction requires distinguishing shared functional background from local, isoform-specific functional remodeling.

This challenge motivated us to look beyond conventional sequence-based annotation transfer. Recent large-scale studies have shown that predicted protein structures can serve as an informative organizing layer for functional inference. Foldseek-based clustering of AlphaFold-scale protein structures revealed remote structural relationships across the known protein universe (Barrio-Hernandez et al., 2023). Related work used large-scale structural comparison to uncover new protein families and folds (Durairaj et al., 2023), classify domains across the AlphaFold structural space through the Encyclopedia of Domains (Lau et al., 2024), expand CATH using predicted structures (Waman et al., 2025), infer viral protein functions through structural comparison (Nomburg et al., 2024), and discover deaminase functions by structure-based protein clustering (Huang et al., 2023). These studies show that large-scale protein structure comparison can reveal functional relationships that are not fully captured by sequence similarity alone.

However, clustering is not the same as standardized protein function prediction. A structural cluster suggests that proteins may share related functions, but it does not directly provide calibrated Pfam or Gene Ontology labels for each protein. Proteins within the same structural neighborhood may still differ in domain boundaries, motif composition, cellular localization, interaction interfaces and isoform-specific truncation patterns. This uncertainty is especially important for isoforms, where the relevant functional difference may be local rather than global. Thus, hard clustering provides a useful discovery framework, but it is not sufficient for precise and traceable isoform-level function annotation.

##### 1.4 Framework implementation in 3DisoDeepPF

We therefore designed 3DisoDeepPF to extend the idea of structure-guided functional inference from clustering to graph-based prediction. Instead of assigning proteins to discrete clusters, we represent canonical proteins and isoforms as nodes in a large sequence–structure similarity graph. Edges integrate sequence similarity and structural similarity, allowing the model to preserve both close homologous relationships and more distant structural neighborhoods. This network representation is better suited to isoform-level prediction because it allows functional information to propagate across related proteins while still preserving node-specific differences.

On top of this graph, we implemented 3DisoDeepPF as a graph-based deep learning model that outputs explicit Pfam and Gene Ontology predictions. This design differs from structure clustering in three ways. First, it replaces hard cluster membership with a continuous sequence–structure similarity network. Second, it uses a supervised graph model to infer concrete functional labels rather than only suggesting cluster-level relatedness. Third, it includes evidence tracing, allowing each prediction to be linked back to supporting graph neighbors and input modalities. 3DisoDeepPF therefore extends structure-guided functional inference from cluster-level discovery to supervised, isoform-centric protein function prediction.

### 2 Supplementary Note 2 | Interpretation of 3DisoDeepPF results and implications

#### 2.1 Biological and structural basis of isoform functional remodeling

3DisoDeepPF is motivated by a biological setting in which isoforms from the same gene often share most of their sequence but may differ in regions that are directly relevant to function. In the 3DisoGalaxy breast cancer atlas, transcript variants were progressively filtered through transcript support, ORF prediction, translational evidence and structure-quality control, yielding a structure-resolved isoform resource for breast cancer analysis (Jiang et al., 2026).

This atlas-level view provides proxy evidence that isoform remodeling is not restricted to transcript architecture. In **Extended Data Fig. 2**, many genes contribute multiple protein isoforms, and a subset of non-reference isoforms show changes in Pfam domain composition relative to the reference isoform. The distribution of ΔPfam counts suggests that most changes are local rather than complete rewiring of the protein. This is important for protein function prediction because local domain loss, gain or reconfiguration can affect molecular function even when the overall sequence remains highly similar to the reference protein.

The localization analyses provide a complementary view. Predicted membrane-associated and subcellular localization changes indicate that some isoforms may differ not only in domain content but also in cellular context. Such differences are biologically plausible because alternative exons can alter protein interaction regions and reshape protein-interaction networks (Ellis et al., 2012). In this sense, Pfam remodeling and inferred localization remodeling provide complementary signals that protein isoforms may carry functionally relevant differences.

These observations do not constitute experimental proof that every remodeled isoform has a distinct function. Rather, they define the biological problem that 3DisoDeepPF is designed to address: isoform-level function prediction must identify local functional differences against a largely shared gene-level background. This is why the framework emphasizes Pfam and GO prediction at isoform resolution, structure-aware representation, and evaluation settings that test whether the model can resolve remodeling among closely related isoforms rather than simply recover shared canonical annotations.

#### 2.2 Canonical benchmark performance as a standard protein function prediction competence test

The canonical protein benchmark was designed as a competence test rather than the main endpoint of 3DisoDeepPF. Because the central goal of this study is isoform-resolution function prediction, it was first necessary to establish that the framework performs reliably under standard protein function prediction settings. For this reason, we evaluated 3DisoDeepPF on a CAFA-aligned canonical protein benchmark constructed from Swiss-Prot, GOA and structural resources, with Pfam and the three Gene Ontology branches evaluated as parallel prediction tasks (**Fig. 1a,b**).

The canonical benchmark results show that the sequence–structure graph framework is competitive in conventional protein annotation. 3DisoDeepPF achieved competitive $F_{\max}$ scores across GO-MF, GO-BP, GO-CC and Pfam, indicating that its thresholded predictions remained effective across ontology-based and domain-based labels (**Fig. 1c**). The AUPR results provide a complementary view by measuring ranking quality across imbalanced label spaces, where many GO and Pfam labels have sparse positive examples (**Fig. 1d**). The micro-averaged precision–recall curves further showed stable ranking behavior across the canonical GO and Pfam prediction tasks, complementing the threshold-based $F_{\max}$ results (**Supplementary Fig. 1a**).

These results should not be interpreted as the main claim that 3DisoDeepPF is simply another canonical protein function predictor. Instead, they establish a necessary baseline: the framework preserves standard PFP performance while adding sequence–structure graph representation, isoform-aware evaluation and evidence tracing. These results establish the canonical benchmark as a competence test before evaluating 3DisoDeepPF in the isoform-resolution setting, where the central question is whether the model can resolve functional differences among closely related isoforms.

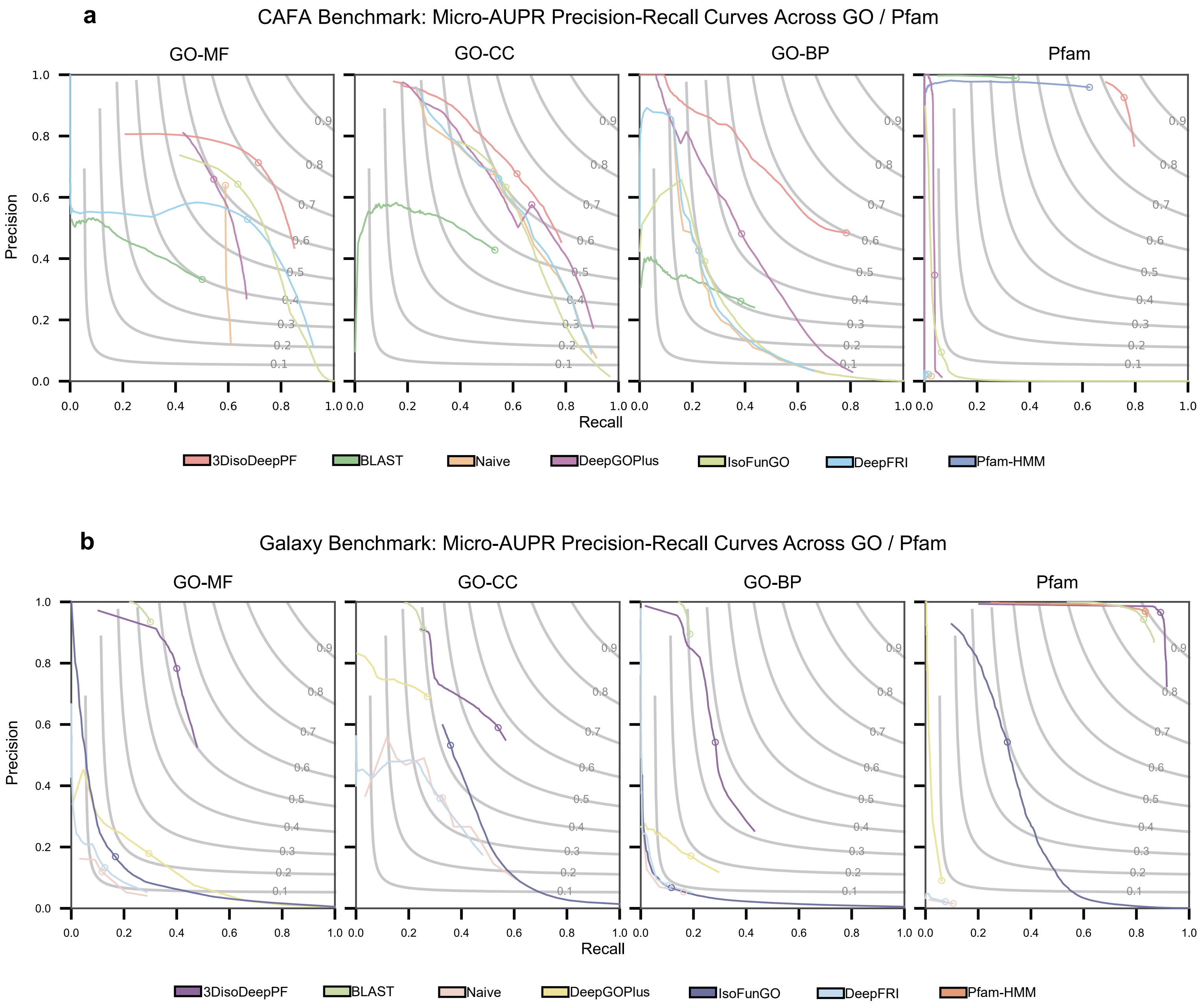

**Supplementary Fig. 1 | Micro-averaged precision–recall performance across canonical and isoform-resolved benchmarks. a,** Micro-averaged precision–recall curves on the CAFA-aligned canonical protein benchmark across Gene Ontology molecular function (GO-MF), cellular component (GO-CC), biological process (GO-BP) and Pfam prediction tasks. **b,** Micro-averaged precision–recall curves on the 3DisoGalaxy isoform-resolved benchmark across the same prediction tasks. Grey contours indicate iso-Fmax levels. Curves summarize within-benchmark ranking behavior and are not intended as a direct comparison of intrinsic task difficulty between the canonical and isoform-resolved settings.

#### 2.3 Isoform-resolution performance in the breast cancer isoform atlas

The breast cancer isoform atlas provided the main setting for testing whether 3DisoDeepPF could move beyond canonical protein annotation and operate at isoform resolution. Unlike the canonical benchmark, this setting is enriched for closely related isoforms from the same genes, where many proteins share most of their sequence but may differ in local domains, motifs or structural regions. This makes it a more stringent and biologically relevant test of whether a method can distinguish shared gene-level function from isoform-specific remodeling.

The sequence–structure graph provided an initial view of how the model organizes this isoform space. In the graph embedding, isoforms with related Pfam annotations formed local neighborhoods, suggesting that the integrated graph captures biologically meaningful structure–function organization rather than only sequence proximity (**Fig. 2a**). This observation should be interpreted as qualitative support for graph coherence, not as an independent accuracy metric. Its main value is to show that the 3DisoDeepPF graph preserves local functional organization relevant to downstream prediction.

The quantitative benchmark results further support the utility of this representation. Across the breast cancer isoform atlas, 3DisoDeepPF showed consistent advantages in Pfam, GO molecular function and GO biological process prediction, while GO cellular component showed a more metric-dependent pattern (**Fig. 2b,c**). The micro-averaged precision–recall curves provided a complementary ranking-based view of these isoform-resolved predictions across Pfam and Gene Ontology tasks (**Supplementary Fig. 1b**). These curves should be interpreted within the 3DisoGalaxy benchmark, rather than as evidence that the isoform-resolved task is directly comparable in difficulty to the canonical benchmark. Pfam and GO molecular function are more directly linked to local sequence and structural features, whereas cellular component labels may depend more strongly on cellular context, expression state and annotation granularity. Thus, the isoform-level benchmark suggests that the sequence–structure graph is most informative for functional categories that are closely coupled to domain organization and molecular activity.

The most important tests are the homology-controlled and directional remodeling analyses. Isoforms from the same gene are often highly homologous, so strong performance can be misleading if a model mainly transfers labels from close relatives or reference proteins. Under maximum sequence-identity thresholds from 30% to 95%,

3DisoDeepPF maintained robust Pfam performance, indicating that its predictions were not solely driven by unrestricted close-homology transfer (**Fig. 2d**). More specifically, the directional Pfam remodeling task evaluated whether the model could identify reference-relative domain gain and loss among isoforms from the same gene. 3DisoDeepPF remained competitive across isoform-to-reference sequence-identity bins, supporting its ability to capture remodeling signals across different levels of isoform divergence (**Fig. 2e**).

The support landscape also helps interpret this task. Domain loss events were more frequent than domain gains across the curated isoform pairs, which is consistent with the idea that many alternative isoforms arise through partial truncation, exon exclusion or local domain disruption rather than acquisition of entirely new domain architectures (**Fig. 2f**). This does not imply that domain loss is universally more important than gain; rather, it indicates that, in this atlas, the loss side provides denser annotation support and a more stable basis for evaluation.

These results indicate that 3DisoDeepPF retains predictive utility in both the canonical benchmark and the intended isoform-resolution setting. The breast cancer atlas results support three linked conclusions: the sequence–structure graph contains biologically coherent functional neighborhoods, the model can use this representation for Pfam and Gene Ontology prediction at isoform resolution, and the strongest evidence for isoform-specific resolution comes from homology-controlled Pfam prediction and reference-relative domain remodeling.

#### 2.4 Evidence tracing and interpretability of individual predictions

For isoform-resolution protein function prediction, a useful prediction should be inspectable rather than reported only as a probability score. This is especially important when predicted labels are used to prioritize disease-associated isoforms for downstream biological analysis. We therefore included an evidence-tracing utility in 3DisoDeepPF to connect individual predictions to their supporting graph context and, when available, modality-level feature support.

The evidence-tracing framework decomposes prediction support into two components. First, topological support is obtained from graph-associated proteins or isoforms that contribute most strongly to a predicted label. Second, node-intrinsic support summarizes the contribution of gated multimodal features to the prediction. This design allows a predicted Pfam or Gene Ontology label to be examined through its supporting neighborhood rather than treated as a black-box output (**Extended Data Fig. 5a**).

We used CIB1 as an illustrative case because its curated and predicted annotations provide a biologically interpretable calcium-related example. 3DisoDeepPF recovered the curated CIB1 Pfam annotation PF13499 with a high prediction score, and its predicted GO molecular function labels matched the curated GO-MF annotations with high confidence (**Supplementary Table 6**). At the Pfam/domain level, evidence tracing linked the EF-hand-related prediction to calcium-sensor-like neighbors in the graph (**Fig. 2g**). At the GO-MF level, traced neighbors were also consistent with a membrane-proximal calcium-signaling context involving integrin-associated, calcineurin-related and calcium-sensor-related proteins (**Extended Data Fig. 5b,c**).

This case illustrates how evidence tracing can make 3DisoDeepPF predictions more biologically reviewable. The traced proteins are not meant to prove that a predicted function is active in a specific cancer context, nor do they establish causal mechanisms. Instead, they provide an inspection layer: users can examine which neighboring proteins, isoforms and feature signals support a predicted label.

To support broader inspection beyond the CIB1 example, we provide the full evidence-tracing outputs in **Supplementary Data 2**. For each traceable test-set protein, one file reports all traceable predicted labels for that protein. For each label, up to the top 20 supporting proteins are listed according to the evidence-tracing score. Some labels contain fewer than 20 supporting proteins because fewer eligible neighbors passed the tracing criteria. This resource allows users to review the supporting evidence for individual predictions across the test set, rather than relying only on aggregate performance metrics.

#### 2.5 Why graph-centered multimodal integration improves isoform-resolution prediction

The ablation analyses help explain why 3DisoDeepPF performs well in the isoform-resolved setting. Among the tested components, removal of the sequence–structure similarity graph produced the largest performance loss across Pfam and Gene Ontology prediction tasks (**Extended Data Fig. 3a–c**). This indicates that the graph is not only an implementation detail, but a central source of predictive information. By connecting proteins and isoforms through sequence and structural similarity, the graph provides contextual evidence that helps the model infer functional labels beyond isolated node features.

Sequence and structure representations also contributed to performance. Removing either modality reduced prediction accuracy across most tasks, supporting the view that these two information sources are complementary rather than redundant (**Extended Data Fig. 3b,c**). Sequence embeddings capture evolutionary and compositional signals, whereas structural representations can retain information about fold similarity, domain organization and local structural context. This complementarity is particularly relevant for isoform-level prediction, where closely related isoforms may share most of their sequence but differ in structurally or functionally important regions.

Single-component analyses further showed that the graph carried the strongest isolated signal among the tested feature groups, whereas Pfam-only and motif-only models captured only part of the predictive information (**Extended Data Fig. 3d**). These results suggest that no single modality is sufficient to explain the full performance of 3DisoDeepPF across GO molecular function, GO biological process, GO cellular component and Pfam prediction.

**Supplementary Table 1. Single-component ablation analysis across GO and Pfam prediction tasks.**

| Metric | 3DisoDeepPF | Graph only | Pfam only | Motif only |
|---|---|---|---|---|
| GO-BP $F_{max}$ | 0.372 | 0.145 | 0.173 | 0.096 |
| GO-BP AUPR | 0.282 | 0.092 | 0.137 | 0.058 |
| GO-MF $F_{max}$ | 0.530 | 0.210 | 0.235 | 0.122 |
| GO-MF AUPR | 0.323 | 0.195 | 0.126 | 0.057 |
| GO-CC $F_{max}$ | 0.563 | 0.185 | 0.231 | 0.126 |
| GO-CC AUPR | 0.226 | 0.084 | 0.073 | 0.037 |
| Pfam $F_{max}$ | 0.927 | 0.412 | 0.338 | 0.229 |
| Pfam AUPR | 0.701 | 0.385 | 0.265 | 0.132 |

The ablation results support a graph-centered multimodal design for isoform-resolution protein function prediction. The graph provides the main contextual scaffold, sequence and structure contribute complementary biological signals, and auxiliary domain or motif features refine prediction. This interpretation is consistent with the broader goal of the framework, which is to resolve functional labels for closely related isoforms using both local molecular features and their position within a larger protein similarity network.

##### 2.6 Relationship to existing protein function prediction methods

Existing protein function prediction methods provide important reference points for evaluating 3DisoDeepPF. Classical approaches such as BLAST-based transfer and CAFA-style Naive predictors remain widely used because they capture two strong sources of signal: sequence homology and label frequency (Radivojac et al., 2013; Jiang et al., 2016). These methods remain useful because they are simple, interpretable and closely aligned with established annotation practice.

Recent deep-learning approaches have expanded canonical protein function prediction by learning sequence-, ontology-, network- or structure-informed representations. Sequence-based and ontology-aware methods, including DeepGO, DeepGOPlus, GOLabeler, NetGO, NetGO 2.0 and TALE, use sequence features, label structure, domain information, text features or interaction networks to improve GO prediction beyond simple homology transfer (Kulmanov et al., 2018; Kulmanov and Hoehndorf, 2020; You et al., 2018; You et al., 2019; Yao et al., 2021; Cao and Shen, 2021). Structure-aware methods, including DeepFRI, DeepGraphGO, TransFun, Struct2GO and PANDA-3D, further incorporate structural graphs, AlphaFold-derived models, protein language model embeddings or equivariant graph representations to improve function prediction from structural context (Gligorijević et al., 2021; You et al., 2021; Boadu et al., 2023; Jiao et al., 2023; Zhao et al., 2024). These approaches define the main methodological landscape for canonical protein function prediction.

However, most existing protein function prediction frameworks are developed and evaluated primarily at the canonical-protein level. This setting is not equivalent to isoform-resolution prediction, where multiple protein products from the same gene may share extensive sequence similarity but differ in domains, motifs, local structural context or interaction interfaces. In this setting, a method must separate shared gene-level function from isoform-specific functional remodeling. Isoform-oriented methods, including IsoFun and IsofunGO, move closer to this problem by explicitly considering alternative-splicing-derived isoforms and by using heterogeneous networks or GO embeddings to distribute functional information across isoforms (Yu et al., 2020; Qiu et al., 2022). Nevertheless, standardized isoform-level benchmarks remain limited, and sparse isoform-specific annotations make it difficult to evaluate fine-grained functional differences between reference and non-reference isoforms.

3DisoDeepPF was developed to address this gap rather than to replace existing protein function prediction methods. Its contribution is to formulate protein function prediction at isoform resolution and to combine sequence–structure graph representation, Pfam and Gene Ontology prediction, homology-aware evaluation, directional isoform remodeling analysis and evidence tracing within one framework. The comparative methods used in this study therefore serve as representative anchors for major annotation strategies, including homology transfer, label-frequency priors, sequence-based deep learning, structure-aware prediction and isoform-aware modeling. Detailed implementation and harmonized benchmarking of these baselines are described in **Supplementary Note 6**.

### 3 Supplementary Note 3 | Structure-aligned knowledge bases and functional proxy annotation

#### 3.1 Benchmark construction and structure-support quality of CAFA-aligned canonical dataset

To establish a standard canonical-protein reference for evaluating 3DisoDeepPF, we curated a CAFA-aligned benchmark from Swiss-Prot, GOA and structure-supported protein resources. The benchmark retained 82,879 proteins with curated Gene Ontology annotations after evidence-code filtering and ontology propagation. Structural information was assembled by integrating experimentally determined PDB structures with AlphaFoldDB models. PDB chains were mapped to UniProt accessions using SIFTS, multi-chain entries were resolved at the chain level, and representative experimental structures were selected according to experimental method, resolution and chain coverage. For proteins lacking experimental structural coverage, we incorporated computationally predicted models from AlphaFoldDB, using a predicted local distance difference test (pLDDT)-based filter to define structure-supported protein sets. We incorporated Pfam and short linear motif (SLiM) annotations as auxiliary functional features.

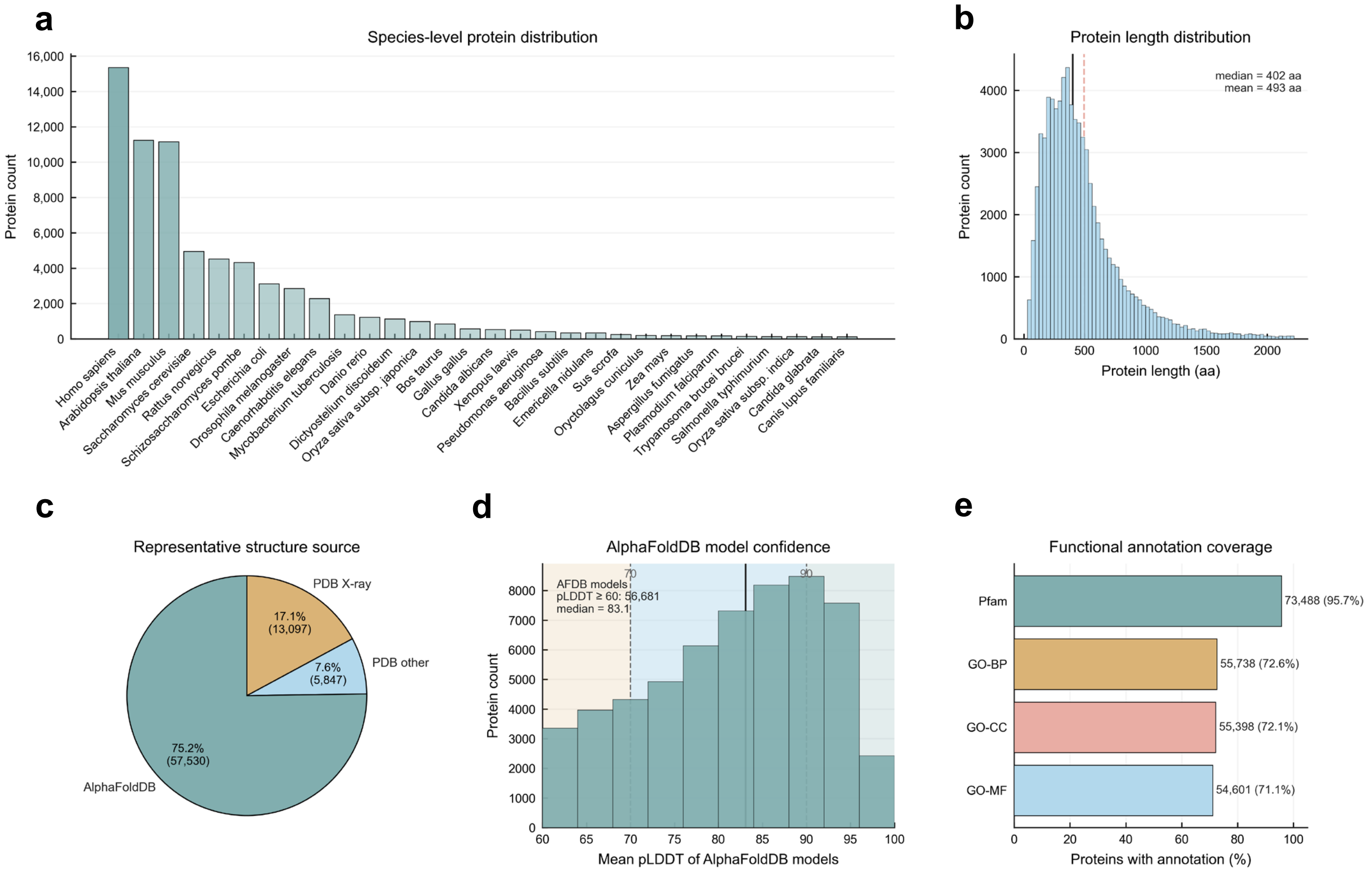

**Supplementary Fig. 2 | Overview of the CAFA-aligned canonical protein benchmark.** This figure summarizes the taxonomic composition, protein length distribution, structure-source coverage, AlphaFoldDB model confidence and functional-annotation coverage of the canonical protein benchmark used for standard protein function prediction evaluation. **a,** Species-level distribution of benchmark proteins, showing broad taxonomic coverage with human proteins as the largest single species group. **b,** Protein length distribution across benchmark proteins; dashed lines indicate the median length of 402 amino acids and the mean length of 493 amino acids. **c,** Structure-source assignment after representative structure selection, including AlphaFoldDB models, PDB X-ray structures and other PDB experimental structures. **d,** Distribution of mean pLDDT scores for AlphaFoldDB-derived models with available confidence scores; dashed lines indicate the retained confidence threshold and the median pLDDT. **e,** Coverage of curated GO annotations and Pfam domain annotations, including GO-MF, GO-BP, GO-CC and Pfam. Together, these summaries show that the benchmark spans diverse species, typical protein-length ranges, broad structural coverage and high functional-label coverage for CAFA-aligned evaluation.

**Supplementary Fig. 2** summarizes the resulting canonical benchmark. The dataset spans a broad taxonomic range, with human proteins forming the largest species group (**Supplementary Fig. 2a**). Protein lengths are broadly distributed, with a median length of 402 amino acids and a mean length of 493 amino acids (**Supplementary Fig. 2b**). Most structure assignments were derived from AlphaFoldDB models, complemented by PDB X-ray and other experimental structures (**Supplementary Fig. 2c**). The AlphaFoldDB-derived subset demonstrated generally high model confidence, with a median mean pLDDT of 83.1 (**Supplementary Fig. 2d**). Known annotation coverage was high across Pfam and the three Gene Ontology branches, supporting the use of this benchmark for standard canonical protein function prediction before extending the framework to isoform-resolution analyses (**Supplementary Fig. 2e**).

##### 3.2 Composition and annotation coverage of the 3DisoGalaxy structure atlas

We next summarized the composition and annotation coverage of the 3DisoGalaxy structure atlas to assess its suitability for isoform-resolution function prediction. The atlas contains 46,601 protein entries, including non-canonical isoforms, canonical proteins and human proteome reference proteins, with broad isoform diversity across genes, including genes with more than ten isoforms (**Supplementary Fig. 3a,b**). Of these, 46,411 structure-supported entries with available sequences were used for downstream Pfam and localization annotation. Protein lengths were broadly distributed, with a median length of 464 amino acids, and predicted structures showed generally high confidence, with a median mean pLDDT of 81.8 (**Supplementary Fig. 3c,d**). These properties support the use of the atlas for structure-based similarity calculation and graph construction. Further details on the construction and biological characterization of 3DisoGalaxy are provided in the companion preprint (Jiang et al., 2026).

The atlas also provided sufficient functional and localization information for model evaluation and downstream inference. Predicted localization labels covered major cellular compartments, suggesting that isoform-level remodeling may involve cellular context as well as domain composition (**Supplementary Fig. 3e**). Functional-label coverage was approximately half of the atlas for each label space, with 44.2% coverage for GO-MF, 49.6% for GO-CC, 47.1% for GO-BP and 49.7% for Pfam (**Supplementary Fig. 3f**). These coverage levels provided enough labeled isoforms for supervised evaluation while retaining a substantial unlabeled fraction for inference. Together, these summaries indicate that 3DisoGalaxy provides an isoform-rich,

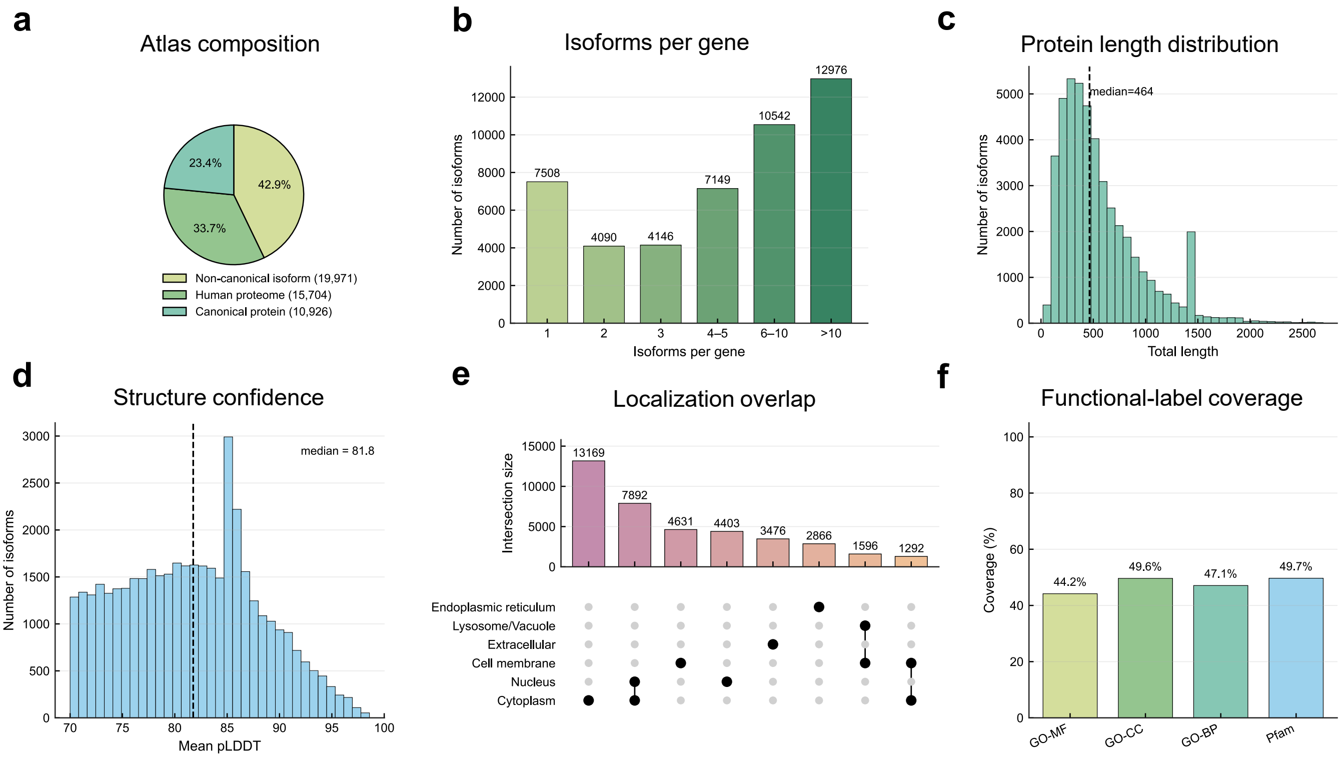

**Supplementary Fig. 3 | Composition, structural quality and annotation coverage of the breast cancer isoform atlas. a,** Composition of protein entries in the atlas, grouped as non-canonical isoforms, canonical proteins and human proteome reference proteins. **b,** Distribution of isoform family sizes across genes, showing the number of isoforms assigned to each gene-level bin. **c,** Distribution of protein sequence lengths across atlas entries; dashed line denotes the median length. **d,** Distribution of mean pLDDT values for retained structural models; dashed line denotes the median mean pLDDT. **e,** UpSet plot summarizing predicted subcellular localization categories across isoforms. Bars show intersection sizes, and connected filled dots indicate the localization categories included in each intersection. **f,** Functional-label coverage across GO molecular function, GO cellular component, GO biological process and Pfam, measured as the percentage of isoforms with at least one label in each category.

structure-supported and partially annotated resource for developing and evaluating 3DisoDeepPF.

##### 3.3 Pfam and localization prediction with existing tools

Pfam domains were annotated using profile hidden Markov models from the Pfam-A database (version 35.0) with HMMER (hmmscan, version 3.4). Protein sequences from 46,411 structure-supported entries were scanned against the Pfam-A HMM library using curated trusted cutoffs to ensure high-confidence domain assignments. Domain-level hits were extracted from HMMER domain tables, and redundant hits for the same Pfam domain within a protein were resolved by retaining the highest-scoring match. The resulting Pfam annotations were used for isoform-level domain presence and reference-relative gain and loss analyses, and subsequently mapped onto protein structures for structural interpretation.

Subcellular localization of protein isoforms was annotated using DeepLoc 2.0 based on amino-acid sequences. The same set of 46,411 protein sequences was used to ensure consistency with Pfam domain annotation for downstream functional inference.

#### 4 Supplementary Note 4 | Dataset construction, partitioning and leakage-control design

To ensure that model evaluation was not driven by inappropriate data partitioning or target-label leakage, we documented the split strategy, temporal-split diagnostics and task-specific masking rules used in 3DisoDeepPF. The same data-cleaning and label-processing principles were applied to both the CAFA-aligned canonical benchmark and the 3DisoGalaxy breast cancer isoform dataset (Jiang et al., 2026). However, because the 3DisoGalaxy dataset is a fixed breast-cancer isoform atlas derived from human tumor and adjacent-normal samples, its GO labels are inherited from external annotation resources rather than generated through a temporally suitable isoform-specific annotation trajectory. We therefore used a stringent random split for this isoform-level setting, while retaining the same cleaning, masking and leakage-control procedures wherever applicable.

#### 4.1 CAFA-aligned canonical benchmark construction and partitions

##### 4.1.1 Raw dataset collection and protein inclusion

We assembled a conservative, CAFA-aligned benchmark to enable fair evaluation and comparison on a public, externally curated dataset. Protein sequences were obtained from UniProtKB/Swiss-Prot (release 2025_04; downloaded September 1, 2025), yielding 573,661 reviewed protein entries (The UniProt Consortium, 2025). Functional annotations were retrieved from the EBI Gene Ontology Annotation (GOA) database on the same date, from which 1,431,969,797 GOA annotation records were scanned (Huntley et al., 2015). The ontology structure was defined by the Gene Ontology basic file (go-basic.obo, version 2025-07-22), comprising 43,230 GO terms (The Gene Ontology Consortium, 2026).

Proteins were retained if they had a Swiss-Prot sequence and at least one MF, BP, or CC GO label remaining after the conservative evidence filtering described above and ontology propagation. This filtering yielded a curated GO-annotated table of 82,879 proteins before structural-coverage harmonization and final benchmark filtering.

##### 4.1.2 Structural coverage and representative selection

To support structure-aware function prediction in the canonical benchmark, we assembled a structural compendium by combining experimentally resolved PDB structures with AlphaFold DB models (Berman et al., 2000; Varadi et al., 2022). PDB structures were downloaded from the wwPDB and mapped at the chain level to canonical UniProt accessions using PDBx/mmCIF metadata and UniProt cross-references, following the principles of SIFTS-based residue- and accession-level mapping (Berman et al., 2007; Velankar et al., 2013). Multi-chain entries were decomposed into chain-only mmCIF files, allowing each structural chain to be linked to a single UniProt accession. This produced 84,800 chain-level experimental structures covering 29,392 UniProt accessions.

Because many proteins were associated with multiple experimental structures, we selected one representative structure per accession to reduce redundancy while preserving structural quality. Candidate structures were ranked using a lexicographic quality score that prioritized experimental method, nominal resolution, X-ray refinement quality when available, and chain coverage. Specifically, X-ray diffraction was prioritized over electron microscopy, NMR and other methods; lower resolution, lower Rfree/Rwork and greater chain residue coverage were preferred, with missing values penalized. This procedure selected one representative experimental structure for each UniProt accession with available PDB coverage.

For proteins without PDB coverage, structures were supplemented from AlphaFold DB as secondary structural evidence. We used a tiered confidence policy: models with mean pLDDT ≥ 70 were retained as high-confidence supplements (47,272 structures), models with 60 ≤ pLDDT < 70 were kept only for coarse-coverage and sensitivity analyses (16,928 structures), and models with pLDDT < 60 were excluded. Thus, the primary structural compendium remained anchored on PDB representatives and high-confidence AlphaFold models, while lower-confidence models were reserved for robustness checks.

###### 4.1.3 Ontology-specific split statistics

The CAFA-aligned temporal split strategy is described in the main **Methods**. In brief, we followed the time-delayed no-knowledge design, in which training labels were restricted to annotations visible before the historical cutoff and test labels were defined from annotations that appeared within the later evaluation window. In the supplementary analysis, we report the resulting split sizes to make the benchmark construction transparent. Because the split was determined by GOA timestamps and ontology-specific label availability, rather than by enforcing a fixed random ratio, the final train, validation and test proportions differed across MF, BP and CC. The resulting protein counts for each split are summarized in **Supplementary Table 2**.

**Supplementary Table 2. GO ontology-specific train, validation and test split summary of CAFA-aligned dataset.**

| GO aspect | Train proteins | Validation proteins | Test proteins | Total proteins | Train (%) | Validation (%) | Test (%) |
|---|---|---|---|---|---|---|---|
| Molecular function | 37,552 | 2,398 | 6,808 | 46,758 | 80.31 | 5.13 | 14.56 |
| Biological process | 49,100 | 1,715 | 1,822 | 52,637 | 93.28 | 3.26 | 3.46 |
| Cellular component | 49,898 | 1,341 | 1,413 | 52,652 | 94.77 | 2.55 | 2.68 |

###### 4.1.4 Overview of a multi-species, structure-annotated CAFA-aligned benchmark

After structural-coverage harmonization and final benchmark filtering, the CAFA-aligned canonical benchmark integrated 76,804 unique proteins, exceeding the scale of the recent CAFA dataset of 59,397 proteins (Wang et al., 2025). In addition to expanding protein coverage, we incorporated representative PDB and AlphaFold DB structures under explicit quality criteria, enabling evaluation on a structurally characterized benchmark for protein function prediction.

The dataset spans 1,648 species-level organism labels, consistent with CAFA's multi-species design. The top 30 species-level groups account for 69,643 proteins, representing 90.7% of the benchmark, with *Homo sapiens* forming the largest single species group (15,348 proteins; 20.0%) (**Supplementary Fig. 2a**). Protein lengths covered a broad but typical range, with a median length of 402 amino acids and a mean length of 493 amino acids (**Supplementary Fig. 2b**).

Most structure-supported proteins were represented by AlphaFold DB models (57,530 proteins; 74.9%), while 18,944 proteins were supported by PDB structures, including 13,097 X-ray structures and 5,847 other experimental structures; 330 proteins had no representative structure in this table (**Supplementary Fig. 2c**). Among AlphaFold DB representatives with mean pLDDT $\geq$ 60, the median mean pLDDT was 83.1, indicating generally high model confidence among retained predicted structures (**Supplementary Fig. 2d**). Functional-feature coverage was also broad, with Pfam annotations for 73,488 proteins (95.7%) and propagated GO labels for 54,601 MF proteins (71.1%), 55,738 BP proteins (72.6%) and 55,398 CC proteins (72.1%) (**Supplementary Fig. 2e**). These summaries define the CAFA-aligned benchmark as a large, multi-species and structurally annotated protein set for canonical protein function prediction evaluation.

##### 4.2 Overview of the 3DisoGalaxy isoform atlas

In addition to the CAFA-aligned canonical benchmark, we evaluated 3DisoDeepPF on the 3DisoGalaxy breast cancer isoform atlas. Unlike the canonical CAFA-style dataset, which is built from public Swiss-Prot proteins and GOA annotations across species, the 3DisoGalaxy dataset is a human, cancer-focused isoform resource derived from transcriptomic, translational and structural reconstruction (Jiang et al., 2026). This setting was designed to test whether the model can operate in an isoform-resolved context, where multiple protein isoforms from the same gene may share extensive sequence similarity but differ in domains, motifs, structural regions or functional annotations.

##### 4.2.1 Why temporal no-knowledge splitting was not applied to 3DisoGalaxy

We next tested whether the CAFA-style temporal split used for the canonical benchmark could be transferred to the 3DisoGalaxy isoform benchmark. We performed a systematic scan of GOA annotation timestamps for atlas-associated proteins and evaluated candidate training visibility cutoffs, $t_0$. This scan revealed a sharp deposition boundary around 2003-09-04: earlier candidate $t_0$ values retained too few pre-$t_0$ labels for training, whereas later cutoffs already included most atlas-associated annotations.

Using 2003-09-04 as a diagnostic candidate $t_0$, most post-$t_0$ GO terms were absent from the pre-$t_0$ label space, including 90.63% of MF terms, 89.18% of CC terms and 91.98% of BP terms (**Supplementary Table 3**). Thus, a TDNK split in this dataset would mainly reflect a database deposition artifact and severe label-space shift, rather than a meaningful no-knowledge evaluation. We therefore used a strict snapshot-based split for the 3DisoGalaxy benchmark after final atlas construction, with ontology-specific task definitions and leakage-control rules.

**Supplementary Table 3. GO label-space shift under a diagnostic CAFA-style temporal split for the 3DisoGalaxy isoform benchmark.**

| Ontology | GO terms before $t_0$ | GO terms after $t_0$ | Proportion of after-$t_0$ GO terms absent before $t_0$ |
|---|---|---|---|
| GO-MF | 392 | 3,908 | 90.63% |
| GO-CC | 164 | 1,516 | 89.18% |
| GO-BP | 813 | 9,945 | 91.98% |

##### 4.2.2 Fixed-snapshot, ontology-specific split strategy for the 3DisoGalaxy isoform benchmark

Because the 3DisoGalaxy benchmark is a curated, snapshot-based breast cancer isoform atlas, we used a snapshot-based split after the final eligible isoform set, structures and functional labels had been defined under the same curation rules. This strategy avoids mixing different annotation-generation stages and provides a consistent evaluation setting within the curated atlas. MF, BP, CC and Pfam were treated as separate prediction tasks because their labeled isoform sets are not identical. For each task, labeled isoforms were randomly divided into training, validation and test sets using an approximately 80/10/10 split. This design was used for isoform-resolution evaluation within the 3DisoGalaxy isoform universe, whereas the CAFA-aligned temporal split described in the main Methods was used for the canonical benchmark.

The resulting split sizes were highly consistent across tasks, with 80.00–80.04% of labeled isoforms assigned to training, 9.99–10.00% to validation and 9.98–10.00% to testing (**Supplementary Table 4**). The final labeled isoform sets contained 20,577 MF-labeled isoforms, 21,942 BP-labeled isoforms, 23,130 CC-labeled isoforms and 23,147 Pfam-labeled isoforms. Reporting these task-specific split statistics makes the benchmark construction transparent and distinguishes the snapshot-based 3DisoGalaxy evaluation from the CAFA-style temporal no-knowledge evaluation.

**Supplementary Table 4: Ontology-specific fixed-snapshot splits used for 3DisoGalaxy architecture validation.**

| Ontology | Train | Validation | Test | Total labeled isoforms |
|---|---|---|---|---|
| GO-MF | 16,469 (80.04%) | 2,055 (9.99%) | 2,053 (9.98%) | 20,577 |
| GO-BP | 17,557 (80.02%) | 2,194 (10.00%) | 2,191 (9.99%) | 21,942 |
| GO-CC | 18,504 (80.00%) | 2,313 (10.00%) | 2,313 (10.00%) | 23,130 |
| Pfam | 18,523 (80.02%) | 2,313 (9.99%) | 2,311 (9.98%) | 23,147 |

##### 4.2.3 Dataset scope and composition of the 3DisoGalaxy isoform benchmark

The final atlas-level benchmark contains 46,601 protein entries, including 19,971 non-canonical isoforms (42.9%), 15,704 human proteome reference proteins (33.7%) and 10,926 canonical proteins (23.4%) (**Supplementary Fig. 3a**). This composition makes the dataset substantially different from a canonical-only benchmark and provides a more demanding setting for isoform-level function prediction. Isoform representation varied across genes, with many genes contributing multiple isoforms and the largest isoform groups coming from genes with more than six isoforms (**Supplementary Fig. 3b**).

This dataset also covers a broad protein-length range (up to 2500 amino acids), with a median length of 464 amino acids (**Supplementary Fig. 3c**). Retained structural models showed generally high confidence, with a median mean pLDDT of 81.8, supporting the use of this dataset for structure-aware analysis (**Supplementary Fig. 3d**). Predicted subcellular localization profiles (**Supplementary Note 3**) showed that the largest isoform groups were assigned to cytoplasm alone (13,169), combined cytoplasmic–nuclear localization (7,892), cell membrane (4,631) and nucleus (4,403). Smaller but well-represented groups included extracellular (3,476), endoplasmic reticulum (2,866) and mixed membrane-associated localization patterns (**Supplementary Fig. 3e**). Functional annotation coverage was broad but incomplete, as expected for an isoform-resolved cancer atlas: 44.2% of isoforms had GO-MF labels, 49.6% had GO-CC labels, 47.1% had GO-BP labels and 49.7% had Pfam annotations (**Supplementary Fig. 3f**). Together, these summaries show that the 3DisoGalaxy atlas provides a human cancer-focused, isoform-enriched and structure-supported dataset for evaluating function prediction beyond the canonical protein setting. Detailed data sources, atlas construction procedures and stepwise analyses are described in Jiang et al. (2026), and the processed data can be accessed through the 3DisoGalaxy portal at http://3disogalaxy.com/.

#### 4.3 Benchmark hygiene, input masking and split-specific graph isolation

##### 4.3.1 Split definition before model training

The two evaluation datasets used different split strategies because they have different annotation structures. For the CAFA-aligned canonical benchmark, splits were defined by GOA timestamps under the TDNK protocol described in the main Methods. For the 3DisoGalaxy isoform benchmark, CAFA-style temporal splitting was first tested but rejected because timestamp diagnostics showed a severe label-space shift. We therefore used fixed-snapshot, ontology-specific splits after the final atlas entries and labels had been defined. In both datasets, splits were fixed before model training, hyperparameter tuning and final evaluation.

##### 4.3.2 Label masking and no-GO-input rule

For GO prediction, we used a strict no-GO-input rule (**Supplementary Table 5**). When predicting any GO branch, all GO-derived annotations were removed from model input, including GO-MF, GO-BP and GO-CC. Thus, GO labels were not allowed to leak across ontology branches. For example, GO-BP or GO-CC annotations could not be used as auxiliary features for GO-MF prediction. Only non-GO features, such as Pfam, motif, sequence-derived and structure-derived features, were allowed under the corresponding input setting. For Pfam prediction, Pfam labels were masked from input.

This rule separates the information used as model input from the labels used as prediction targets.

**Supplementary Table 5. Benchmark hygiene and task-specific leakage-control rules used across both evaluation datasets.**

| Prediction task | Labels excluded from model input | Auxiliary features allowed | Graph construction | Split-specific graph rule |
|---|---|---|---|---|
| GO-MF | All GO labels: GO-MF, GO-BP and GO-CC | Non-GO features only, including Pfam, motif and sequence/structure-derived features | Sequence and structure similarity only | Training used only the training-induced graph; Target labels for non-training partitions are strictly masked |
| GO-BP | All GO labels: GO-MF, GO-BP and GO-CC | Non-GO features only, including Pfam, motif and sequence/structure-derived features | Sequence and structure similarity only | Training used only the training-induced graph; Target labels for non-training partitions are strictly masked |
| GO-CC | All GO labels: GO-MF, GO-BP and GO-CC | Non-GO features only, including Pfam, motif and sequence/structure-derived features | Sequence and structure similarity only | Training used only the training-induced graph; Target labels for non-training partitions are strictly masked |
| Pfam | Pfam labels | Non-Pfam features under the corresponding input setting | Sequence and structure similarity only | Training used only the training-induced graph; Target labels for non-training partitions are strictly masked |

##### 4.3.3 Split-specific graph isolation

To leverage the global topology of the protein similarity space, we adopted a transductive graph learning configuration. To maintain strict evaluation hygiene, target labels for the validation, test, and unlabeled nodes were masked during model training. This design ensures that parameters are optimized exclusively on the supervised loss of the training partition, eliminating look-ahead label leakage.

### 5 Supplementary Note 5 | Graph-based function inference from integrated sequence–structure similarity

#### 5.1 Graph construction and protein similarity calculation

Protein similarities were computed separately at the sequence and structural levels before graph construction. For sequence similarity, each protein sequence was searched against the complete protein sequence collection using BLASTP from BLAST+ v2.5.0, with self-matches removed and hits retained at an E-value threshold of 0.0118. For structural similarity, predicted or experimentally resolved protein structures were compared using Foldseek v8.ef4e960 in TM-align mode (using parameters --alignment-type 1 --cluster-search 0 --tmalign-fast 1) to estimate pairwise structural similarity between protein models. Before integration, sequence- and structure-derived similarity scores were linearly rescaled to the 0–1 range when required, allowing the two similarity measures to be combined on a common scale as described in the Methods.

#### 5.2 Empirical selection of the sequence–structure similarity cutoff for graph construction

The integrated sequence and structure similarity graph requires a cutoff that removes weak edges while preserving sufficient connectivity for neighborhood-based information transfer. We therefore evaluated cutoffs ranging from 0.1 to 0.9 in both the CAFA-aligned benchmark and the 3DisoGalaxy isoform-centric evaluation set using graph topology, local annotation coherence, masked-label recovery, and downstream 3DisoDeepPF performance.

For each cutoff, we measured the fraction of retained edges, the fraction of nodes contained in the largest connected component and the mean Pfam label overlap among connected node pairs. These three quantities were rescaled to 0–1 for visualization. In both datasets, increasing the cutoff improved local Pfam coherence but progressively reduced graph connectivity, indicating the expected trade-off between annotation consistency and graph coverage (**Supplementary Fig. 4a and 5a**). Stepwise analysis between adjacent cutoffs further showed that the 0.2-to-0.3 transition provided an early gain in Pfam label overlap before the larger connectivity losses observed at more stringent cutoffs (**Supplementary Fig. 4b and 5b**).

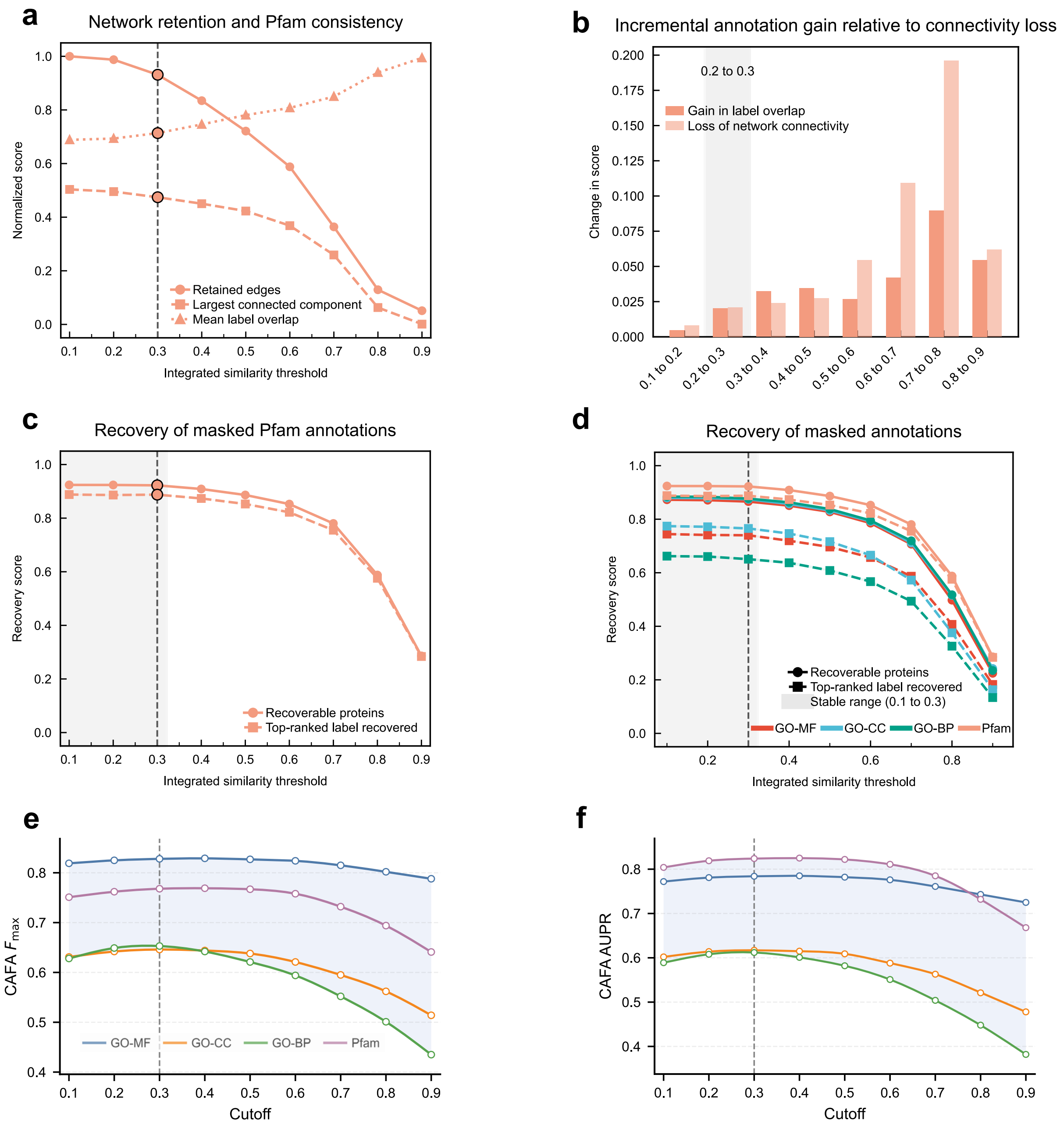

**Supplementary Fig. 4 | Cutoff selection for graph construction on the CAFA-aligned benchmark. a**, Network retention and Pfam consistency across integrated similarity thresholds. The x axis denotes the minimum integrated sequence–structure similarity required to retain an edge. The y axis shows the fraction of retained edges, the fraction of proteins in the largest connected component and the mean Pfam label overlap, rescaled to 0–1 for display. Increasing the cutoff improves local Pfam consistency but reduces graph connectivity. **b**, Stepwise change in Pfam label overlap and network connectivity between adjacent cutoffs. The x axis denotes each cutoff transition. The y axis shows the absolute change between adjacent thresholds; darker bars indicate the increase in mean Pfam label overlap, and lighter bars indicate the decrease in network connectivity. **c**, Pfam masked-label recovery across cutoffs. The x axis shows the graph cutoff. The y axis shows two recovery rates: the fraction of proteins for which at least one masked Pfam label was recovered and the fraction for which the top-ranked recovered Pfam label matched a masked label. **d**, Masked-label recovery across GO molecular function, GO biological process, GO cellular component and Pfam, with axes and line types defined as in **c**. **e,f**, 3DisoDeepPF performance on the CAFA-aligned benchmark across graph cutoffs, measured by Fmax (**e**) and AUPR (**f**). The selected cutoff of 0.3 lies within the stable recovery and performance range while preserving graph connectivity.

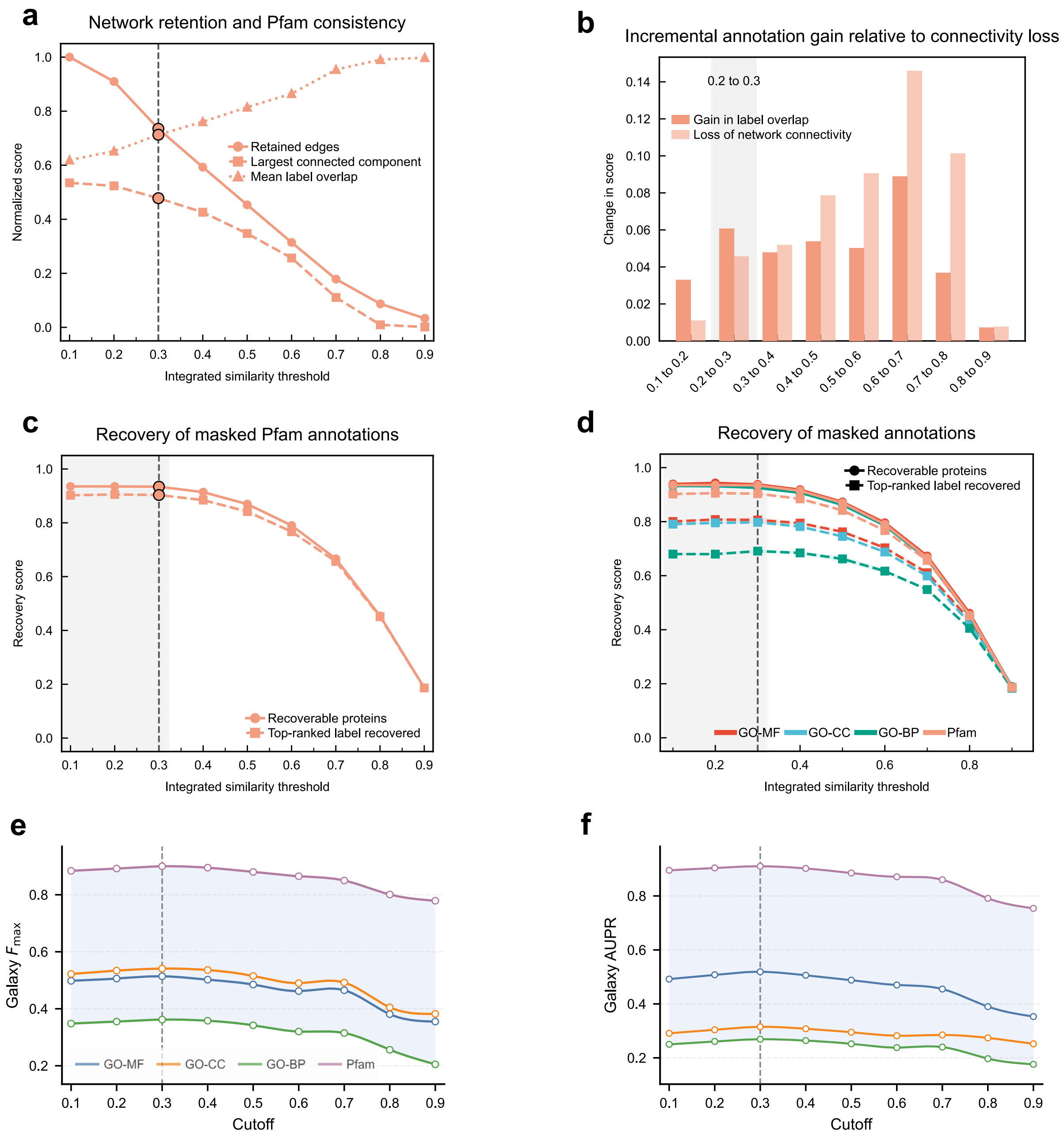

**Supplementary Fig. 5 | Cutoff selection for graph construction on the 3DisoGalaxy evaluation set. a**, Network retention and Pfam consistency across integrated similarity thresholds. The x axis denotes the minimum integrated sequence–structure similarity required to retain an edge. The y axis shows three quantities rescaled to 0–1 for display: the fraction of retained edges, the fraction of nodes in the largest connected component and the mean Pfam label overlap among connected node pairs. Increasing the cutoff improves local Pfam consistency but reduces graph connectivity. **b**, Stepwise change in Pfam label overlap and network connectivity between adjacent cutoffs. The x axis denotes each cutoff transition, and the y axis shows the absolute change between adjacent thresholds; darker bars indicate the increase in mean Pfam label overlap, and lighter bars indicate the decrease in network connectivity. **c**, Pfam masked-label recovery across cutoffs. The x axis shows the graph cutoff. The y axis shows two recovery rates: the fraction of protein isoforms for which at least one masked Pfam label was recovered and the fraction for which the top-ranked recovered Pfam label matched a masked label. **d**, Masked-label recovery across GO molecular function, GO biological process, GO cellular component and Pfam, with axes and line types defined as in **c**. **e,f**, 3DisoDeepPF performance on the 3DisoGalaxy evaluation set across graph cutoffs, measured by Fmax (**e**) and AUPR (**f**). The selected cutoff of 0.3 lies within the stable recovery range and provides a practical balance between graph connectivity, local label coherence and downstream prediction performance.

We next assessed whether graph pruning affected label recoverability. For masked-label recovery, we measured two rates: the fraction of nodes for which at least one masked label was recovered, and the fraction for which the top-ranked recovered label matched a masked label. Pfam recovery remained stable from 0.1 to 0.3 and declined at higher cutoffs in both datasets (**Supplementary Fig. 4c and 5c**). Similar recovery trends were observed across GO-MF, GO-BP, GO-CC, and Pfam, supporting the use of the same cutoff range across independent annotation spaces (**Supplementary Fig. 4d and 5d**).

Finally, we evaluated 3DisoDeepPF performance across cutoffs using $F_{\max}$ and AUPR. Performance remained stable around the selected cutoff and decreased under stronger graph pruning, especially at higher thresholds (**Supplementary Fig. 4e,f and 5e,f**). We therefore selected 0.3 as the operating cutoff for graph construction. This value should be interpreted as an empirical compromise rather than a unique optimum, balancing graph connectivity, local Pfam coherence, masked-label recoverability and downstream prediction performance in both evaluation settings.

##### 5.3 Two-dimensional visualization of the isoform similarity network

The integrated isoform similarity graph was constructed from sequence- and structure-derived protein similarities using the selected edge-retention cutoff of 0.3 (**Supplementary Fig. 6a**). Nodes represent proteins, and edges represent retained pairwise similarity relationships weighted by the integrated similarity index. For interpretation, nodes in the graph were grouped into three annotation categories: annotated canonical proteins, annotated human proteome proteins and unannotated non-canonical proteins (**Supplementary Fig. 6a**). Annotated canonical and human proteome proteins provide known Pfam and GO functional context, whereas unannotated non-canonical proteins represent isoform-derived proteins with limited or absent direct annotation. This graph design allows unannotated non-canonical proteins, as well as sparsely annotated isoform nodes, to be placed within neighborhoods of functionally characterized proteins, providing the graph context used by 3DisoDeepPF for isoform-level functional inference.

After graph construction, we visualized the topology of the integrated isoform similarity graph using Gephi (version 0.10.1) (**Supplementary Fig. 6b**). The graph was visualized with the ForceAtlas2 layout using the following settings: threads = 31; tolerance (speed) = 1.0; approximate repulsion enabled (approximation = 1.2); scaling

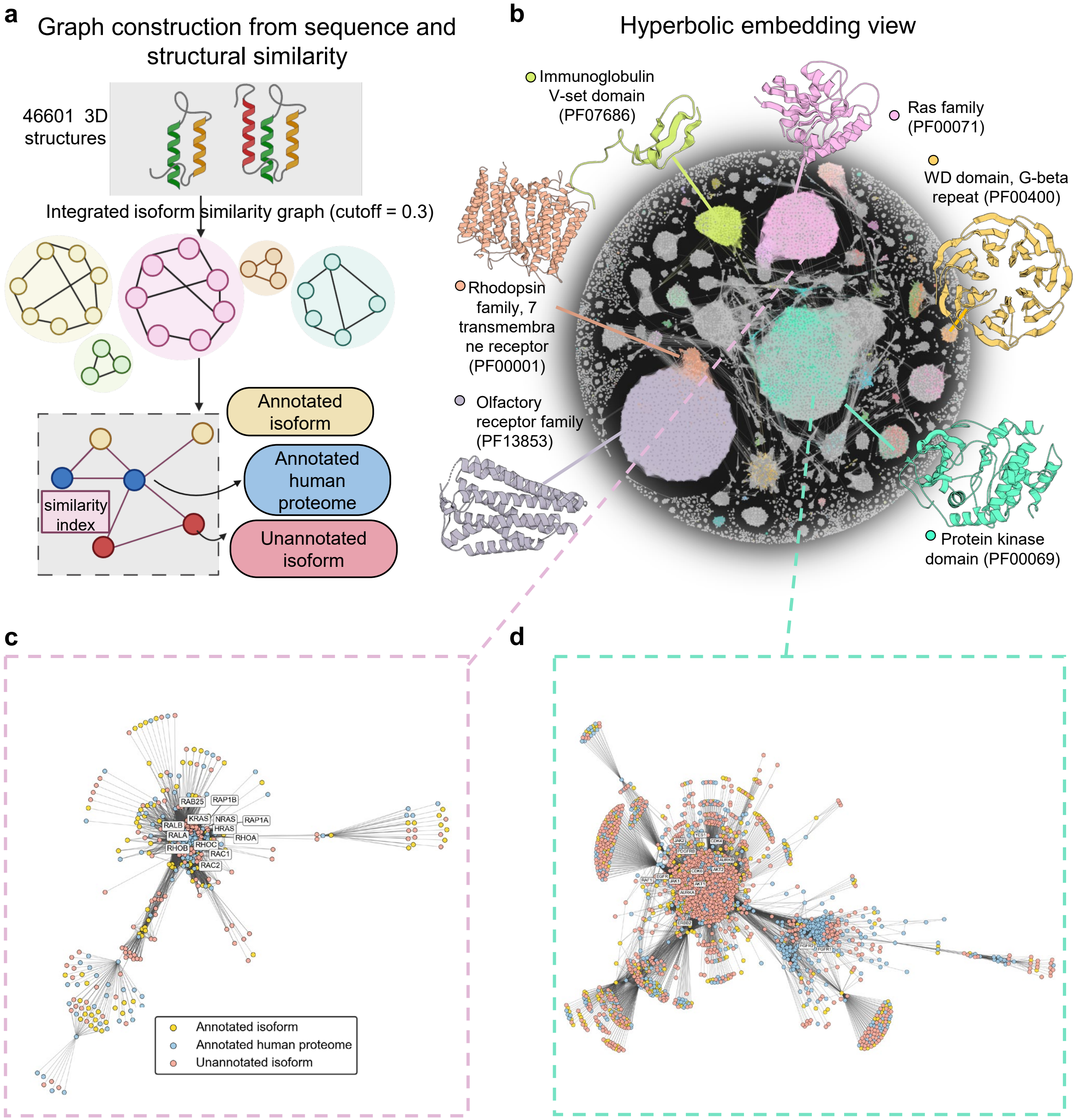

**Supplementary Fig. 6 | Integrated protein isoform similarity graph and representative local neighborhoods.** **a**, Schematic of graph construction from integrated sequence and structural similarity. Nodes are grouped into annotated canonical proteins, annotated human-proteome proteins and unannotated non-canonical protein isoforms. **b**, Global view of the resulting protein isoform similarity graph, with selected domain-enriched regions highlighted by representative structures. **c**,**d**, Expanded local neighborhoods enriched for the protein kinase domain PF00069 (**c**) and the Ras family domain PF00071 (**d**). Blue nodes indicate proteins annotated with the focal Pfam domain, whereas other nodes are colored by the protein categories defined in **a**. The 15 most frequent oncogene-associated gene symbols in each neighborhood are labelled next to their corresponding nodes to highlight disease-relevant graph regions.

= 10.0; stronger gravity enabled (gravity = 0.1); dissuade hubs enabled; LinLog mode disabled; prevent overlap enabled; edge weight influence = 1.0; normalize edge weights disabled; and inverted edge weights disabled. In the global view, domain-enriched regions were annotated using representative Pfam labels and corresponding protein structures, and proteins without Pfam annotation were shown in gray (**Supplementary Fig. 6b**).

To make local graph organization more interpretable, we expanded two large Pfam-enriched clusters dominated by the protein kinase domain (PF00069) and the Ras family domain (PF00071), respectively (**Supplementary Fig. 6c,d**). These clusters were selected by cluster size and focal-domain enrichment, rather than by requiring all nodes to carry the focal Pfam label. In each cluster, blue nodes indicate proteins annotated with the focal domain, whereas the remaining nodes follow the protein categories defined in **Supplementary Fig. 6a**. To highlight disease-relevant neighborhoods, we labeled the 15 oncogene-associated nodes with the highest graph degree in each cluster. These examples illustrate how annotated human proteome proteins and annotated isoforms can provide local graph context for unannotated isoforms. Pfam is shown as an interpretable domain-level readout, while the same graph-construction framework was used for GO-MF, GO-BP, GO-CC and Pfam prediction. Molecular structures and residue-level highlights were rendered separately in PyMOL (version 3.1.6.1), and the Gephi layout was used purely for visualization, not as a model input.

#### 5.4 Three-dimensional hyperbolic embedding of the isoform similarity network

We generated a hyperbolic visualization of the breast-cancer isoform similarity graph to reproduce the graph input used by 3DisoDeepPF under the same construction and cutoff settings and to assess whether the resulting neighborhood structure and clusters were biologically coherent. The weighted undirected graph was embedded with node2vec to obtain 64-dimensional node representations. Node2vec was run with return and in–out parameters $p = 0.5$ and $q = 2.0$.

For static visualization in the manuscript, we used a two-dimensional hyperbolic UMAP representation of the learned graph embeddings, which was used to visualize the full and held-out datasets and to inspect neighborhood-level consistency of curated UniProtKB annotations and 3DisoDeepPF predictions. For interactive exploration in the web portal, the embeddings were additionally rendered in a three-dimensional embedding view to support navigation and neighborhood inspection.

To summarize geometric coherence, we further applied k-means clustering to the embedded coordinates and inspected cluster-level coherence using Pfam and GO annotations as qualitative sanity checks. These visualizations were intended as qualitative assessments of graph organization and neighborhood coherence rather than quantitative evidence of predictive accuracy.

##### 5.5 Interactive visualization of annotation and prediction consistency

The resulting hyperbolic embedding produced a radial, branch-like organization of the integrated isoform similarity graph, in which major annotation-enriched neighborhoods could be visually inspected across the embedded protein nodes (**Supplementary Fig. 7**). To compare curated annotations with model outputs, we generated paired interactive HTML visualizations for four annotation spaces: Pfam, GO molecular function, GO biological process and GO cellular component. For each annotation space, the left panel displays curated annotations and the right panel displays 3DisoDeepPF predictions for the same protein nodes, using matched colors for shared label categories. We generated visualizations for multiple data partitions, including the full graph, training, validation, test and unlabeled node sets where applicable. For each partition, we provided both all-label views and top-50-label views, with the latter used to improve visual interpretability in densely annotated graphs.

The interactive HTML files allow users to rotate, drag, and zoom the three-dimensional embedding to inspect the spatial distribution of protein labels and local neighborhood structure. To support more flexible exploration, we also provide web-based interactive views of the four annotation spaces at http://www.3disodeeppf.com/, where users can search and inspect results by annotation label, gene name, and associated protein structure. These visualizations are provided as qualitative companions to the quantitative benchmark analyses, rather than as independent evidence of predictive accuracy. The interactive files are available from the 3DisoDeepPF GitHub repository at https://github.com/FeliciaTJiang/3DisoDeepPF and are also provided as Supplementary Data.

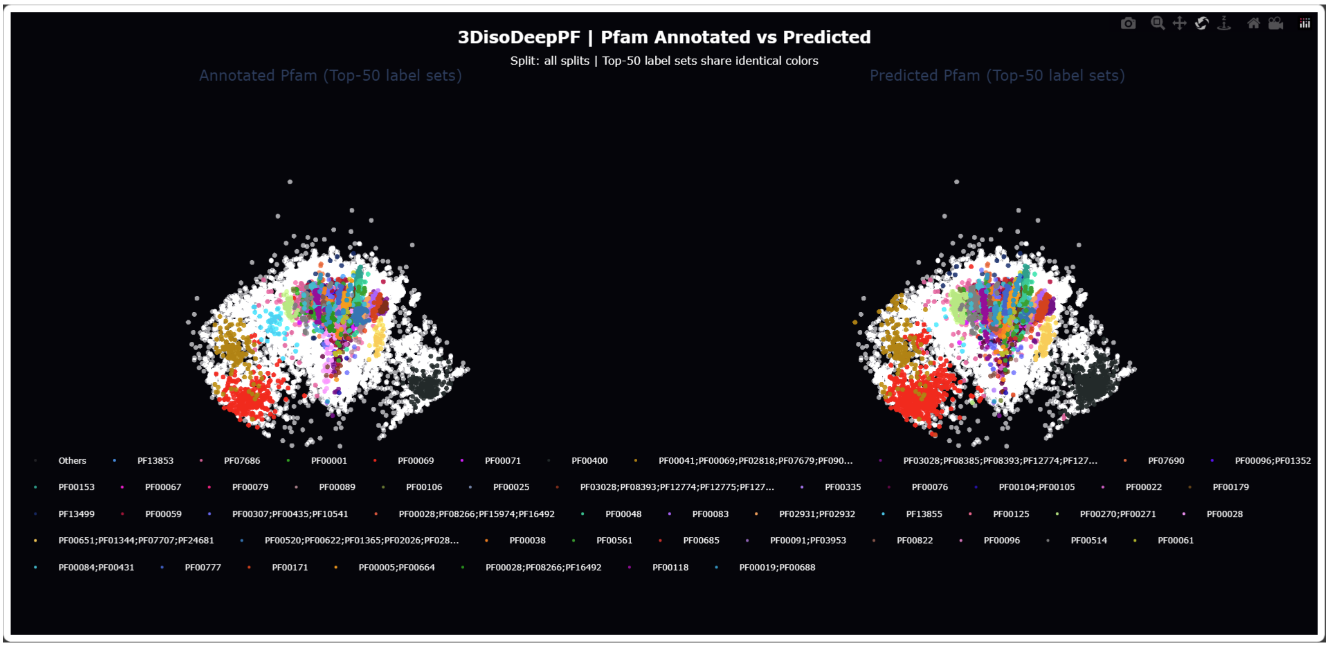

**Supplementary Fig. 7 | Interactive hyperbolic visualization of curated and predicted Pfam annotations across the integrated isoform similarity graph.** Screenshot of the interactive three-dimensional hyperbolic embedding generated from the integrated isoform similarity graph constructed from 3DisoGalaxy-derived protein nodes and annotated reference protein nodes. The embedding shows a radial, branch-like organization of the graph in hyperbolic space. The left panel displays curated Pfam annotations, whereas the right panel displays Pfam labels predicted by 3DisoDeepPF for the same protein nodes. Colors indicate the top 50 Pfam label sets, with matched colors used between the curated and predicted panels to enable direct visual comparison. White or gray nodes indicate proteins outside the displayed top Pfam label sets or without a displayed Pfam annotation. This visualization was used as an interactive qualitative companion to the quantitative Pfam prediction analyses.

### 6 Supplementary Note 6 | Implementation, baseline harmonization and evaluation metrics

#### 6.1 3DisoDeepPF implementation and training configuration

##### 6.1.1 Model Architecture and Hyperparameters

The 3DisoDeepPF framework integrates a protein language model (ESM-2) featuring a 12-layer transformer encoder to extract deep sequence representations. Graph-level context propagation is handled via a 2-layer graph neural network (GNN) module coupled with a gated multimodal fusion layer. The network enforces structural parity across embedded modalities, setting the base embedding dimensions for structural lookup nodes, linear motifs, and auxiliary task projections to 128, while the ESM sequence embeddings are projected to 512 dimensions. The intermediate hidden layer dimension dedicated to cross-modal feature fusion and sequential graph propagation is set to 256.

Model optimization is performed using the Adam optimizer with a learning rate ($\eta$) of 0.01 for a maximum of 30,000 epochs, monitored under an early stopping patience of 100 epochs to prevent over-fitting. The model operates under a transductive full-batch configuration for graph neural network training, utilizing an effective batch size of 16 exclusively during tokenized ESM feature extraction. To regularize internal representations, a dropout ratio of 0.4 is applied across the corresponding hidden layers of both the multimodal fusion and GNN modules. For multi-task scaling, the loss weight for the auxiliary task is set to 0.4, with the sequence-modality attention parameter initialized with an inductive prior weight of 2.0. Gradient descent steps are regularized by clipping the maximum gradient norm at a strict threshold of 1.0.

##### 6.1.2 Data hygiene and leakage prevention

To prevent data leakage and ensure rigorous evaluation, the framework establishes two main guardrails across the evaluation benchmarks. Under the no-GO-input rule, when predicting any specific Gene Ontology branch among GO-MF, GO-BP, or GO-CC, all GO-derived annotations are completely scrubbed from the auxiliary input tracking modules. This restriction constrains the auxiliary cross-modal tracking features exclusively to sequence models, linear motifs, and structural domains from Pfam, thereby eliminating cross-ontology shortcut leakage. For split-specific graph isolation, target labels for the validation and test partitions are strictly masked during the optimization phase, preventing information leakage.

##### 6.2 Harmonized benchmark setting for fair comparison

To minimize confounding from data partitioning, label processing and metric implementation, all methods were evaluated under a harmonized benchmark setting. Following the benchmarking practice used in recent protein function prediction studies, we standardized the evaluation environment across methods by fixing the benchmark datasets, train–validation–test partitions, label spaces, output format and metric calculation pipeline. This design was intended to ensure that performance differences primarily reflected differences in prediction strategy rather than inconsistencies in preprocessing or evaluation.

For both the CAFA-aligned canonical benchmark and the 3DisoGalaxy isoform benchmark, 3DisoDeepPF and all applicable baselines were evaluated on the same train, validation and test splits. The same protein or isoform entries were used for each benchmark, and predictions were assessed against the same held-out labels. For GO prediction, GO-MF, GO-BP and GO-CC were evaluated separately using the same ontology handling and true-path propagation procedure. For Pfam prediction, all applicable methods were evaluated against the same Pfam label universe and the same train–test label definitions.

To avoid label leakage, target labels from the evaluated ontology or Pfam space were masked from model inputs where applicable. This rule was especially important for the isoform-resolution benchmark, in which reference and non-reference isoforms from the same gene may share extensive sequence similarity and overlapping annotations. The same reference/non-reference isoform definitions and the same split assignments were used across methods.

Because different methods produce predictions in different native formats, all outputs were converted into a unified protein-by-label score matrix before evaluation. Methods were evaluated only in the label spaces for which they produced applicable predictions; for example, Pfam-HMM was used for Pfam evaluation. All reported metrics were then computed using the same evaluation scripts, score matrices, ground-truth matrices and thresholding procedure. This harmonized setting allowed direct comparison of homology transfer, profile/domain transfer, sequence-based deep learning, structure-aware deep learning, isoform-aware modeling and 3DisoDeepPF under a shared evaluation framework.

##### 6.3 Representative baseline methods

To comprehensively benchmark 3DisoDeepPF, we compared against baselines spanning four complementary families: homology transfer, profile/domain transfer, sequence-only deep learning, and isoform-aware modeling. All methods were evaluated on the same CAFA/Galaxy train-validation-test splits with the same ontology handling and metric pipeline.

*BLAST* (homology transfer) is the standard sequence-similarity baseline for protein function prediction. It quantifies how much performance can be explained by direct homology transfer alone, providing a strong and interpretable non-neural reference.

*Naive* prior baseline (label-frequency prior). The Naive baseline predicts by training-set label frequency only (protein-independent prior). This defines the minimum meaningful reference and helps verify that all stronger methods truly learn protein-specific signal rather than label imbalance.

*DeepGOPlus* and *DeepFRI* (general deep models). DeepGOPlus is a sequence-based deep model, and DeepFRI is a structure/graph-aware deep model. Including both allows us to compare 3DisoDeepPF against strong modern neural baselines from two different representation regimes (sequence-only versus structure-aware), instead of only against traditional transfer methods.

*IsoFunGO* (isoform-aware baseline). We specifically include IsoFunGO because our task is isoform-level function prediction rather than only canonical-protein prediction. IsoFunGO is an explicitly isoform-oriented method that models functional relationships at the isoform level, making it the most direct and fair baseline for testing whether 3DisoDeepPF improves isoform-specific functional discrimination.

These baselines cover the major signal categories used in protein function prediction: sequence homology, evolutionary/domain profiles, label priors, sequence deep representations, structure-aware representations and isoform-aware modelling. This baseline set therefore provides a mechanistically interpretable comparison framework for evaluating 3DisoDeepPF.

#### 6.4 Evaluation metrics

Model performance was evaluated with CAFA-aligned metrics centered on $F_{\max}$ and AUPR, with additional analyses for isoform-difference sensitivity and term informativeness.

$F_{\max}$. Let $Y_{i,j} \in \{0,1\}$ denote the ground-truth label of protein $i$ on term $j$, and $\hat{Y}_{i,j} \in [0,1]$ the predicted score. At threshold $t$, binary predictions are

$$B_{i,j}(t) = \mathbf{I}\left[\hat{Y}_{i,j} \geq t\right]. \tag{1}$$

For protein $i$,

$$\begin{aligned} TP_i(t) &= \sum_j Y_{i,j} B_{i,j}(t), \\ \text{Pred}_i(t) &= \sum_j B_{i,j}(t), \\ \text{True}_i &= \sum_j Y_{i,j}. \end{aligned} \tag{2}$$

Per-protein precision and recall are

$$\begin{aligned} \text{Prec}_i(t) &= \begin{cases} \dfrac{TP_i(t)}{\text{Pred}_i(t)}, & \text{Pred}_i(t) > 0 \\ 0, & \text{Pred}_i(t) = 0 \end{cases}, \\ \text{Rec}_i(t) &= \begin{cases} \dfrac{TP_i(t)}{\text{True}_i}, & \text{True}_i > 0 \\ 0, & \text{True}_i = 0 \end{cases}. \end{aligned} \tag{3}$$

Define $M = \{i \mid \text{True}_i > 0\}$. Macro-averaged precision and recall are

$$\begin{aligned} P_{\text{macro}}(t) &= \frac{1}{|M|} \sum_{i \in M} \text{Prec}_i(t), \\ R_{\text{macro}}(t) &= \frac{1}{|M|} \sum_{i \in M} \text{Rec}_i(t). \end{aligned} \tag{4}$$

Then

$$F(t)=\begin{cases}\dfrac{2P_{\text{macro}}(t)R_{\text{macro}}(t)}{P_{\text{macro}}(t)+R_{\text{macro}}(t)}, & P_{\text{macro}}(t)+R_{\text{macro}}(t)>0\\ 0, & \text{otherwise}\end{cases},\qquad (5)$$
$$F_{\max}=\max_{t\in\mathcal{T}}F(t),$$

where $\mathcal{T}$ is scanned with step 0.01.

AUPR (global flattened mode). We flatten $Y$ and $\hat{Y}$ over all protein-term pairs into vectors $y$ and $s$, construct the global precision-recall curve, and compute

$$\text{AUPR}=\int_0^1\text{Precision}(r)dr,\qquad (6)$$

implemented numerically as trapezoidal area on the PR curve.

micro-AUPR. For each isoform pair $p$, define a binary label vector $y^p$ and score vector $s^p$ for a task (gain or loss). Concatenate all pairs:

$$\begin{aligned}y&=\text{concat}\left(y^{(1)},\ldots,y^{(P)}\right)\\ s&=\text{concat}\left(s^{(1)},\ldots,s^{(P)}\right)\end{aligned}\qquad (7)$$

Compute one global precision-recall curve from (y,s), then

$$\text{micro-AUPR}=\int_0^1\text{Precision}(r)dr\qquad (8)$$

numerically computed by trapezoidal integration on the PR curve.

IC-binned AUPR. Let $\text{IC}(j)$ be the information content of GO term $j$, and let $B_k$ be the *k*-th IC bin:

$$B_k = \{j \mid \tau_k \le \mathrm{IC}(j) < \tau_{k+1}\} \quad (9)$$

Restrict protein-term pairs to terms in bin $B_k$:

$$\Omega_k = \{(i, j) \mid j \in B_k\} \quad (10)$$

Flatten labels/scores on $\Omega_k$ into $(y_k, s_k)$, then define

$$\mathrm{AUPR}_{B_k} = \int_0^1 \mathrm{Precision}_k(r)\,dr \quad (11)$$

This yields one AUPR per IC bin.

#### 7 Supplementary Note 7 | Evidence tracing and prediction inspection

We selected CIB1 as an illustrative test-set example for evidence tracing because it satisfied three practical criteria: it had curated Pfam and GO-MF annotations, 3DisoDeepPF recovered these annotations with high prediction scores, and its traced supporting proteins formed a biologically interpretable calcium-related neighborhood. The example was chosen for visualization after model evaluation and was not used for model training, model selection or performance estimation. We use it to demonstrate how a user can inspect the graph support behind an individual prediction, rather than to claim that all predictions have the same level of biological interpretability.

To reduce case-selection bias, we provide the full evidence-tracing outputs for all traceable test-set proteins in **Supplementary Data 2**. Each file corresponds to one query protein or isoform and reports all traceable predicted labels for that query. For each label, up to the top 20 supporting proteins are listed; labels with fewer than 20 entries indicate that fewer eligible supporting proteins were available under the tracing criteria.

**Supplementary Table 6. 3DisoDeepPF predicted Pfam and GO-MF annotation of CIB1.**

| Annotation | Ground Truth | 3DisoDeepPF |
|---|---|---|

| | | |
|---|---|---|
| Pfam | PF13499 | PF13499 (score: 0.9999999) |
| | GO:0000287 | GO:0000287 (score: 0.9930541) |
| | GO:0005509 | GO:0005509 (score: 1.0) |
| | GO:0008427 | GO:0008427 (score: 1.0) |
| GO-MF | GO:0030291 | GO:0030291 (score: 0.9999167) |
| | GO:0031267 | GO:0031267 (score: 0.99280244) |
| | GO:0043495 GO:0044325 | GO:0043495 (score: 0.999913) |
| | | GO:0044325 (score: 0.999032) |

### 8 Supplementary Note 8 | Additional results

#### 8.1 Benchmark-specific $F_{max}$ profiles across canonical and isoform-resolved settings

We compared $F_{max}$ profiles across the CAFA-aligned canonical benchmark and the 3DisoGalaxy isoform-resolved benchmark to provide a compact view of method behavior across evaluation settings (**Supplementary Fig. 8a–d**). Because the two benchmarks differ in species scope, label composition, annotation density and split design, these results should be interpreted as benchmark-specific performance profiles rather than as a direct comparison of task difficulty. Across GO-MF, GO-BP, GO-CC and Pfam, 3DisoDeepPF showed favorable performance in the isoform-resolved setting while retaining competitive performance in the canonical benchmark.

This comparison is consistent with the intended positioning of 3DisoDeepPF as a framework that connects standard protein function prediction with isoform-resolution prediction. The canonical benchmark provides a conventional reference for protein-level function prediction, whereas the 3DisoGalaxy benchmark evaluates performance in an isoform-rich setting where closely related proteins may differ in domains, motifs or local structural regions. The paired profiles therefore show that 3DisoDeepPF remains informative across both conventional and isoform-resolved evaluation contexts, without implying that the two benchmark settings have equivalent difficulty.

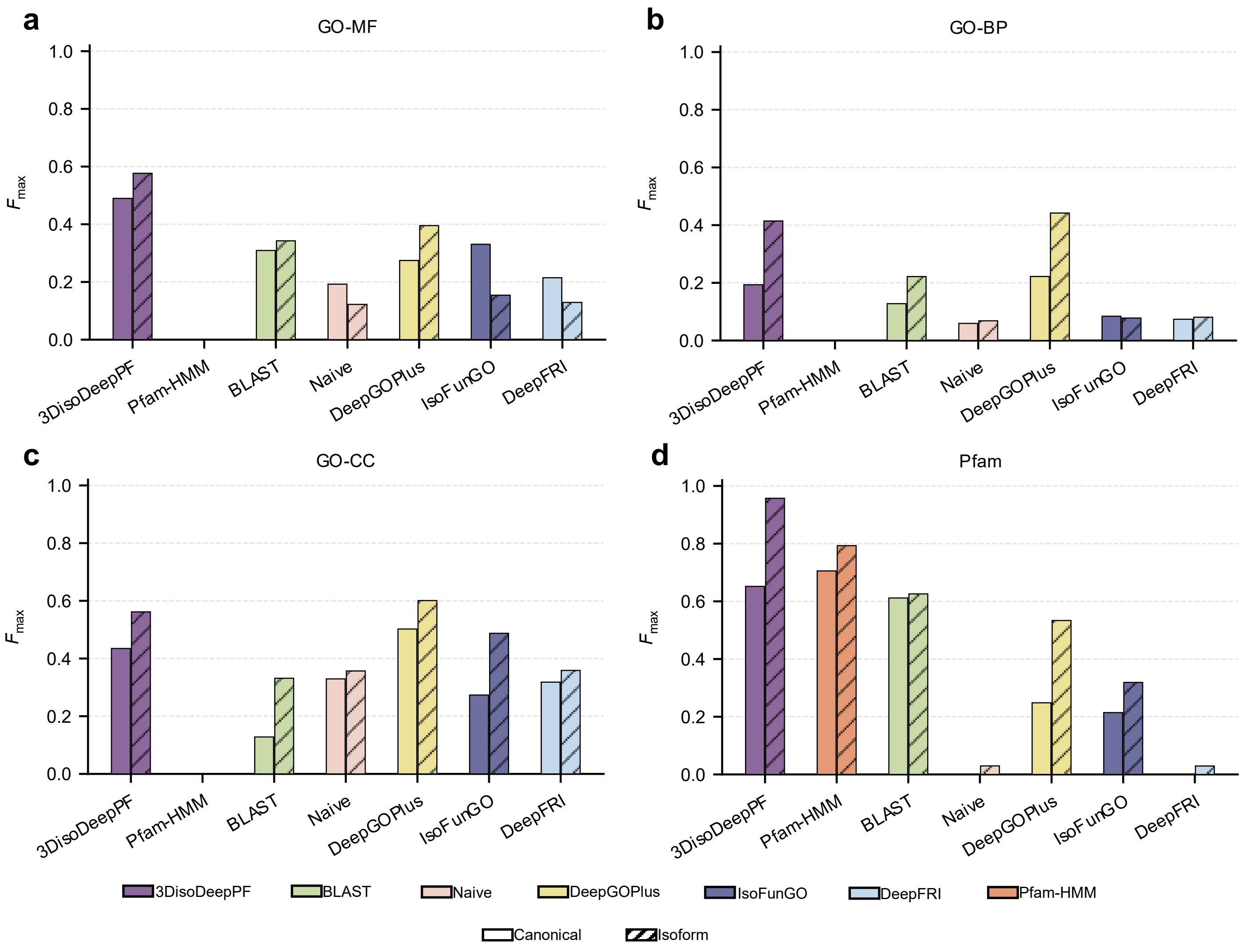

**Supplementary Fig. 8 | Fmax profiles across canonical and isoform-resolved benchmarks. a–d,** Comparison of Fmax scores across the CAFA-aligned canonical benchmark and the 3DisoGalaxy isoform-resolved benchmark for GO-MF (**a**), GO-BP (**b**), GO-CC (**c**) and Pfam (**d**) prediction tasks. Solid bars indicate performance on the canonical benchmark, and hatched bars indicate performance on the isoform-resolved benchmark. Methods are shown using consistent colors across panels. The comparison summarizes benchmark-specific performance profiles and is not intended as a direct comparison of intrinsic task difficulty between the canonical and isoform-resolved settings.

#### 8.2 Distribution and recurrence of predicted labels across Pfam and Gene Ontology label spaces

We next summarized the predicted label landscape generated by 3DisoDeepPF across Pfam and the three Gene Ontology branches: biological process (GO-BP), cellular component (GO-CC) and molecular function (GO-MF). This analysis was intended to describe prediction coverage, missingness, no-prediction-associated input features and label-frequency structure across the breast cancer isoform atlas, rather than to assess prediction accuracy.

Most protein isoforms received at least one predicted label in each annotation space, with predicted-label coverage ranging from 93.3% for Pfam to 94.3% for GO-BP **(Supplementary Fig. 9a)**. Isoforms without predicted labels were unevenly distributed across label spaces. The largest missing-label group consisted of isoforms without predicted labels in all four annotation spaces, whereas smaller groups lacked predictions in selected combinations of Pfam and GO labels **(Supplementary Fig. 9b)**. This overlap suggests that missing predictions were concentrated in a subset of isoforms rather than being restricted to one annotation space.

To further characterize this subset, we examined input-level features associated with the absence of predicted labels across all four label spaces. In an adjusted logistic-regression model, lower pLDDT, shorter ORF length, higher predicted disordered ratio and singleton isoform-family status were each associated with higher odds of lacking predicted labels **(Supplementary Fig. 9c)**. These associations are consistent with the interpretation that no-prediction isoforms may have weaker structural evidence, shorter sequence context, higher disorder or limited isoform-family context. Because this analysis uses computational features and prediction outputs, it should be interpreted as a diagnostic association rather than evidence for a causal mechanism.

The predicted label load also differed across annotation spaces. GO-BP showed a broader per-isoform prediction load than Pfam, GO-CC and GO-MF **(Supplementary Fig. 9d)**. At the vocabulary level, GO-BP also contained the largest number of distinct predicted labels, followed by Pfam, GO-MF and GO-CC **(Supplementary Fig. 9e)**. These differences are expected in part because Pfam and GO differ in vocabulary size, ontology structure and biological granularity. Therefore, label counts should not be interpreted as directly comparable measures of biological complexity across annotation spaces.

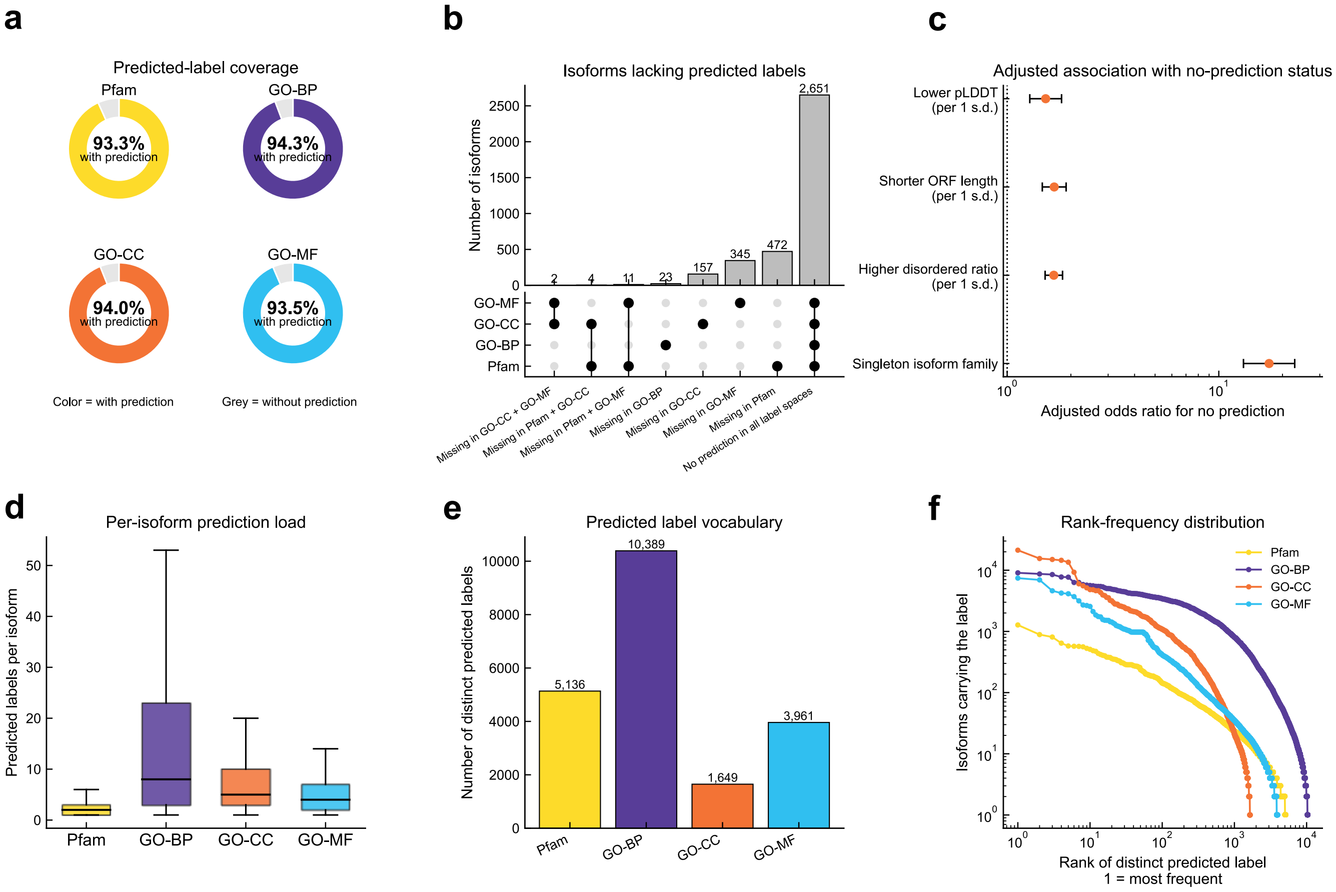

**Supplementary Fig. 9 | Predicted label coverage and no-prediction-associated features across Pfam and Gene Ontology label spaces. a**, Fraction of protein isoforms with at least one predicted label in Pfam, Gene Ontology biological process (GO-BP), cellular component (GO-CC) or molecular function (GO-MF). Colored sectors denote isoforms with predicted labels; grey sectors denote isoforms without predicted labels. **b**, Overlap of isoforms without predicted labels across the four label spaces. Bars show the number of isoforms in each intersection, and connected dots indicate the label spaces involved. **c**, Adjusted logistic-regression analysis of features associated with no-prediction status. No-prediction isoforms were defined as protein isoforms lacking predicted labels in Pfam, GO-BP, GO-CC and GO-MF. Continuous variables were scaled per one standard deviation in the indicated direction. Odds ratios above 1 indicate higher odds of lacking predicted labels. **d**, Number of predicted labels per isoform in each label space. **e**, Number of distinct predicted labels used in each label space. **f**, Rank-frequency distribution of predicted labels, showing the long-tailed frequency structure of each predicted label space.

Finally, rank-frequency analysis showed a long-tailed distribution across all four label spaces, with a limited number of broadly assigned labels and many labels assigned to relatively few isoforms **(Supplementary Fig. 9f)**. Together, these analyses provide a descriptive overview of predicted-label coverage, no-prediction-associated features and label-frequency structure used in downstream isoform-family and functional-remodeling analyses.

##### 8.3 Isoform variation within isoform family predicted by 3DisoDeepPF

Predicted isoform-level annotation remodeling was assessed by comparing each non-reference isoform with the reference isoform from the same gene across Pfam and the three Gene Ontology branches. Each isoform-reference pair was classified as unchanged, gain, loss or reconfigured according to whether predicted labels were retained, newly gained, lost or simultaneously gained and lost (**Supplementary Fig. 10a**). Pfam labels were largely conserved, whereas GO-BP and GO-MF showed broader predicted remodeling, indicating that isoform-level differences were more frequently reflected in process- and function-level annotations than in domain composition.

We next quantified the magnitude of these changes. GO-BP showed the largest number of changed labels per changed isoform-reference pair, followed by GO-CC and GO-MF, whereas Pfam changes were typically smaller in scale (**Supplementary Fig. 10b**). Consistently, large-shift analysis showed that GO-BP retained the highest fraction of changed pairs even under increasingly stringent thresholds for the minimum number of changed labels (**Supplementary Fig. 10c**). These results suggest that predicted isoform-level remodeling is label-space dependent, with Pfam capturing more discrete domain changes and GO-BP capturing broader functional shifts.

To examine cancer relevance, we compared changed isoforms from cancer-driver genes with those from other genes. Cancer-driver genes showed a higher fraction of isoforms with predicted label changes across all four annotation spaces (**Supplementary Fig. 10d**). Enrichment analysis further confirmed that changed isoforms were enriched in cancer-driver genes at both isoform and gene levels (**Supplementary Fig. 10e**). This indicates that predicted isoform-level functional remodeling is not randomly distributed, but is preferentially associated with genes of known cancer relevance.

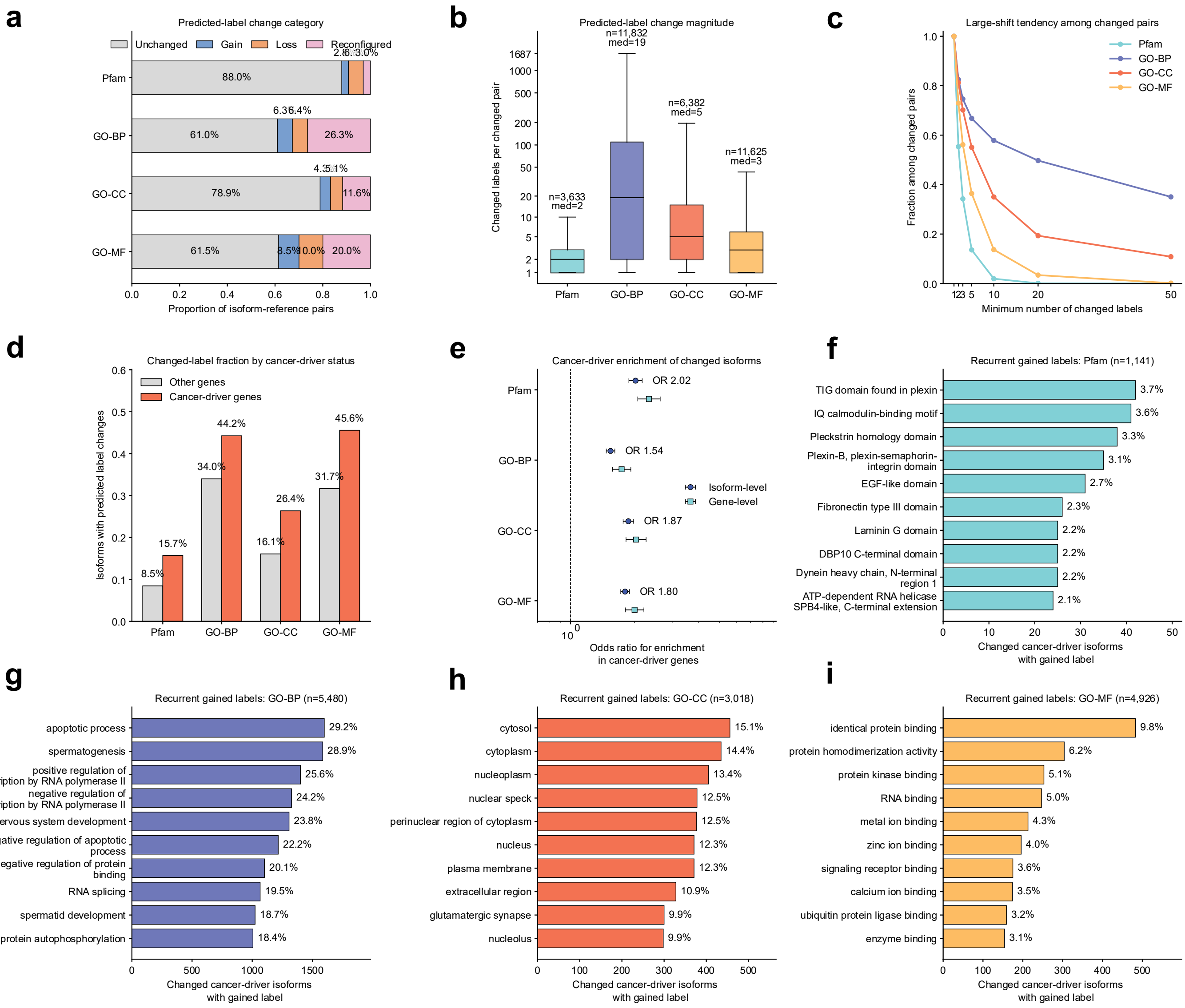

**Supplementary Fig. 10 | Predicted isoform-level annotation remodeling and cancer-driver association. a,** Proportion of isoform-reference pairs classified as unchanged, gain, loss or reconfigured across Pfam, GO-BP, GO-CC and GO-MF predicted labels. Gain indicates labels present in the non-reference isoform but absent from the reference isoform; loss indicates labels absent from the non-reference isoform but present in the reference; reconfigured indicates simultaneous gain and loss. **b,** Distribution of change magnitude among changed isoform-reference pairs, measured as the number of changed labels per pair. Box plots show the median and interquartile range; numbers indicate the number of changed pairs and median changed-label count. **c,** Large-shift tendency, showing the fraction of changed pairs retained after requiring at least the indicated number of changed labels. **d,** Fraction of isoforms with predicted label changes in cancer-driver genes and other genes. Cancer-driver genes showed a higher fraction of changed isoforms across all four label spaces. **e,** Odds-ratio analysis for enrichment of changed isoforms in cancer-driver genes at isoform and gene levels. The dashed line indicates no enrichment. **f–i,** Most frequent gained labels among changed cancer-driver isoforms for Pfam (**f**), GO-BP (**g**), GO-CC (**h**) and GO-MF (**i**). These labels provide functional context for the cancer-driver-associated remodeling observed in **d** and **e**, highlighting predicted changes related to interaction and signaling domains, apoptosis and transcriptional regulation, nuclear/cytoplasmic and membrane-associated localization, and binding-related molecular functions.

Finally, we summarized the most frequent gained labels among changed cancer-driver isoforms to interpret the functional content of this remodeling. Gained Pfam labels included interaction- and signaling-related domains, such as plexin-associated, calmodulin-binding, pleckstrin homology, EGF-like and extracellular matrix-associated domains (**Supplementary Fig. 10f**). Gained GO-BP labels highlighted apoptosis, transcriptional regulation, RNA splicing and phosphorylation-related processes (**Supplementary Fig. 10g**). Gained GO-CC labels pointed to nuclear, cytoplasmic, membrane-associated and extracellular compartments (**Supplementary Fig. 10h**), whereas gained GO-MF labels emphasized protein binding, dimerization, kinase binding, RNA binding and ion-binding functions (**Supplementary Fig. 10i**). Together, these analyses provide functional context for the cancer-driver-associated remodeling observed in **Supplementary Fig. 10d,e**, supporting the use of isoform-resolved prediction to prioritize cancer-relevant functional shifts.

##### 8.4 Prioritization of tumor-associated isoform functional-shift candidates

After showing that predicted isoform-level GO remodeling was associated with cancer-driver genes, we next asked whether these predictions could be used to define a focused catalogue of tumor-associated isoform functional-shift candidates. The aim of this analysis was to support candidate inspection and downstream hypothesis generation, rather than to establish validated therapeutic targets.

We first selected isoforms whose 3DisoDeepPF-predicted GO annotations overlapped cancer hallmark-associated GO terms. The hallmark GO set was derived from the consensus mapping by Chen et al. (2021), who compared Gene Ontology- and pathway-based mappings of cancer hallmarks and defined consensus GO terms supported across mapping schemes. This step connected the predicted GO changes described above with cancer-relevant biological processes.

We then retained candidates with tumor-associated evidence and isoform-level support. The final catalogue records each candidate's identifier, canonical or non-canonical class, gene symbol, associated target, reference isoform, 3DisoDeepPF-predicted GO/Pfam annotations, available UniProtKB annotations, ORF support, protein length, structural confidence, transcript structural category, detection rate, localization, isoform-specificity tier and final integrated prioritization score. Incomplete-splice-match candidates were removed during filtering, while canonical and non-canonical non-ISM candidates were retained.

The resulting candidate catalogue is provided as **Supplementary Data 3**, and the corresponding column definitions are provided in **Supplementary Table 7**. These candidates should be interpreted as prioritized tumor-associated isoform functional-shift candidates for further inspection and experimental follow-up, not as validated functional mechanisms or therapeutic targets.

**Supplementary Table 7. Column definitions for the prioritized catalogue of tumor-associated isoform functional-shift candidates.**

| Column | Description |
|---|---|
| id | Candidate isoform or protein identifier used as the primary entry identifier. |
| id_class | Candidate class based on annotation status. canonical denotes candidates matched to annotated transcript isoforms in public reference transcript catalogues, such as Ensembl/GENCODE. non-canonical denotes isoform-derived candidates not represented as annotated reference isoforms in those catalogues. |
| gene_symbol | Gene symbol associated with the candidate. |
| target | Whether a target and target gene symbol curated by Therapeutic target database 2026 |
| reference_isoform_id | Annotated reference isoform used as the within-gene comparator for isoform-level interpretation. |
| 3DisoDeepPF_pred_go_bp_label | GO biological process labels predicted by 3DisoDeepPF. These labels describe predicted biological processes and were used in the cancer hallmark-associated functional filtering step. |
| 3DisoDeepPF_pred_go_mf_label | GO molecular function labels predicted by 3DisoDeepPF. These labels describe predicted molecular activities, such as binding or catalytic functions, where available. |
| 3DisoDeepPF_pred_go_cc_label | GO cellular component labels predicted by 3DisoDeepPF. These labels describe predicted cellular-component or localization-associated annotations. |

| Column | Description |
| --- | --- |
| 3DisoDeepPF_pred_pfam | Pfam labels predicted by 3DisoDeepPF, representing predicted protein domain or family-level annotations. |
| uniprotKB_anno_go_bp_label | Existing UniProtKB GO biological process annotation where available. Empty values indicate no corresponding annotation in the integrated table. |
| uniprotKB_anno_go_mf_label | Existing UniProtKB GO molecular function annotation where available. |
| uniprotKB_anno_go_cc_label | Existing UniProtKB GO cellular component annotation where available. |
| uniprotKB_anno_pfam | Existing UniProtKB/Pfam annotation where available. |
| ORF_type | ORF classification reported in the 3DisoGalaxy isoform translatome analysis datasets predicted by RiboCode. This field is retained as input evidence and was not redefined during candidate-list construction. |
| mean_ORF_count | Mean ORF-support count across supporting evidence. Positive values were used as translation-support evidence in the isoform-specificity assessment. |
| protein_length | Length of the predicted protein product in amino acids. |
| pLDDT | Mean predicted local distance difference test score of the structural model. pLDDT is reported on a 0–100 scale. Scores >90 are generally interpreted as very high local confidence, 70–90 as confident, 50–70 as low confidence and <50 as very low confidence. This metric reflects local structural confidence and does not assess relative domain orientation. |
| structural_category | Transcript structural category reported by the transcript reconstruction/annotation workflow by SQANTI3. Incomplete-splice-match entries were removed during filtering; full-splice-match and other non-incomplete categories were retained. |

| Column | Description |
| --- | --- |
| DetectionRate | Fraction or frequency of samples in which the candidate was detected. This field was used as a detectability or prevalence proxy, not as evidence of functional causality. |
| Localizations | Predicted or annotated subcellular localization information used for localization-based feasibility assessment. |
| isoform_specificity_tier | Tier summarizing isoform-level support. The tier was derived from a composite score based on translation support, junction/structural-category evidence and sequence-divergence proxies. Higher tiers indicate stronger isoform-level support in the prioritization workflow. |
| final_integrated_score | Heuristic ranking score used to order candidates after integrated filtering. The score was used only for prioritization and hypothesis generation, not as a calibrated probability, functional validation score or therapeutic target score. |

To move beyond catalogue-level prioritization, we selected twelve non-canonical protein isoform candidates from **Supplementary Data 3** and inspected how their predicted functional profiles differed from their matched canonical reference proteins. The examples were arranged according to the dominant type of difference. **Supplementary Fig. 11** focuses on candidates in which the main signal is GO-driven functional remodeling, whereas **Supplementary Fig. 12** focuses on domain-level changes or mixed structural–functional divergence. To make representative isoform-level changes interpretable, we separated the candidate examples by the annotation layer in which the query and reference proteins differed most clearly. Query protein isoforms were labelled by the Ensembl transcript ID of the transcript model used to derive the coding sequence, whereas canonical reference proteins were labelled by UniProtKB accessions, allowing each non-canonical isoform model to remain traceable to its source transcript and each reference protein to remain linked to a curated protein entry (Dyer et al., 2025; The UniProt Consortium, 2025). Ensembl provides genome and transcript annotations, while UniProtKB provides protein sequence and functional annotation resources. The comparison summarizes query-specific predicted labels, reference-specific annotated labels and shared labels across GO and Pfam, which describe biological processes, molecular functions, cellular components and protein-

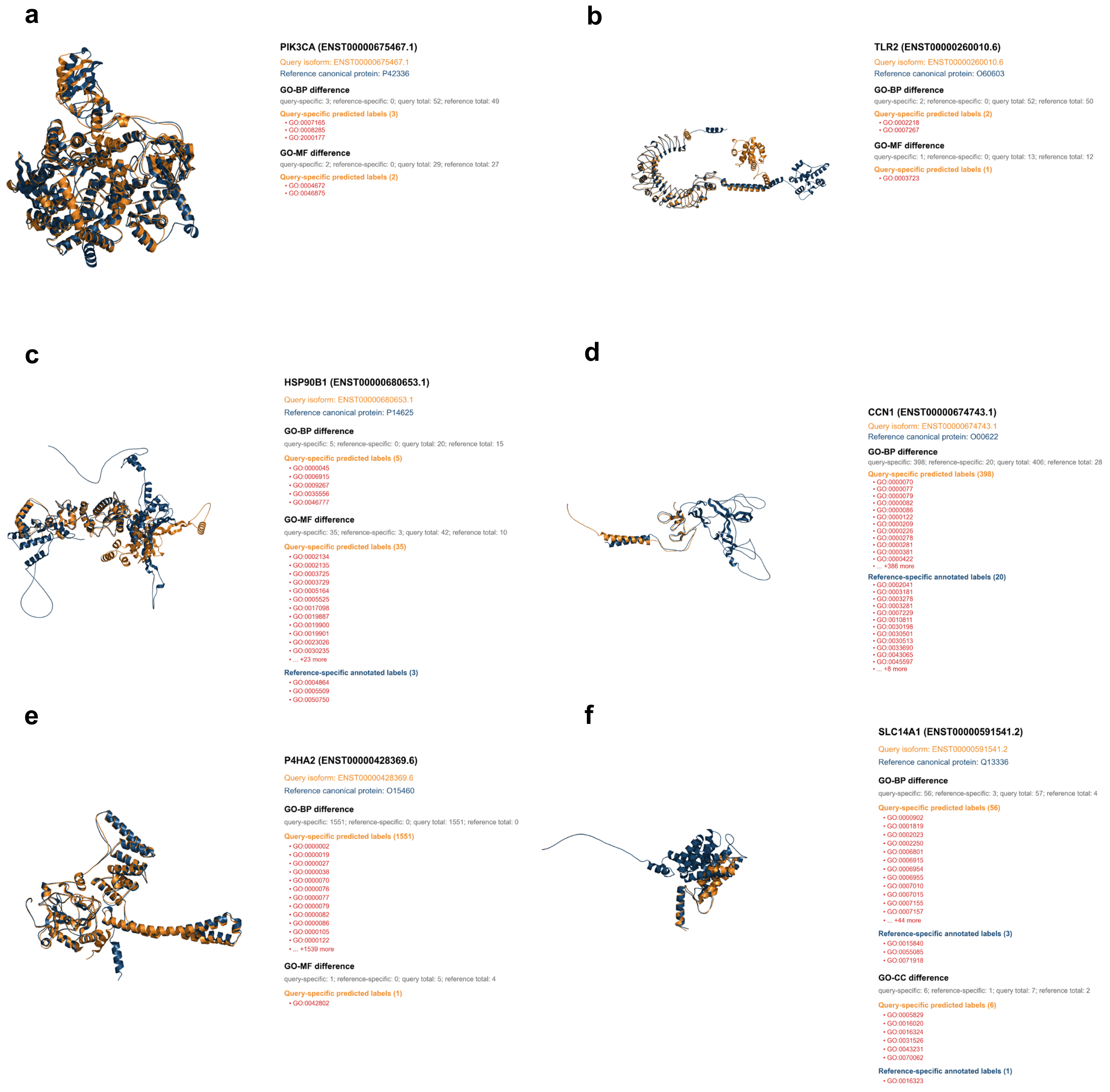

**Supplementary Fig. 11 | GO-label divergence in tumor-associated non-canonical protein isoform candidates. a–f**, Structural alignment and GO-label comparison between selected query protein isoforms and their matched canonical reference proteins: **a**, PIK3CA ENST00000675467.1 versus P42336; **b**, TLR2 ENST00000260010.6 versus O60603; **c**, HSP90B1 ENST00000608653.1 versus P14625; **d**, CCN1 ENST00000674743.1 versus O00622; **e**, P4HA2 ENST00000428369.6 versus O15460; and **f**, SLC14A1 ENST00000591541.2 versus Q13336. Query isoforms are shown in yellow and canonical references in blue. For each query isoform, the label indicates the Ensembl transcript ID of the transcript model used to derive its coding sequence; the matched canonical protein is labelled by its UniProtKB accession. Label summaries show query-specific predicted labels, reference-specific annotated labels and shared labels across GO biological process, molecular function and cellular component terms.

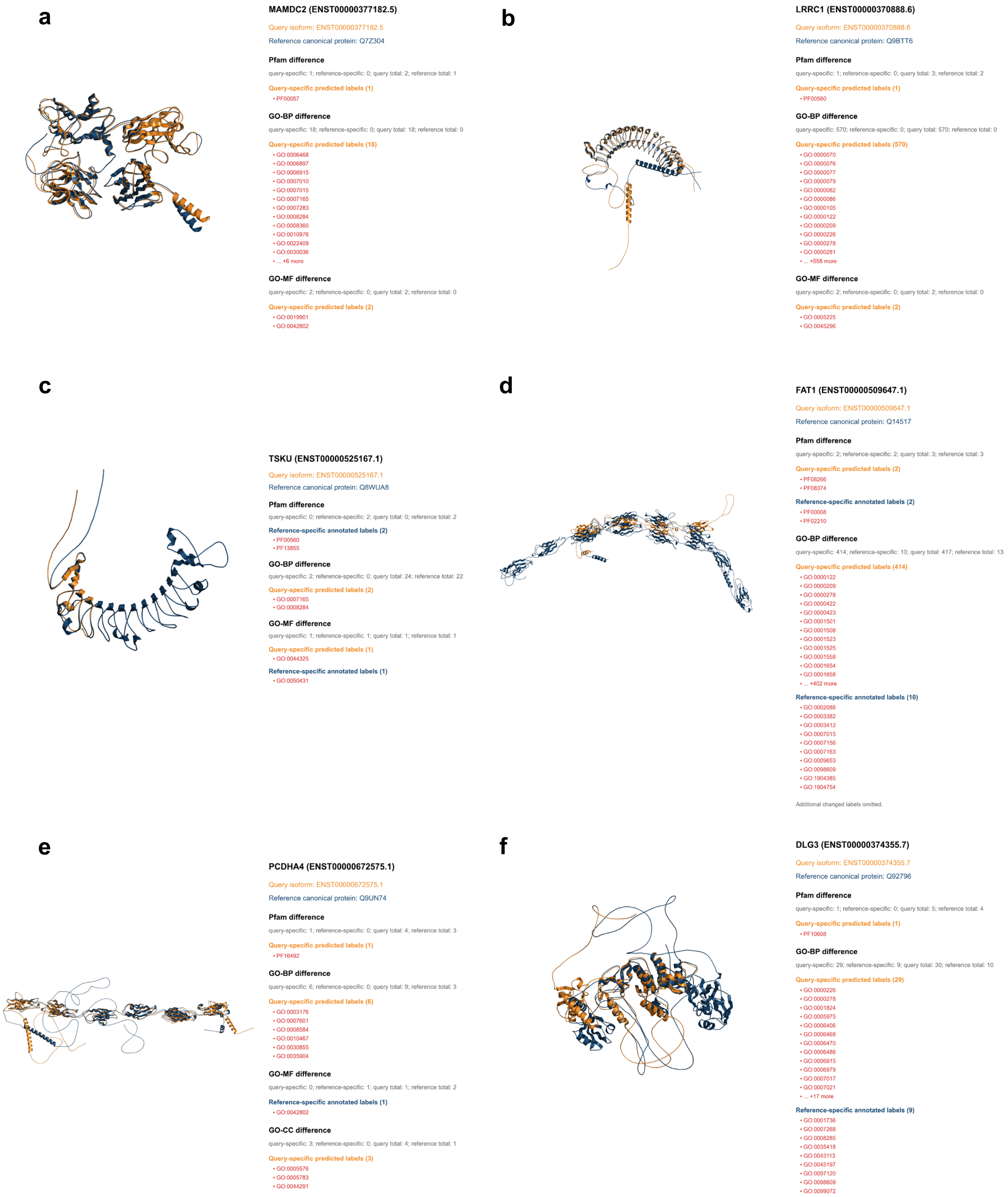

**Supplementary Fig. 12 | Pfam-domain divergence in tumor-associated non-canonical protein isoform candidates. a–f**, Structural alignment and Pfam/GO-label comparison between selected query protein isoforms and their matched canonical reference proteins: **a**, MAMDC2 ENST00000377182.5 versus Q7Z304; **b**, LRRC1 ENST00000370888.6 versus Q9BTT6; **c**, TSKU ENST00000525167.1 versus Q8WUA8; **d**, FAT1 ENST00000509647.1 versus Q14517; **e**, PCDHA4 ENST00000672575.1 versus Q9UN74; and **f**, DLG3 ENST00000374355.7 versus Q92796. Query isoforms are shown in yellow and canonical references in blue. For each query isoform, the label indicates the Ensembl transcript ID of the transcript model used to derive its coding sequence; the matched canonical protein is labelled by its UniProtKB accession. Label summaries show query-specific predicted labels, reference-specific annotated labels and shared labels across Pfam domains and GO biological process, molecular function and cellular component terms.

family/domain features (The Gene Ontology Consortium, 2026; Paysan-Lafosse et al., 2025; The UniProt Consortium, 2025).

The GO-driven examples first asked whether the predicted differences pointed back to recognizable cancer-relevant biology. PIK3CA provided the clearest pathway-level case in **Supplementary Fig. 11a**. The query isoform gained labels for signal transduction (GO:0007165), phosphatidylinositol-mediated signaling (GO:0048015) and transferase activity transferring phosphorus-containing groups (GO:0016772). These terms match the expected PI3K signaling axis and are consistent with the role of PIK3CA as a clinically relevant PI3K catalytic-subunit gene in breast cancer (André et al., 2019; The UniProt Consortium, 2025). This example should not be read as proof of an isoform-specific PIK3CA mechanism, but it shows that the predicted query-specific labels recover the correct pathway context for a high-value cancer gene.

A more conservative example was TLR2 in **Supplementary Fig. 11b**. Here, the query-specific gains were limited to activation of innate immune response (GO:0002218) and cell–cell signaling (GO:0007267). These labels are closely aligned with TLR2 receptor biology, which centers on innate immune recognition and inflammatory signaling (The UniProt Consortium, 2025). The small number of gained terms makes this panel useful as a sanity-check example: the model does not need a large label expansion to recover a coherent immune and tumour-microenvironment-related axis.

HSP90B1 in **Supplementary Fig. 11c** extended the same logic to protein-homeostasis biology. HSP90B1 encodes GRP94/endoplasmin, an endoplasmic-reticulum HSP90-family chaperone involved in protein folding and quality control (Marzec et al., 2012; The UniProt Consortium, 2025). The figure shows predicted BP and MF differences relative to the canonical reference protein. Because the exact term list should be finalized from the source table rather than read manually from the image, this panel is best framed at the pathway level: the candidate falls within a proteostasis and stress-response axis that is biologically consistent with the gene.

The remaining panels in **Supplementary Fig. 11** illustrate broader GO-driven remodeling. CCN1 in **Supplementary Fig. 11d** gained a large set of predicted BP labels. Rather than interpreting each term separately, the most relevant visible terms are angiogenesis (GO:0001525) and response to hypoxia (GO:0001666), which are consistent with the matricellular role of CCN1/CYR61 in adhesion, migration, angiogenesis, inflammation and tissue remodeling (Lau, 2011; The UniProt Consortium, 2025). P4HA2 in **Supplementary Fig. 11e** showed an even broader BP expansion. Its gene-level context is extracellular matrix and collagen modification, a process connected to breast cancer invasion and metastasis (Cox, 2021). In both cases, the biological message is not a single precise mechanism, but a broad candidate-level signal pointing to tumour-associated extracellular-matrix and stress-response programs.

SLC14A1 in **Supplementary Fig. 11f** showed a more contrasted pattern. The canonical reference retained transporter-associated labels, including urea transport (GO:0015840), transmembrane transport (GO:0055085) and urea transmembrane transport (GO:0071918). By contrast, the query isoform gained labels related to inflammatory response (GO:0006954), immune response (GO:0006955), apoptotic process (GO:0006915), cell adhesion (GO:0007155), signal transduction (GO:0007165) and positive regulation of cell population proliferation (GO:0008284), together with cellular-component labels such as membrane (GO:0016020), extracellular exosome (GO:0070062) and cytosol (GO:0005829). This contrast is biologically plausible as a predicted shift from canonical transporter annotation toward broader tumour-associated processes, although SLC14A1 should still be interpreted cautiously because its best-established cancer association is in bladder cancer susceptibility rather than breast cancer (Garcia-Closas et al., 2011; The UniProt Consortium, 2025).

The second group focused on cases where domain-level or bidirectional remodeling was more informative than GO gain alone. MAMDC2 was the cleanest example in **Supplementary Fig. 12a**. The query isoform gained the LDL receptor class A domain PF00057, together with predicted labels related to apoptotic process (GO:0006915), cytoskeleton organization (GO:0007010), actin filament organization (GO:0007015), signal transduction (GO:0007165), positive regulation of cell population proliferation (GO:0008284), regulation of cell shape (GO:0008360), protein kinase binding (GO:0019901) and identical protein binding (GO:0042802). This pattern is coherent for an extracellular domain-containing protein and fits reports that MAMDC2 may be a breast cancer biomarker with tumour-suppressive activity (Lee et al., 2020). It is therefore a useful domain-level candidate, while still remaining computational rather than experimentally validated.

LRRC1 in **Supplementary Fig. 12b** showed a related but more cautious pattern. The query gained PF00560, a leucine-rich-repeat-related Pfam label, which is consistent with the architecture of an LRR-containing protein. The same panel also contains broad BP gains, including cell-cycle and checkpoint-associated terms such as DNA-damage checkpoint signaling (GO:0000077), G1/S transition of mitotic cell cycle

(GO:0000082), G2/M transition of mitotic cell cycle (GO:0000086) and mitotic cell cycle (GO:0000278). These labels are cancer-relevant, but their breadth makes them less specific. The domain gain, rather than the size of the GO-BP set, is therefore the most defensible interpretation.

TSKU in **Supplementary Fig. 12c** linked the two evidence types. The query gained only two BP labels, signal transduction (GO:0007165) and positive regulation of cell population proliferation (GO:0008284), whereas the canonical reference retained LRR-related Pfam labels PF00560 and PF13555. This creates a compact example in which limited functional gain is accompanied by possible domain-level divergence. Tsukushi is an extracellular small leucine-rich proteoglycan involved in signaling and developmental processes, and cancer studies have linked TSKU to proliferation, epithelial–mesenchymal transition or immune-infiltration-associated prognosis in lung cancer (Yamada et al., 2019; Huang et al., 2021). The evidence therefore supports gene-program compatibility rather than a breast-cancer-specific mechanism.

FAT1, PCDHA4 and DLG3 in **Supplementary Fig. 12d–f** illustrate mixed remodeling, where both query-specific and reference-specific differences are informative. FAT1 showed bidirectional Pfam and GO-BP differences, with readable query-specific labels including negative regulation of transcription by RNA polymerase II (GO:0000122), protein polyubiquitination (GO:0000209), mitotic cell cycle (GO:0000278) and mitophagy (GO:0000422). These terms point to regulatory and cell-state processes that are plausible for a large atypical cadherin involved in cancer-associated adhesion and signaling biology (Wang et al., 2025). PCDHA4 showed Pfam, BP and CC differences consistent with protocadherin membrane and adhesion biology, although exact term-level claims should rely on the source table. DLG3 showed one query-specific Pfam label together with mixed BP remodeling; as a MAGUK scaffold protein, it is better interpreted through altered scaffold, junction or interaction-related functions than through a single gained process (The UniProt Consortium, 2025).

Taken together, these examples support a restrained conclusion. The strongest cases are not those with the largest number of predicted labels, but those in which the direction of change is specific and compatible with known gene biology. PIK3CA and TLR2 recover pathway and immune-receptor biology through compact GO gains; MAMDC2 and LRRC1 show domain-level changes that match protein architecture; TSKU combines limited BP gain with domain divergence; and FAT1, PCDHA4 and DLG3 illustrate broader structural–functional divergence in adhesion or scaffold-related genes.

These observations support the use of 3DisoDeepPF as a candidate-ranking and inspection framework for disease-associated protein isoforms, while remaining computational predictions that require transcript-, protein- and function-level validation.

#### References

- Aebersold, R. *et al.* How many human proteoforms are there? *Nat. Chem. Biol.* **14**, 206–214 (2018).
- André, F. *et al.* Alpelisib for PIK3CA-mutated, hormone receptor-positive advanced breast cancer. *N. Engl. J. Med.* **380**, 1929–1940 (2019).
- Barrio-Hernandez, I. *et al.* Clustering predicted structures at the scale of the known protein universe. *Nature* **622**, 637–645 (2023).
- Berman, H.M. *et al.* The Protein Data Bank. *Nucleic Acids Res.* **28**, 235–242 (2000).
- Berman, H.M., Henrick, K., Nakamura, H. & Markley, J.L. The worldwide Protein Data Bank (wwPDB): ensuring a single, uniform archive of PDB data. *Nucleic Acids Res.* **35**, D301–D303 (2007).
- Boadu, F., Cao, H. & Cheng, J. Combining protein sequences and structures with transformers and equivariant graph neural networks to predict protein function. *Bioinformatics* **39**, i318–i325 (2023).
- Cao, Y. & Shen, Y. TALE: Transformer-based protein function annotation with joint sequence–label embedding. *Bioinformatics* **37**, 2825–2833 (2021).
- Chen, Y., Verbeek, F.J. & Wolstencroft, K. Establishing a consensus for the hallmarks of cancer based on gene ontology and pathway annotations. *BMC Bioinformatics* **22**, 178 (2021).
- Durairaj, J. *et al.* Uncovering new families and folds in the natural protein universe. *Nature* **622**, 646–653 (2023).
- Dyer, S.C. *et al.* Ensembl 2025. *Nucleic Acids Res.* **53**, D948–D957 (2025).
- Ellis, J.D. *et al.* Tissue-specific alternative splicing remodels protein–protein interaction networks. *Mol. Cell* **46**, 884–892 (2012).
- Garcia-Closas, M. *et al.* A genome-wide association study of bladder cancer identifies a new susceptibility locus within *SLC14A1*, a urea transporter gene on chromosome 18q12.3. *Hum. Mol. Genet.* **20**, 4282–4289 (2011).
- Gligorijević, V. *et al.* Structure-based protein function prediction using graph convolutional networks. *Nat. Commun.* **12**, 3168 (2021).
- Huang, H. *et al.* Tsukushi is a novel prognostic biomarker and correlates with tumor-infiltrating B cells in non-small cell lung cancer. *Aging* **13**, 4428–4451 (2021).

- Huang, J. *et al.* Discovery of deaminase functions by structure-based protein clustering. *Cell* **186**, 3182–3195.e14 (2023).
- Jiang, F. T. et al. The Structural Code of Breast Cancer Proteoform: Alternative Splicing-driven Protein Isoform Variation and Functional Diversification. Preprint at bioRxiv https://doi.org/10.64898/2026.04.30.722115 (2026).
- Jiang, Y. *et al.* An expanded evaluation of protein function prediction methods shows an improvement in accuracy. *Genome Biol.* **17**, 184 (2016).
- Jiao, P. *et al.* Struct2GO: protein function prediction based on graph pooling algorithm and AlphaFold2 structure information. *Bioinformatics* **39**, btad637 (2023).
- Kulmanov, M., Khan, M.A. & Hoehndorf, R. DeepGO: predicting protein functions from sequence and interactions using a deep ontology-aware classifier. *Bioinformatics* **34**, 660–668 (2018).
- Kulmanov, M. & Hoehndorf, R. DeepGOPlus: improved protein function prediction from sequence. *Bioinformatics* **36**, 422–429 (2020).
- Lau, A.M. *et al.* Exploring structural diversity across the protein universe with The Encyclopedia of Domains. *Science* **386**, eadq4946 (2024).
- Lau, L.F. CCN1/CYR61: the very model of a modern matricellular protein. *Cell. Mol. Life Sci.* **68**, 3149–3163 (2011).
- Lee, H. *et al.* MAM domain containing 2 is a potential breast cancer biomarker that exhibits tumour-suppressive activity. *Cell Prolif.* **53**, e12883 (2020).
- Liu, Y. *et al.* Impact of alternative splicing on the human proteome. *Cell Rep.* **20**, 1229–1241 (2017).
- Marzec, M., Eletto, D. & Argon, Y. GRP94: an HSP90-like protein specialized for protein folding and quality control in the endoplasmic reticulum. *Biochim. Biophys. Acta* **1823**, 774–787 (2012).
- Nilsen, T.W. & Graveley, B.R. Expansion of the eukaryotic proteome by alternative splicing. *Nature* **463**, 457–463 (2010).
- Nomburg, J. *et al.* Birth of protein folds and functions in the virome. *Nature* **633**, 710–717 (2024).
- Paysan-Lafosse, T. *et al.* The Pfam protein families database: embracing AI/ML. *Nucleic Acids Res.* **53**, D523–D534 (2025).
- Qiu, S., Yu, G., Lu, X., Domeniconi, C. & Guo, M. Isoform function prediction by Gene Ontology embedding. *Bioinformatics* **38**, 4581–4588 (2022).
- Radivojac, P. *et al.* A large-scale evaluation of computational protein function prediction. *Nat. Methods* **10**, 221–227 (2013).
- Scotti, M.M. & Swanson, M.S. RNA mis-splicing in disease. *Nat. Rev. Genet.* **17**, 19–32 (2016).

- Smith, L.M. & Kelleher, N.L. Proteoform: a single term describing protein complexity. *Nat. Methods* **10**, 186–187 (2013).
- Smith, L.M. *et al.* Proteoforms as the next proteomics currency. *Science* **359**, 1106–1107 (2018).
- The UniProt Consortium. UniProt: the Universal Protein Knowledgebase in 2025. *Nucleic Acids Res.* **53**, D609–D617 (2025).
- Urbanski, L.M., Leclair, N. & Anczuków, O. Alternative-splicing defects in cancer: splicing regulators and their downstream targets, guiding the way to novel cancer therapeutics. *WIREs RNA* **9**, e1476 (2018).
- Varadi, M. *et al.* AlphaFold Protein Structure Database: massively expanding the structural coverage of protein-sequence space with high-accuracy models. *Nucleic Acids Res.* **50**, D439–D444 (2022).
- Velankar, S. *et al.* SIFTS: Structure Integration with Function, Taxonomy and Sequences resource. *Nucleic Acids Res.* **41**, D483–D489 (2013).
- Waman, V.P. *et al.* CATH v4.4: major expansion of CATH by experimental and predicted structural data. *Nucleic Acids Res.* **53**, D348–D356 (2025).
- Wang, W. *et al.* DPFunc: accurately predicting protein function via deep learning with domain-guided structure information. *Nat. Commun.* **16**, 70 (2025).
- Weatheritt, R.J., Sterne-Weiler, T. & Blencowe, B.J. The ribosome-engaged landscape of alternative splicing. *Nat. Struct. Mol. Biol.* **23**, 1117–1123 (2016).
- Yamada, T. *et al.* Significance of Tsukushi in lung cancer. *Lung Cancer* **131**, 104–111 (2019).
- Yao, S. *et al.* NetGO 2.0: improving large-scale protein function prediction with massive sequence, text, domain, family and network information. *Nucleic Acids Res.* **49**, W469–W475 (2021).
- You, R. *et al.* GOLabeler: improving sequence-based large-scale protein function prediction by learning to rank. *Bioinformatics* **34**, 2465–2473 (2018).
- You, R. *et al.* NetGO: improving large-scale protein function prediction with massive network information. *Nucleic Acids Res.* **47**, W379–W387 (2019).
- You, R., Yao, S., Mamitsuka, H. & Zhu, S. DeepGraphGO: graph neural network for large-scale, multispecies protein function prediction. *Bioinformatics* **37**, i262–i271 (2021).
- Yu, G. *et al.* Isoform function prediction based on bi-random walks on a heterogeneous network. *Bioinformatics* **36**, 303–310 (2020).
- Zhang, Y. *et al.* Alternative splicing and cancer: a systematic review. *Signal Transduct. Target. Ther.* **6**, 78 (2021).

- Zhao, C. *et al.* PANDA-3D: protein function prediction based on AlphaFold models. *NAR Genom. Bioinform.* **6**, lqae094 (2024).
- Huntley, R. P. *et al*. The GOA database: Gene Ontology annotation updates for 2015. *Nucleic Acids Res*. **43**, D1057–D1063 (2015).
- The Gene Ontology Consortium. The Gene Ontology knowledgebase in 2026. Nucleic Acids Res. 54, D1779–D1792 (2026).
- Cox, T.R. The matrix in cancer. *Nat. Rev. Cancer* **21**, 217–238 (2021).
